# The gut bacterial community of black soldier fly larvae is a reservoir of antibiotic resistance and virulence genes

**DOI:** 10.64898/2026.06.18.732884

**Authors:** Davis Roma, Conor JR Scott, Matteo Brilli, Giuseppina Sequino, Alessia Esposito, Francesca De Filippis, Gianluca Tettamanti, Morena Casartelli, Silvia Caccia

**Author notes:** Corresponding authors: Morena Casartelli, Department of Biosciences, University of Milan, via Celoria 26, 20133, Milan, Italy. email address, Silvia Caccia, Department of Biosciences, University of Milan, via Celoria 26, 20133, Milan, Italy. These authors have equally contributed to this work (Davis Roma, Conor JR Scott).

## Abstract

Antimicrobial resistance (AMR) is a serious threat to global health. Agricultural practices that have contributed greatly to AMR spread urgently require innovation to address this issue, and more broadly challenges of sustainability and environmental concern. The larvae of black soldier fly (BSFL), *Hermetia illucens*, are considered a promising resource for advancing sustainable and circular agri-food systems given their ability to bioconvert organic waste streams into protein-and lipid-rich biomass suitable for feed applications and the use of the rearing residues (i.e., frass) as organic fertilisers. However, despite their emerging industrial applications, the risks of antibiotic resistance spread through their use remain underexplored. To elucidate this aspect, the profiles of antibiotic resistance genes (ARGs) and virulence factors (VFs), and their occurrence on plasmids were predicted from the midgut bacterial community of BSFL. Shotgun metagenomics revealed candidate resistance genes for 26 classes of antibiotics, and virulence via 9 mechanisms (with mobility and biofilm formation as major ones), with taxa belonging to the Pseudomonadota phylum as the dominant contributors. Highly relevant to public health was the identification of genes encoding resistance to carbapenem class antibiotics in bacterial genomes and mobile plasmids. Reconstruction of metagenomes enabled more precise taxonomic resolution and revealed taxa harbouring multiple resistance and virulence genes, including a *Pseudomonas* species with 42 VFs and 7 ARGs. Notably, for the first time antibiotic resistant bacterial species were isolated from the gut microbiota of BSFL, validating and complementing the results obtained *in silico*. Together, this work represents a comprehensive profile of the BSFL midgut bacterial resistome, while also providing relevant context on virulence and mobility. Importantly, it emphasises the urgent need to adopt strategies to mitigate potential risks arising from the development of emerging technologies related to the use of insect-mediated bioconversion and derived products.

## 2. Introduction

The spread of antimicrobial resistance genes (ARGs) has become a crucial global health threat, menacing the efficacy of the treatment and prevention of infections. Roughly 5 million deaths were associated with bacterial antimicrobial resistance (AMR) in 2019, and this number is projected to increase up to 10 million by 2050 (Ho *et al*. 2025). Among other factors, agricultural practices and livestock production have greatly contributed to the spread of ARGs in the environment, with recent demands for a reduction in the use of antibiotics and the establishment of more sustainable processes (Wee *et al*. 2020; Ho *et al*. 2025). ARG dissemination occurs via horizontal gene transfer of ARG-encoding DNA either through conjugation-based transfer of ARG-carrying mobile genetic elements (Jian *et al*. 2021; Wang and Dagan 2024; Munshi *et al*. 2025), transformation through the uptake of extracellular DNA (Bender *et al*. 2022), or other bacteriophage and membrane vesicle-based mechanisms (Tao *et al*. 2022). Additionally, the horizontal transfer and dissemination of virulence factor (VF) genes through the same mechanisms is of growing concern as these VF genes might provide a selective advantage to ARG-carrying bacteria by improving growth, reproduction and resistance in stressful environments (Niu *et al*. 2013), while potentially enabling pathogenicity (Fang *et al*. 2024; Zhou *et al*. 2024).

The European Union’s concern about this problem is highlighted by the One Health Action Plan against AMR which aims to improve the surveillance of ARGs and reduce antibiotic use by linking human health to ecosystem, crop, and livestock conditions (European Commission 2017). However, as highlighted by the adoption of the circular economy action plan for sustainable growth by the European Union, the spread of AMR and pathogens are not the sole challenges facing the agricultural industry (European Commission 2020). The urgent need to innovate practices in the food system, while simultaneously improving the circularity of waste streams to reduce the overall waste generation, must also consider the AMR risks associated with any innovation. One such innovation is the development of insect-based biorefineries to support circular economy processes (Drewery *et al*. 2022; Mannaa *et al*. 2023; Kim *et al*. 2025).

Insects have emerged as agents in the renovation of agricultural practices and livestock production towards sustainability (Mannaa *et al*. 2023; Bruno *et al*. 2025b; Kim *et al*. 2025). Among them, the Black Soldier Fly (BSF) *Hermetia illucens* is increasingly recognised as a model for a wide range of biotechnological applications (Tettamanti and Bruno 2024; Bruno *et al*. 2025b; Caccia *et al*. 2025). Principally, scientific research into BSF larvae (BSFL) has focused on their capability to efficiently bioconvert organic wastes and byproducts into biomass rich in proteins and lipids that can be used in the feed and food sectors (Bonelli *et al*. 2020; Drewery *et al*. 2022; Kim *et al*. 2025). Proteins and lipids from BSFL are also suitable for bioplastic and biodiesel production, respectively (Setti *et al*. 2020; Jung *et al*. 2022; Bruno *et al*. 2025b). Additionally, the frass (i.e., the residues at the end of insect rearing) is suitable as fertiliser (Jenkins *et al*. 2023; Lomonaco *et al*. 2024). Therefore, bioconversion processes driven by BSFL represent a powerful means for creating circular-economy value chains in which waste is transformed into valuable resources.

The production of BSF biomass from waste streams is, however, strictly regulated in the European Union. For example, the use of BSF meals in the feed sector is restricted to insects reared on substrates of vegetable origin or specifically allowed materials of animal origin considered as microbiologically and chemically safe (European Commission 2017; Lähteenmäki-Uutela *et al*. 2021). In other countries, regulation of insect rearing, production processes, and applications may vary considerably (Lähteenmäki-Uutela *et al*. 2021).

The capability of BSFL to efficiently bioconvert a wide range of organic materials, including nutritionally poor substrates, is derived from their saprophagous feeding habits and the high functional plasticity of their gut and the associated microbiota (Bruno *et al*. 2019; Bonelli *et al*. 2020; De Filippis *et al*. 2023; 2025; Vandeweyer *et al*. 2023). Unraveling the role of the complex BSFL gut microbial community as reservoir of ARGs must be clarified to establish safe and informed BSFL-mediated processes, especially in low-and middle-income countries, where the AMR-related morbidity or mortality are particularly severe and regulation on insect rearing may be lenient (Lähteenmäki-Uutela *et al*. 2021; Ho *et al*. 2025). Despite the growing relevance of the topic, only a few studies have investigated the diversity and distribution of ARGs in the BSFL microbiome (Zhao *et al*. 2023a; 2023b; Bohm *et al*. 2024; Rong *et al*. 2024; Schokker *et al*. 2025). These studies, based only on metagenomic analyses, provided important insight into the fate of ARGs across different waste streams. Moreover, they showed diet-dependent changes in both microbial community structure and resistome profiles with AMR bacteria still present in the final products of bioconversion processes which may pose a risk in their use (Schokker *et al*. 2025). However, these studies did not fully characterise the co-occurrence of VFs and ARGs in microorganisms of the BSFL midgut community. Most importantly, *in silico* predictions were not supported by *in vivo* microbiological evidence. Therefore, effectively profiling the midgut resistome and virulome of BSFL is not only valuable to increase basic knowledge on this topic, but will also provide an assessment of the real environmental and public health risks associated with the biotechnological applications based on BSFL.

Herein, remaining knowledge gaps were addressed by establishing a comprehensive platform of knowledge based on metagenomic analysis of the ARGs and VFs, and the mobile plasmids encoded within the BSFL midgut. The investigation of antibiotic resistance was extended by the isolation and species-level identification of antibiotic-resistant strains from the BSFL midgut. Through these analyses, we present a thorough profile of the BSFL midgut resistome together with virulome and plasmid profiles, to provide a valuable resource for the BSFL research community for the informed use of these larvae in agricultural processes.

## 3. Materials and Methods

### 3.1 Shotgun metagenomics of the BSFL gut microbiota

All *in silico* analyses were performed using shotgun metagenomic data of the midgut community of BSFL reared on standard diet (STD: 50 % (w/w) wheat bran; 20 % (w/w) alfalfa meal; 30 % (w/w) corn meal, mixed with water in 1:1 (w/w) ratio), obtained previously in De Filippis *et al*. (2023). Peritrophic matrix enclosed midguts were pooled into six independent samples of 15-30 midguts for DNA extraction, purification and library preparation with Illumina NovaSeq paired-end read sequencing. Host reads contamination was removed mapping reads to the *H. illucens* genome (NCBI Accession Number: PRJEB37575) by using BMTagger (NCBI 2011) as reported previously (De Filippis *et al*. 2023). The raw sequence reads generated previously were deposited in the Sequence Read Archive (SRA) of the NCBI under accession number PRJNA864640.

### 3.2 Coassembly and read mapping

Shotgun metagenomic reads were quality-filtered using PRINSEQ 0.20.4 (Schmieder and Edwards 2011), to keep only reads with a Phred score ≥ 20 and length ≥ 75 bp previously (De Filippis *et al*. 2023). FastQC 0.11.5 was used to assess read quality prior to coassembly (Andrews 2010) and quality control results were compiled with MultiQC 1.16 into an interactive report for confirmation of sufficient read quality in all samples (Ewels *et al*. 2016) (**Supplementary File 1**). Coassembly of contigs using merged forward and reverse reads from all six biological replicates was performed using MEGAHIT 1.2.9 with a minimum contig length of 1,000 bp, and the ‘meta-large’ preset enabled for high biodiversity samples (Li *et al*. 2015). The resulting coassembly consisted of 112,981 contigs with a total length of 535,307,860 bp, with contigs ranging from 1,000 bp to 1,615,041 bp in size, and with an N50 value of 12,625 bp. The coassembly was indexed and reads for each sample were mapped to the coassembly with Bowtie2 2.3.4.1 (Langmead and Salzberg 2012), with success rates across samples ranging from 89.0 to 91.7 %. Samtools 1.7 was used to sort and index the alignment file for downstream analysis (Li *et al*. 2009). The ‘jgi_summarize_bam_contig_depths’ function of MetaBAT2 2.2.17 was used to generate average read depth values and standard deviation of read depth values for coassembly contigs independently for each sample (Kang *et al*. 2019).

### 3.3 Contig taxonomic assignment

For contig-level taxonomic assignment, the coassembly was filtered to retain 112,892 contigs with a minimum contig length of 1,000 bp and a minimum average read depth of 2 based on MetaBAT2 2.2.17 depth summary results. Prodigal 2.6.3 was used to predict 579,379 open reading frames (ORFs) representing 475,501,254 bp out of the 535,209,308 bp of the filtered metagenomic contigs (Hyatt *et al*. 2010). The resulting predicted amino acid sequences were then used for taxonomic classification of contigs with the Contig Annotation Tool (CAT) 6.0.1 due to the higher precision of CAT on bacterial benchmarks compared to other tools resulting from the ORF-based strategy that made it a suitable choice for the classification of longer contigs compared to shorter reads (Mirdita *et al*. 2021). CAT leverages DIAMOND 2.1.12 for protein sequence alignment of ORFs against a pre-constructed NCBI non-redundant protein database obtained from the CAT repository (Buchfink *et al*. 2021; von Meijenfeldt *et al*. 2019). For each predicted ORF, DIAMOND hits with a bitscore ≥ 50 % of the top-scoring hit were retained, and a weighted consensus across these hits was used to assign taxonomy to the contig. Of the 112,892 contigs retained after filtering, 93046 (82.4 %) were assigned to the superkingdom bacteria by CAT. The relative abundance of bacterial contigs was similar across samples with an average of 89.7 % and a standard deviation of ± 1.19 %. Classification success and unique classifications at each taxonomic rank can be seen in (**Supplementary table 7.1**). For additional context, Kraken2 2.1.6 was also used to predict taxonomic IDs of contigs using the pre-built July 2025 standard Kraken2 database and with a default confidence threshold of 0.2 (Wood *et al*. 2019). TaxonKit 0.20.0 was used to obtain full taxonomic lineages for Kraken2 assigned taxonomic IDs in combination with the NCBI Taxonomy database (Sayers *et al*. 2022). Due to the previously reported increased false positive rate associated with the used of Kraken2 (Wood *et al*. 2019), Kraken2 was used solely to provide alternative taxonomic classifications for readers.

### 3.4 Contig binning and MAG taxonomic classification

For contig binning into metagenome-assembled genomes (MAGs), the coassembly was filtered to retain 37,454 contigs with a minimum length of 2,000 bp, a minimum average read depth of 5 based on MetaBAT2 2.2.17 depth summary results, and a maximum average coverage variance of 50. Binning of contigs to MAGs was performed using MetaBAT2 2.2.17 (Kang *et al*. 2019), achieving binning of 94.0 % of the filtered contigs representing 307,506,758 bp across 110 bins. The completeness and contamination percentages of MAGs were then estimated with CheckM 1.2.4 (Parks *et al*. 2015). Then, 48 ‘medium-quality MAGs’ were retained by filtering MAGs to a minimum completeness of 50 % and a maximum contamination of 5 %. Taxonomic assignment of MAGs was performed with GTDB-Tk 2.3.2, referencing the GTDB release r207 v2 (Chaumeil *et al*. 2020). The ‘classify_wf’ workflow was run with default parameters, skipping any dereplication or redundancy filtering with the ‘skip_ani_screen’ argument. MAGs were classified to species using FastANI 1.32 where average nucleotide identity (ANI) and alignment fraction (AF) to the closest reference genome met the GTDB thresholds of 95 % and 65 %, respectively (Jain *et al*. 2018). Otherwise, taxonomic classification was assigned based on the alignment of MAGs to marker genes in the GTDB-Tk alignment pipeline and placement into the bacterial reference tree using pplacer 1.1 (Matsen *et al*. 2010). GTDB-Tk taxonomic classification was successful for 47 MAGs.

### 3.5 ARG, VF, and plasmid sequence prediction

Contig fasta sequences were analysed using the ABRicate software 1.0.1 (Seemann 2025). The Comprehensive Antibiotic Resistance Database (CARD) (Alcock *et al*. 2023), the Virulence Factor Database (VFDB) (Zhou *et al*. 2024), and PlasmidFinder (Carattoli *et al*. 2014) were utilised to search for ARG, VF, and plasmid sequences, respectively. All databases used with ABRicate were updated in January 2025. For additional ARG sequence metadata, “CARD data” 4.0.1 was downloaded from CARD and metadata extracted for identified ARG sequences using the predicted gene name. Predicted VF sequences were classified to 14 base categories according to the VFDB database based on VF product descriptions and gene names (Liu *et al*. 2022). Unique plasmid accessions identified in the BSFL midgut microbiota were searched within the “ALL Plasmid Meta Data”, “ALL Plasmid Antimicrobial Resistance Gene Meta Data”, and “ALL Plasmid Virulent Factor Meta Data” databases available for download from the PlasmidScope database for predictions of plasmid mobility, ARG genes, and VF genes, respectively (Li *et al*. 2024).

### 3.6 Isolation of antibiotic-resistant bacteria

BSF eggs were collected from a colony established in 2015 at the University of Insubria (Varese, Italy) and maintained as reported in Bruno *et al*. (2019). The eggs were laid on a Petri dish (9 × 1.5 cm) with STD for fly larvae rearing (Hogsette 1992) and maintained in a humid chamber at 27 °C until hatching. Nipagin (methyl 4-hydroxybenzoate) was added to the diet to avoid mold growth. After emergence, BSF larvae were reared on this diet for 4 days and then placed in plastics boxes (16 × 16 × 9 cm and 12 × 12 × 8 cm) containing STD without nipagin. Actively feeding last instar larvae were collected, rinsed with tap water, and then dried and washed with 70 % (v/v) ethanol. For each sample, the midguts from five larvae were isolated in sterile PBS in a sterile Petri dish (5.5 × 1.3 cm) using tweezers and scissors previously washed with 70 % (v/v) ethanol (Bruno et al., 2025a). The midgut content enclosed in the peritrophic matrix was separated from the epithelium and collected in 1.5 mL tubes (Bruno *et al*. 2025a). Then, 10 µL of sterile PBS for each mg of midgut content was added and resulting suspensions were homogenised with a plastic pestle. Concomitantly with the sampling of BSFL, approximately 100 mg of the rearing diet were also collected in 1.5 mL tubes, with 3 µL of sterile PBS added for each mg of diet and the resulting suspensions were crushed with a plastic pestle and vortexed. Five 10-fold dilutions of midgut content and diet samples were prepared in 1.5 mL tubes using sterile PBS. Luria Bertani (LB: 10 g/L tryptone, 5 g/L yeast extract, 10 g/L NaCl, 10 g/L agar) plates were prepared adding 8 different antibiotics (100 µg/mL ampicillin, 10 µg/mL tetracycline, 20 µg/mL chloramphenicol, 100 µg/mL streptomycin, 50 µg/mL kanamycin, 10 µg/mL gentamicin, 10 µg/mL ciprofloxacin and 10 µg/mL vancomycin). Antibiotic concentrations were chosen as common working concentration for laboratory plate preparation. Similar concentrations were used in previous works on antimicrobial resistance bacteria from environmental samples (Armalytė *et al*. 2019; Perelomov *et al*. 2023). The last three 10-fold dilutions (1/1,000, 1/10,000, 1/100,000) of midgut content suspensions and diet sample suspensions were plated onto LB plates with antibiotics and were grown overnight at 37 °C. Resulting colonies were then picked and streaked onto another LB plate containing the corresponding antibiotic and grown overnight at 37 °C to isolate unique taxa. For each isolate plate, a single colony was picked with an inoculation loop and dispersed into a solution of 50 % (v/v) glycerol in liquid LB medium within cryovials which were subsequently stored at-80 °C. Samples of freshly prepared STD were also prepared in the same way and plated, however no growth was observed on antibiotic plates.

### 3.7 MALDI-TOF species identification of resistant isolates

Resistant isolates were streaked on fresh plates of LB agar and incubated at 37 °C for 24 hours. Identification was performed using a MALDI-TOF Sirius Biotyper (Matrix-Assisted Laser Desorption/Ionization Time-Of-Flight; Bruker, USA), according to the manufacturer’s standard protocol. Briefly, 1 µL of 70 % (v/v) formic acid solution in water and 1 µL of α-cyano-4-hydrocinnamic acid matrix solution (Bruker, USA) were added directly onto the colonies and allowed to air dry. Spectra were analyzed with Bruker Biotyper Compass HT RUO software 5.1.3 with the MBT Compass Library 12.0. Calibration was performed for each target plate using Bruker Bacterial Test Standard (BTS). To ensure confidence in the identification of bacterial taxa from samples, a log score of spectral similarity above the 1.7 threshold was considered.

### 3.8 Data analysis

Read depths for each contig were normalised to relative abundances based on the total read depth per sample. To investigate the bacterial community structure, contigs were filtered to those predicted as “Bacteria” by CAT and Kraken2 separately and the read depths re-normalised for these subsets of contigs to obtain ‘relative bacterial abundances’. All data analysis and visualisation was performed in RStudio 2025.09.0+387 “Cucumberleaf Sunflower” release built on R 4.5.1 (Posit team 2025). Phylum-level community structures for the entire bacterial community, the counts of VFs belonging to the VF product classes, and MAG phylum-level classification numbers were plotted with the ‘ggplot2’ package (Wickham 2016). Heatmaps of antibiotic resistance mechanism against drug class, phylum level-contributions to ARGs and VFs of bacterial contigs, and genus-level contributions of bacterial MAGs to ARGs and VFs were plotted with the ‘pheatmap’ package (Kolde 2025). Plasmid mobility and contributions to antibiotic resistance, and antibiotic resistance of isolates were visualised with alluvial plots using the ‘ggalluvial’ package (Brunson 2020). To visualise MAG-gene associations for ARG and VF genes, an edge-table was generated from links between MAGs and genes based on contig IDs of predicted genes and binned contigs. The ‘igraph’ package was used to generate a graph object distinguishing Taxon, ARG, and VF nodes based on size and colour, and edges were distinguished based on single taxon or multiple taxon connections (Csárdi *et al*. 2025). The network was plotted using the ‘ggraph’ package utilising a Fruchterman-Reingold layout (Pedersen 2024). Multi-panel figures of ggplot objects were combined with the ‘patchwork’ package (Pedersen 2025). Multiple pheatmap plots were combined with the ‘gridExtra’ package (Auguie 2017). Combinations of ggplot and pheatmap objects for figures were made by saving objects as individual image files and combining with the ‘magick’ package (Ooms 2025).

Interactive kingdom and phylum bacterial relative abundance plots were created using the ‘ggplot’ and ‘plotly’ packages (Wickham 2016; Sievert 2020). Interactive sankey diagrams of the bacterial relative abundance ordered hierarchically from Class to Order to Family taxonomic classifications were constructed using the ‘sankeyNetwork’ function of the ‘networkD3’ package, and were rendered with customisation using the ‘htmlwidgets’ package (Vaidyanathan *et al*. 2023; Allaire *et al*. 2025). Interactive heatmaps of genus-and species-level community structures across samples were visualised using the ‘heatmaply’ package (Galili *et al*. 2017). Interactive community figures for both CAT-and Kraken2-classified bacterial contig communities can be seen in **Supplementary File 2**. The interactive sunburst plot depicting the hierarchical structure of MAG taxonomic classifications was plotted using the ‘plotly’ package (Sievert 2020). MAG genus and species classifications by GTDB-Tk alongside FastANI and pplacer reference genome accessions were presented in table format using the ‘kableExtra’ package (Zhu 2024). The interactive MAG taxonomic classifications are visible in **Supplementary File 4**.

The submitted manuscript was generated in Word.docx format and the interactive figures were rendered in.html supplementary files from ‘rmarkdown’ using the ‘knitr’ and ‘bookdown’ packages (Xie 2014; 2016; Xie *et al*. 2020). Tables in the.docx manuscript were generated using the ‘flextable’ package (Gohel and Skintzos 2025).

## 4 Results

### 4.1 Bacterial community structure of the BSFL midgut microbiota

Shotgun metagenomic contigs used for the resistome and virulome characterization were first analysed to obtain a taxonomic overview of bacterial midgut community in larvae reared on the standard diet (**Figure 4.1**). The results at Phylum level were similar to previous assessments based on *16S rRNA* gene amplicon sequencing or marker-gene based assignment of shotgun metagenomic reads for larvae reared on the same diet (De Filippis *et al*. 2023).

**Figure 4.1:**
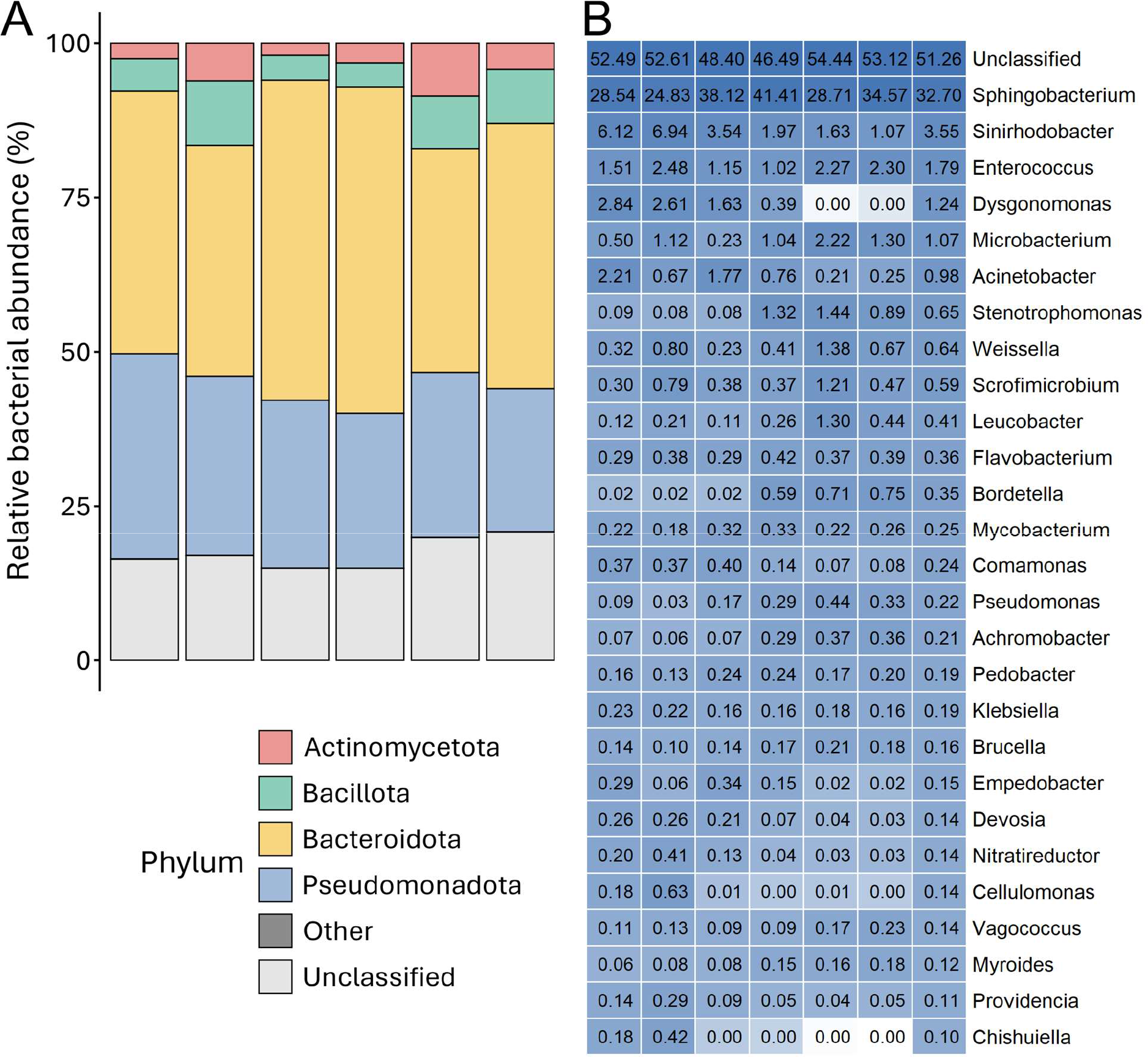
Bacterial community structure of the BSFL midgut. (**A**) The relative bacterial abundances of bacterial phyla for each sample. Phyla with relative bacterial abundances < 1 % were grouped into Other. (**B**) Genera with average relative bacterial abundances > 0.1 % across the six samples. Bacterial relative abundance values for genera are presented within cells for each sample, with genera ordered in descending mean relative bacterial abundance. Columns represent individual samples.

Bacteroidota was the most abundant phylum (43.9 % on average), while Pseudomonadota contributed on average to 27.5 % of the relative bacterial abundance (**Figure 4.1**, **Supplementary table 7.2**). Actinomycetota and Bacillota phyla constituted on average 4.47 % and 6.81 %, respectively. The remaining classified phyla together contributed only 0.0378 %.

The remaining 17.3 % of the relative bacterial abundance of the community structure was derived from contigs that were unclassified at phylum level by CAT. About 51.3 % of bacterial contigs were attributed to taxa unclassified at the genus level by CAT. Contigs unclassified at genus were distributed across 15 phyla including Actinomycetota, Bacillota, Bacteroidota, and Pseudomonadota. *Sphingobacterium* was observed with an average bacterial relative abundance of 32.7 % while *Sinirhodobacter* represented the 3.55 %. The community structure from kingdom to species taxonomic ranks can be explored interactively in more detail for CAT and Kraken2 predicted communities in **Supplementary File 2**.

### 4.2 Antibiotic resistance and virulence in the BSFL midgut microbiota

In total, 318 ARGs were predicted across all contigs, of which 212 genes were identified on 169 unique bacterial contigs. The remaining ARG-carrying contigs were not assigned taxonomically. The relative abundance of bacterial contigs carrying ARGs accounted for 0.376% of the total bacterial contig abundance. In comparison, 333 VFs were predicted across all contigs, of which 297 were identified on 198 bacterial contigs. Again, all other VF-carrying contigs were unassigned. The bacterial VF-carrying contigs accounted for 0.182 % of the total bacterial contig abundance. The entire database of all predicted ARGs and VFs alongside relative abundance across samples and contig taxonomic classifications can be found in **Supplementary File 3**.

In total, 784 incidences of resistance were predicted to be conferred to 26 classes of antibiotic compound by ARGs predicted on bacterial contigs, where each gene may confer an incidence of resistance to multiple antibiotic classes. (**Supplementary table 7.3**). The fluoroquinolone class of antibiotics had the highest number of incidences of resistance conferred against it with a total of 99 incidences of resistance. Antibiotic efflux was the most abundant encoded mechanism of resistance with 132 genes which conferred 506 incidences of resistance against 21 classes of antibiotics (**Figure 4.2A**). Smaller numbers were predicted to be conferred by any other mechanism. Additionally, 57 ARGs were unclassified in their mechanism of action and conferred 243 incidences of resistance against 22 classes of antibiotics.

**Figure 4.2:**
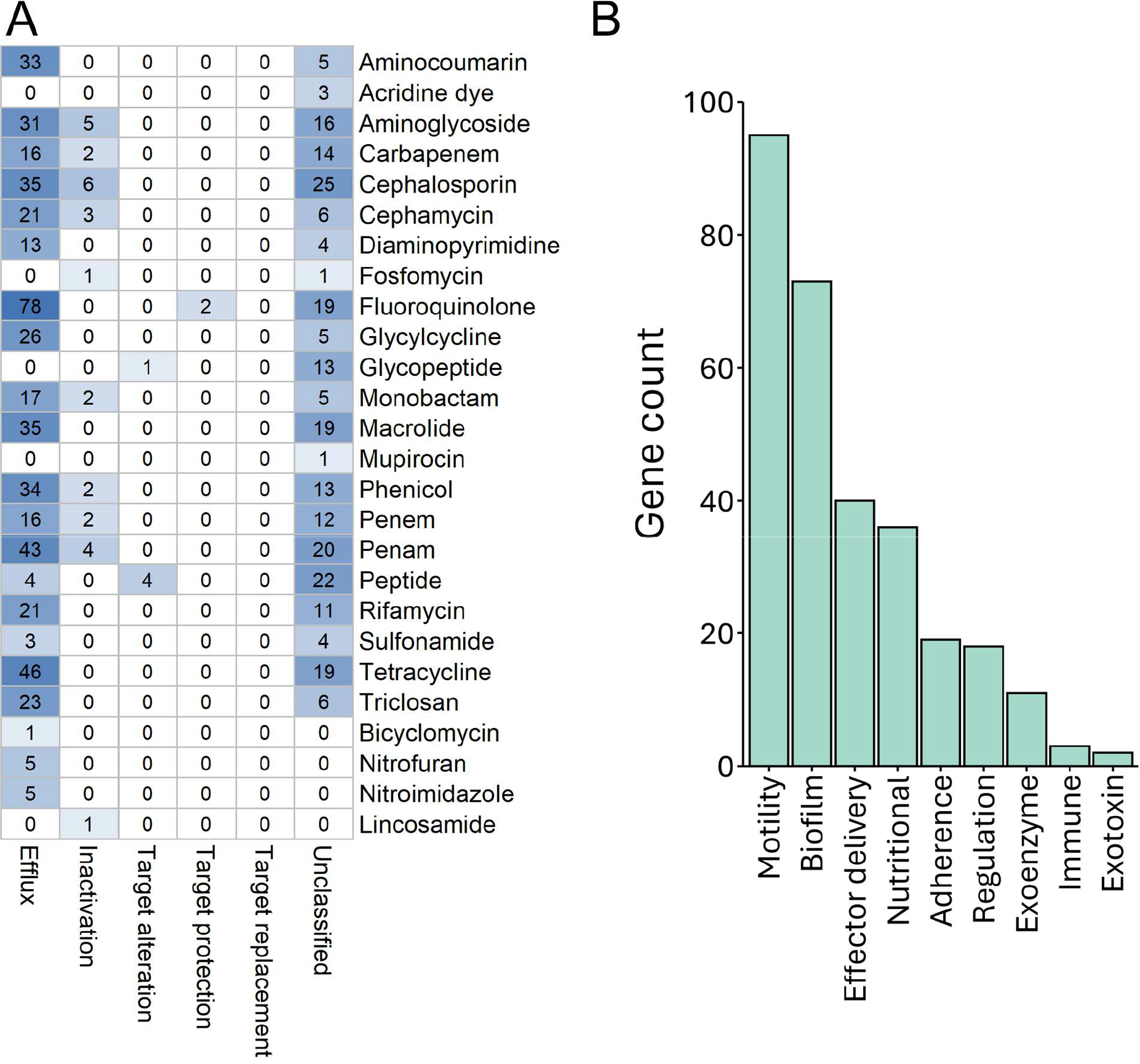
Antibiotic resistance and virulence genes in the BSFL bacterial midgut community. (**A**) The incidences of resistance predicted to be conferred to each of the antibiotic classes by the ARGs of each resistance mechanism for ARGs identified on bacterial contigs. (**B**) The number of predicted VF genes on bacterial contigs belonging to each VF class.

Regarding virulence, motility class VFs were the most common with 95 VF genes predicted for this class, followed by 73 for biofilm class VFs (**Figure 4.2B**). In total, nine unique modes of virulence of the 14 possible VF product classes in the VFDB database were encoded by genes in the bacterial community of the BSFL midgut.

Pseudomonadota phylum contributed the most to antibiotic resistance (**Figure 4.3**), with 422 total incidences of resistance conferred against 22 classes of antibiotics. However, 335 incidences of resistance against 24 classes resulted from ARG genes on bacterial contigs unclassified to the phylum level. Of the total number of contigs in the bacterial community that were classified to phylum, 54.5 % belonged to the Pseudomonadota phylum. This increased to 79.6 % when examining the bacterial ARG-carrying contigs classified to phylum.

**Figure 4.3:**
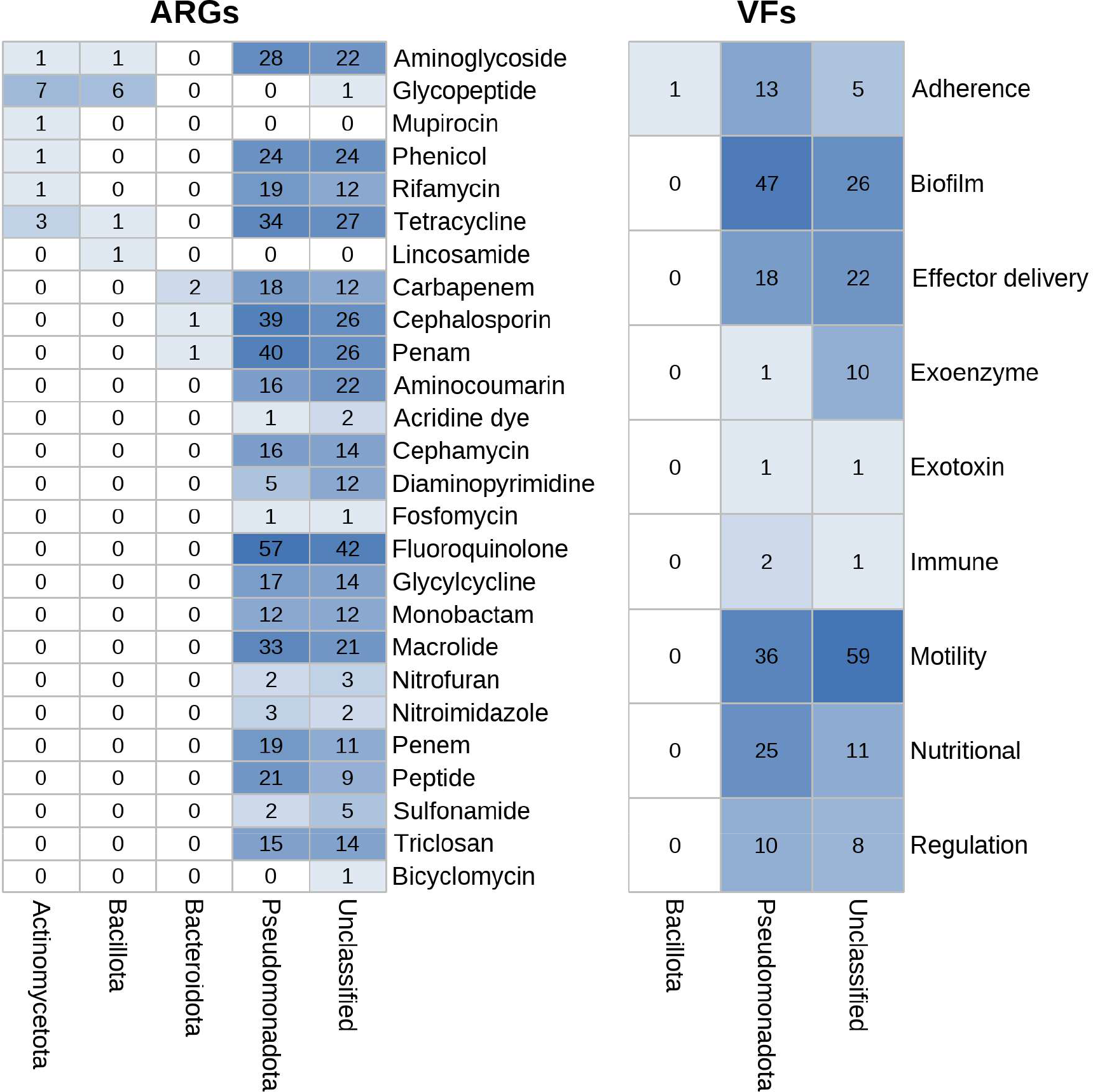
Resistant and virulent phyla in the BSFL bacterial midgut community. The number of incidences of resistance conferred by ARGs to the antibiotic classes and the counts of VF genes belonging to each product class, for genes identified on bacterial contigs classified to each bacterial phylum.

Pseudomonadota phylum contributed to all nine VF classes identified here via 153 predicted VFs on contigs assigned to this phylum (**Figure 4.3**). Only one VF was identified on a Bacillota assigned contig of the membrane class of VFs. The remaining 143 predicted VFs, spanning nine VF product classes, were identified on bacterial contigs unassigned to phylum. Interestingly, the 54.5 % of the number of bacterial phylum-classified contigs assigned to the Pseudomonadota phylum increased to 99 % when investigating the bacterial VF-carrying contigs classified to phylum.

### 4.3 Plasmids in the BSFL midgut microbiota

Abricate prediction of plasmids identified 53 contigs that encoded sequences with similarity to plasmid genes in the PlasmidFinder database, with 24 unique plasmid accessions identified overall. Only three contigs were predicted to encode both a plasmid and ARG sequences, and these ARG sequences were described as plasmid-mediated (**Supplementary table 7.4**). No contigs were predicted to encode both a plasmid gene and a VF. Searching the unique plasmid accession numbers in the PlasmidScope database of ARG metadata identified four of these unique plasmids recorded as encoding resistance, with one of them also identified from one of the three contigs encoding both a plasmid and an ARG sequence. Therefore, six unique plasmids were identified in total either from contigs also encoding an ARG sequence, or with accessions of ARG-carrying plasmids according to the PlasmidScope database.

Based on PlasmidScope metadata of the unique plasmid accessions identified in the contig coassembly by PlasmidFinder, three plasmids were predicted to be non-mobilisable, and three to be mobilisable, with two of them through means of conjugation. In total, 33 incidences of resistance were conferred against 13 unique classes of antibiotic compounds by the ARG genes predicted to occur on the plasmids identified in the BSFL midgut community (**Figure 4.4**). Furthermore, 25 of the incidences of resistance were encoded by the three mobilisable plasmids. Searching the accessions of the plasmids against the PlasmidScope VF metadata identified two unique plasmids predicted to encode VF sequences. Both plasmids (JN233704 and JN420336) were predicted to be mobile (**Supplementary table 7.5**) and identified as containing ARG sequences (**Figure 4.4**). The entire database of predicted plasmids alongside relative abundances across samples and contig taxonomic classifications can be seen in **Supplementary File 3**.

**Figure 4.4:**
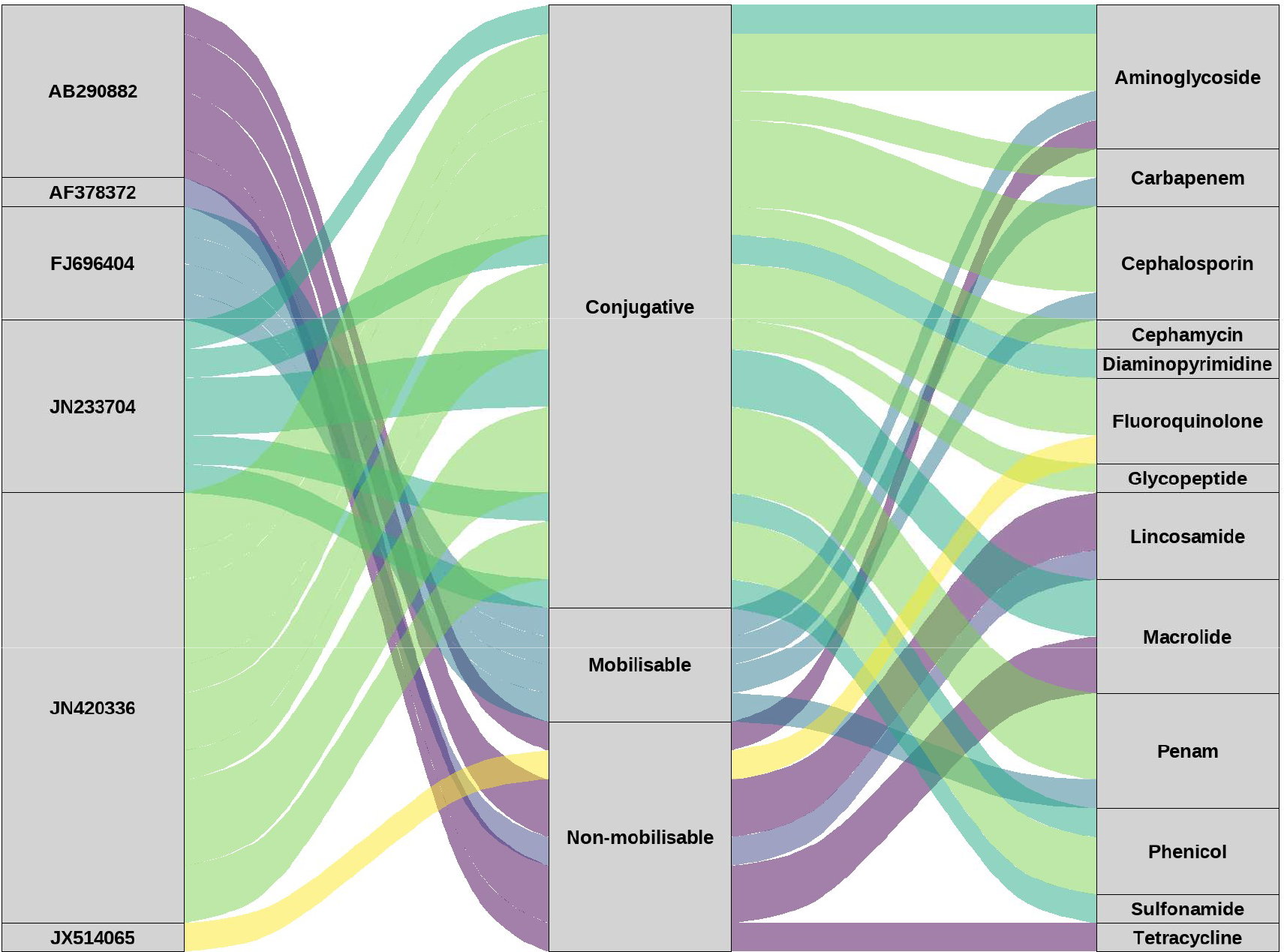
Plasmid-encoded resistance in the BSFL midgut community. The predicted mobility and the number of incidences of resistance conferred against antibiotic classes encoded by ARGs for unique plasmids identified in the BSFL midgut community.

### 4.4 Reconstruction of bacterial MAGs in the BSFL gut community

Fortyseven MAGs were identified at least to the genus level, and 26 were further identified to the species level. Most MAGs were assigned to the Pseudomonadota phylum (21), followed by Actinobacteriota (14), Bacillota (6), Bacteroidota (5), and one MAG was assigned to the phylum Bdellovibrionota (**Supplementary figure 7.1**). The full MAG taxonomic classifications, alongside genome accessions, can be seen in (**Supplementary File 4**).

Pseudomonadota phylum classified MAGs (**Figure 4.5**), contributed 184 of the 197 incidences of resistance conferred to 20 of the 21 classes of antibiotics from ARGs on contigs binned into MAGs. Within the Pseudomonadota phylum, the *Phytobacter* genus contributed 53 incidences of resistance against 17 classes,the *Pseudomonas* genus contributed 50 incidences of resistance against 17 classes, and the *Kosakonia* genus contributed 36 incidences of resistance against 16 classes.

**Figure 4.5:**
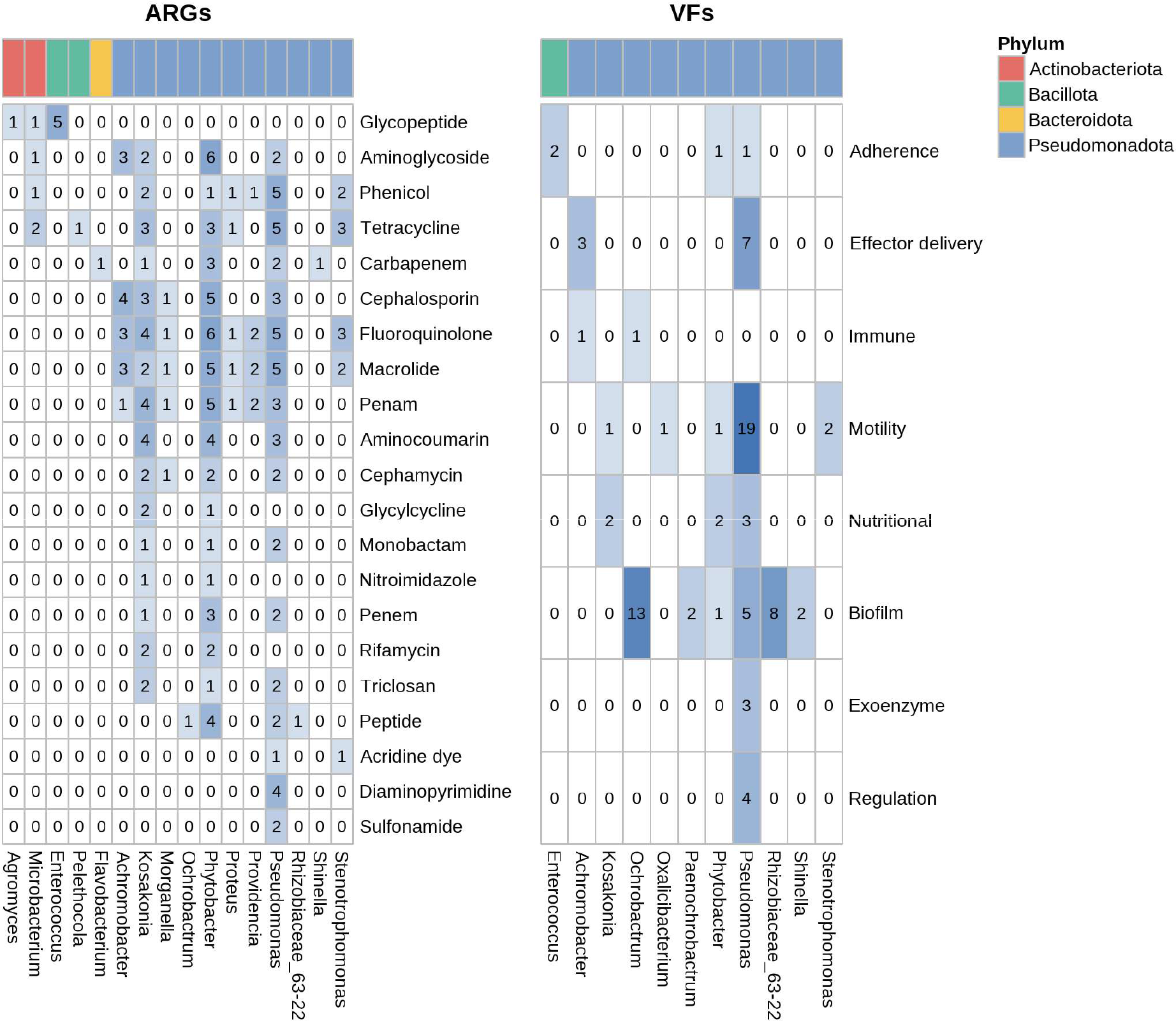
MAG contributions to antibiotic resistance and virulence. The number of incidences of resistance conferred to the antibiotic classes for ARGs, and the number of VF genes assigned to each VF product class for genes identified on contigs binned into MAGs. The colour of the label at the top of each column indicates the phylum to which the genus belongs.

Pseudomonadota phylum classified MAGs (**Figure 4.5**), contributed 83 of the 85 total VFs across all eight of the VF product classes encoded by these genes on contigs binned into MAGs. Within the Pseudomonadota phylum, the *Pseudomonas* genus contributed 42 VFs across seven classes, and the *Ochrobactrum* genus contributed 14 VFs across two classes. Of the 47 MAGs, nine were assembled from both ARG-and VF-encoding contigs and eight of these ARG-and VF-encoding MAGs were classified to the Pseudomonadota phylum, with only one to the Bacillota phylum.

The MAG of one *Pseudomonas* genus taxon consisted of contigs encoding in total 42 VFs and 7 ARGs (**Figure 4.6**).

**Figure 4.6:**
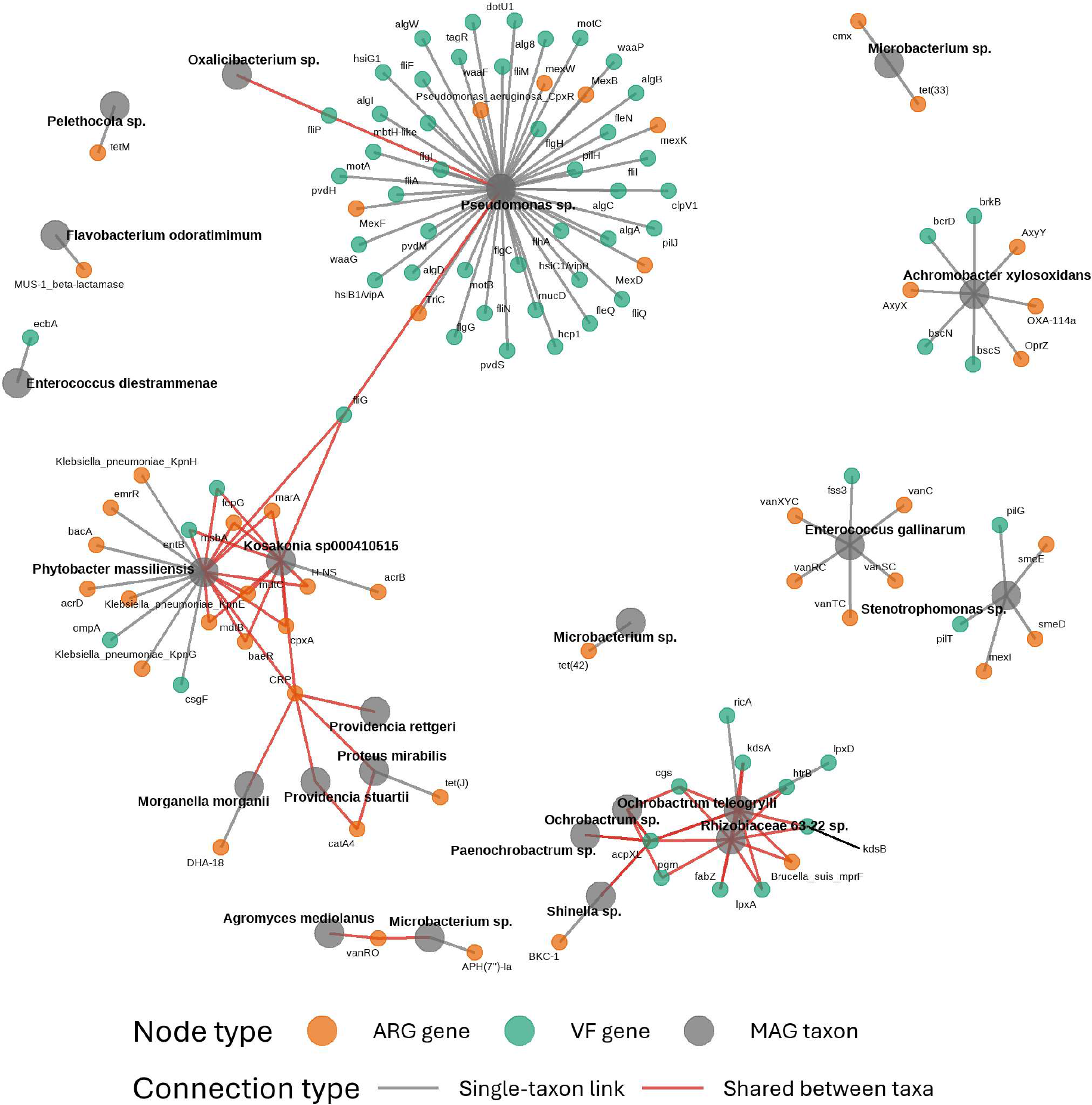
ARG and VF encoding MAGs. The ARG and VF genes encoded by contigs binned into taxonomically assigned MAGs. Unique taxa are represented by dark grey large nodes with ARG nodes in orange, and VF nodes in green. Grey links represent genes identified in the MAGs of a single taxon, red links represent genes identified in the MAGs of more than one taxon.

Only two of these genes, both VFs, were shared with other taxa. The *Kosakonia sp000410515* and *Phytobacter massiliensis* taxa, also belonging to the Pseudomonadota phylum, each had 12 and 19 genes, respectively. Although, 11 of these genes were shared. *Achromobacter xylosoxidans*, *Enterococcus gallinarum*, and a *Stenotrophomonas* species all had both ARG and VF sequences but shared none of their encoded genes with other MAGs. *Ochrobactrum teleogrylli*, a *Rhizobiaceae_63-22* species, and a *Shinella* species all encoded both ARG and VF sequences. An interactive version of this network can be seen in **Supplementary File 5**.

### 4.5 Isolation of antibiotic-resistant bacteria from the BSFL midgut

In total, nine taxa were identified from plating BSFL midgut lumen contents onto eight different antibiotic agar plates (**Figure 4.7A**). *Alcaligenes faecalis*, *Myroides injenensis*, and *Pseudomonas aeruginosa* species were all identified on more than one type of antibiotic-containing plate. *Bordetella trematum*, *Klebsiella pneumoniae*, *Leucobacter holotrichiae*, *Morganella morganii*, *Ochrobactrum intermedium*, and *Stenotrophomonas maltophilia* species were all identified on a single type of antibiotic plate investigated here. *A. faecalis*, *M. injenensis*, and *P. aeruginosa* species were all identified additionally in the rearing diet during larval growth (**Figure 4.7B**).

**Figure 4.7:**
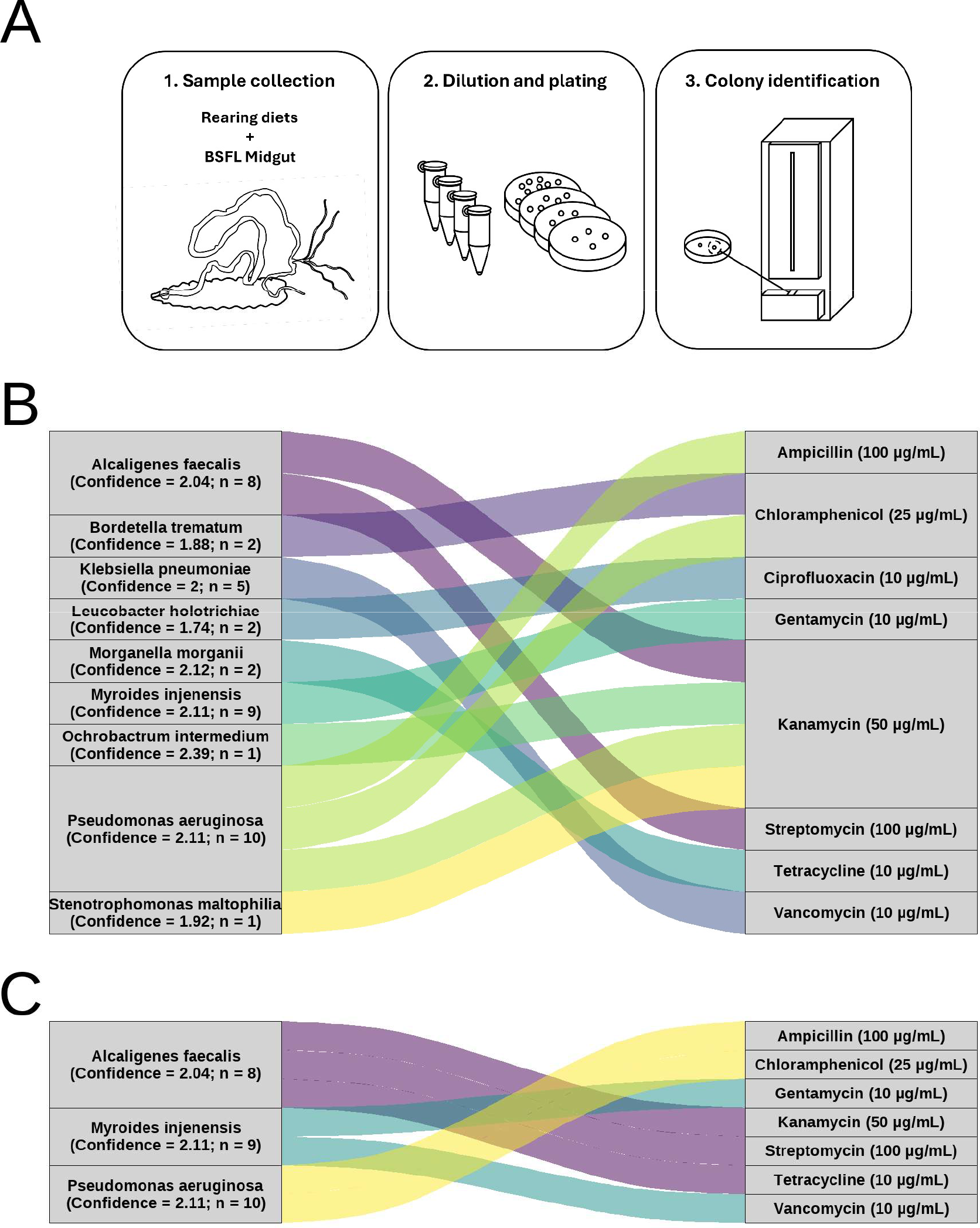
Resistant bacterial isolates of the BSFL bacterial midgut community. (**A**) Diagram of resistant midgut bacteria isolation. Midgut lumen content (**B**) or rearing diet during larval growth (**C**) was prepared and plated onto LB-agar plates with different antibiotics and incubated at 37 °C for 24h. Classification at species level was obtained using MALDI-TOF ID. The average log score of spectral similarity and the number of identifications were presented alongside identifications.

For the *M. morganii* isolate a potentially corresponding MAG was reconstructed (**Figure 4.5**). Contigs were identified classified to *B. trematum*, *S. maltophilia*, *K. pneumoniae*, *A. faecalis*, *M. morganii*, and *P. aeruginosa* species according to CAT taxonomic classification. Although contigs classified to *O. intermedium*, *M. injenensis*, and *L. holotrichiae* species were not identified, contigs classified to these genera were identified.

## 5 Discussion

Through the metagenomic analysis of the BSFL midgut microbiota performed in this study, it was possible to detect resistance genes for over one third of the 64 unique antibiotic classes reported in CARD, the most complete peer-reviewed database on antibiotic resistance (Alcock *et al*. 2023). The most abundant form of antibiotic resistance encoded in the bacterial community was mediated by efflux pumps. This was unsurprising due to the prevalence of this resistance strategy employed by a wide range of bacteria, and due to the lack of chemical or structural specificity with which efflux pumps export diverse antibiotics (Gaurav *et al*. 2023). Indeed, several of these efflux pumps might exert their activity against antibiotics as secondary targets and be involved in other functions. In the BSFL midgut microbiome, incidences of resistance to the fluoroquinolone class of antibiotics were the most common, reflecting the long-term use of these antibiotics beginning in the early 1960s (Kherroubi *et al*. 2024). Penam, including penicillins, and cephalosporins were the next most abundant resistance classes. Together these antibiotics constitute to 55 % of antibiotic consumption (Wang *et al*. 2025). Over 50 incidences of resistance were conferred to the tetracycline class, again a reflection of their long-term use in the past 60 years, and cause concern due to their wide target activity against both Gram-positive and-negative pathogens (Grossman 2016). The glycopeptide class of antibiotics, considered as “last resort” antibiotics (Selim 2022), had 14 incidences of resistance. In only 3 of the 26 classes a single incidence of resistance was observed. Overall, bacterial contigs predicted to encode sequences for ARGs were found to contribute only slightly to the community structure. The low relative abundances of ARGs identified in this study was similar to the low abundances of ARGs observed previously in BSFL (Rong *et al*. 2024; Schokker *et al*. 2025), This is unsurprising, as in the absence of selection ARGs could represent a metabolic burden for bacteria (Martínez and Rojo 2011). Similarly low numbers of ARGs have also been found in other environmental ARG reservoirs (Wang *et al*. 2020; Zhu *et al*. 2025), which still represents a serious concern.

A comprehensive profile of VFs encoded in the BSFL midgut microbiota was documented here. This not only provided an understanding of the potential pathogenicity associated with the midgut bacterial community, but may also be useful in combating antibiotic resistance, as it has been suggested that resistant pathogens could be disarmed through suppression of their virulence genes (Zhou *et al*. 2024). Investigating virulence identified 297 VF genes, with motility as the most abundant class of VF genes identified in the microbiota likely due to the large numbers of genes required to construct the highly complex structures enabling bacterial movement (Kao *et al*. 2014).

The second most abundant VF gene class in the midgut community was the biofilm formation class. Although biofilm formation is not directly involved in causing disease, it may allow pathogenic or resistant bacteria to persist in the environment, thus increasing the risk of ARG spread (Niu *et al*. 2013; Bai *et al*. 2021). Other VFs more directly related to pathogenicity such as effector delivery, exotoxins, and exoenzymes, were also identified in the BSFL midgut microbiome. Taxa harboring these genes alongside those for antibiotic resistance could represent a significant public health risk, and as discussed before, the dissemination of DNA encoding these traits could also be dangerous (Lichev *et al*. 2019). Similarly to the predicted ARGs, the Pseudomonadota phylum was the dominant taxonomic reservoir. Pseudomonadota is indeed a diverse phylum containing a range of pathogenic and antibiotic-resistant families (Leão *et al*. 2023). Furthermore, the dominance of this phylum in the ARG-encoding community of the BSFL midgut microbiota has been highlighted previously (Rong *et al*. 2024; Schokker *et al*. 2025). Several contigs containing typical plasmid sequences were identified in the BSFL midgut community, and in particular three mobile plasmids encoded antibiotic resistance. Plasmid JN420336 was predicted to encode resistance against the penam class of antibiotics and the widely used cephalosporins (Wang *et al*. 2025), while also encoding a VF for adherence and therefore with the potential to form films that increase the likelihood of the transfer of this conjugatively mobile plasmid (Dadeh Amirfard *et al*. 2024; Stalder and Top 2016). Plasmids JN420336 and JX514065, known vehicles for carbapenemase dissemination (Villa *et al*. 2012; Dolejska *et al*. 2013), were also reported here. The rise of carbapenemase resistant members of *Enterobacteriaceae* family is considered a serious concern and are therefore highly relevant for public health (Tuhamize and Bazira 2024). As discussed earlier however, plasmids are not the sole distributors of genes horizontally. Although the risk of horizontal gene transfer is hard to assess for genomic DNA which may become extracellular (Bender *et al*. 2022), transformation with extracellular DNA has contributed to the spread of both resistance and virulence (Lichev *et al*. 2019). Indeed, in combination with the capacity for biofilm formation in the BSFL midgut bacterial community, the likelihood of cell-cell contacts and therefore the spread of these ARG-encoding plasmids between community members and the environmental microbiome may even be increased (Dadeh Amirfard *et al*. 2024; Stalder and Top 2016).

Members of the BSFL midgut bacterial community with tools to cause disease and resist treatment already pose a risk if introduced into food systems. MAG reconstruction revealed a *Pseudomonas* taxon which encoded seven ARGs providing resistance theoretically to 17 classes of antibiotics, alongside 42 VFs mainly of the motility class but also of the effector delivery, nutritional modification, biofilm development, exoenzyme production, adherence, and modulation of transcriptional regulation classes of VFs. The disease-causing abilities of many *Pseudomonas* species are well documented, lying within the top six leading causes (73 %) of AMR-associated deaths in 2019 (Girard *et al*. 2021; Ho *et al*. 2025), but this was not the only taxa identified encoding both ARG and VF genes. *Kosakonia sp000410515* from the occasionally human-infecting genus *Kosakonia* (Chen *et al*. 2023; Merlino *et al*. 2025), and its phylogenetic neighbour *P. massiliensis* share 11 ARG and VF genes (Ma *et al*. 2020). *P. massiliensis* was part of a cluster of shared genes comprising other *Enterobacteraceae* species (i.e., *M. morganii*, *Providencia rettgeri* and *Providencia stuartii*). Some of the ARGs found in *P. massiliensis* encoded resistance against carbapenem and “last resort” antibiotics such as colistin. The spread of carbapenem resistant genes in the *Enterobacteriaceae* family is of high relevance for public health, as these antibiotics are the last line of defence for many serious infections (Tuhamize and Bazira 2024). These results could suggest that lesser-known members of the Enterobacteriaceae family such as *Phytobacter* can be important reservoirs of ARGs and their spread should be monitored, especially in new and emerging contexts like insect rearing facilities. *O. teleogrylli*, *A. xylosoxidans*, *E. gallinarum*, a species of the *Rhizobiaceae* genus, a species of the *Shinella* genus, and a species of the *Stenotrophomonas* genus all encoded both ARG and VF genes. Members of some of these taxa were already reported in clinical samples or healthcare-associated infections (Ryan *et al*. 2009; Monticelli *et al*. 2018; Isler *et al*. 2020; Ryan and Pembroke 2020), highlighting the need for careful monitoring of the microbiological safety of BSFL derivatives used in agriculture and food system processes.

To validate the bioinformatic analyses of metagenomic data, antibiotic-resistant bacterial species were isolated from the BSFL midgut and from the rearing diet using a culture-based approach with a selection of antibiotics. Nine unique bacterial isolates were identified at species level through MALDI-TOF across the eight antibiotics tested. Contigs belonging to all nine genera were identified in the shotgun metagenomic data. Three resistant species, i.e., *A. faecalis*, *M. injenensis*, and *P. aeruginosa* were also found in the frass. The transfer of genes through horizontal gene transfer has been reported for both *A. faecalis* (Lang *et al*. 2022) and *P. aeruginosa* (Karampatakis *et al*. 2025).

Overall, the results suggested that the microbiome of BSFL comprises several potentially pathogenic bacterial species carrying ARGs and VFs in their genomes, with microbiological evidence of the spread of some of these taxa to the rearing diet. This calls attention to the safety of workers in BSF facilities but should also demand the careful monitoring of BSFL-derived products. Moreover, despite the reduction of ARGs observed in some studies on diets with high prevalences of antimicrobial resistant bacteria (Zhao *et al*. 2023a; Zhao *et al*. 2023b), these genes are present during and at the end of the bioconversion practices. As suggested by the same authors, it will be important to establish downstream processes to reduce the risk of their spread in the environment, preventing the uptake of ARGs by the environmental microbiome or their introduction in the food chain.

BSFL remains an important biotechnological tool for the development of innovative processes that can offer innovation to agricultural and industrial processes. However, the associated risks should encourage careful monitoring and the adoption of procedures to mitigate ARG and VF dissemination into the environmental microbiome. Treatment strategies similar to those applied to manure to reduce ARG abundance could be adopted for BSFL frass (Zalewska *et al*. 2023), or phage-based approaches could be used to target specific high-risk bacterial taxa in BSFL-derived products (Liao *et al*. 2025). Overall, this study represents an important contribution to the BSFL research community, improving our understanding of the complex microbiome, and the potential health risks associated with its dissemination into food systems and the environment.

## 7 Supplementary material

### 7.1 Supplementary files

1. **Shotgun metagenomic read quality report.** MultiQC report of fastqc results of quality control checks for shotgun metagenomic reads files after trimming and filtering with PRINSEQ 0.20.4 for Phred score ≥ 20 and read length ≥ 75 bp.
2. **Interactive taxonomic community structures.** Interactive document with multiple interactive figures allowing all taxonomic levels from kingdom to species to be explored for the CAT and Kraken2 predicted the BSFL midgut microbiota bacterial community.
3. **ARG, VF, and Plasmid database.** Database of all predicted ARGs, VFs, and plasmids in the BSFL midgut microbial community coassembly from the CARD, VFDB, and PlasmidFinder databases, respectively.
4. **Interactive MAG taxonomic classification.** The GTDB-Tk taxonomic classification of the MAGs of the BSFL midgut microbiota in an interactive hierarchical sunburst plot alongside genus and species classifications with matching reference genome accessions identified with FastANI or pplacer approaches.
5. **Interactive MAG network plot of ARG and VF genes.** Interactive network plot depicting the connections between taxon nodes and encoded ARGs and VFs within MAGs of the BSFL midgut microbiota.
6. **R Markdown manuscript file.** The R markdown file used to generate this manuscript containing all text and the code for analysis and plotting.

## 7.2 Supplementary tables

**Table 7.1:** Taxonomic assignments of bacterial contigs in the BSFL midgut community. The number of contigs assigned at each taxonomic level and the number of unique classifications within each taxonomic level for contigs assigned to the bacterial superkingdom.

| **Taxonomic level** | **Contigs assigned** | **Unique classifications** |
| --- | --- | --- |
| Kingdom | 78,006 | 3 |
| Phylum | 75,549 | 19 |
| Class | 65,330 | 32 |
| Order | 53,068 | 62 |
| Family | 41,637 | 130 |
| Genus | 31,057 | 339 |
| Species | 13,140 | 984 |

**Table 7.2:**
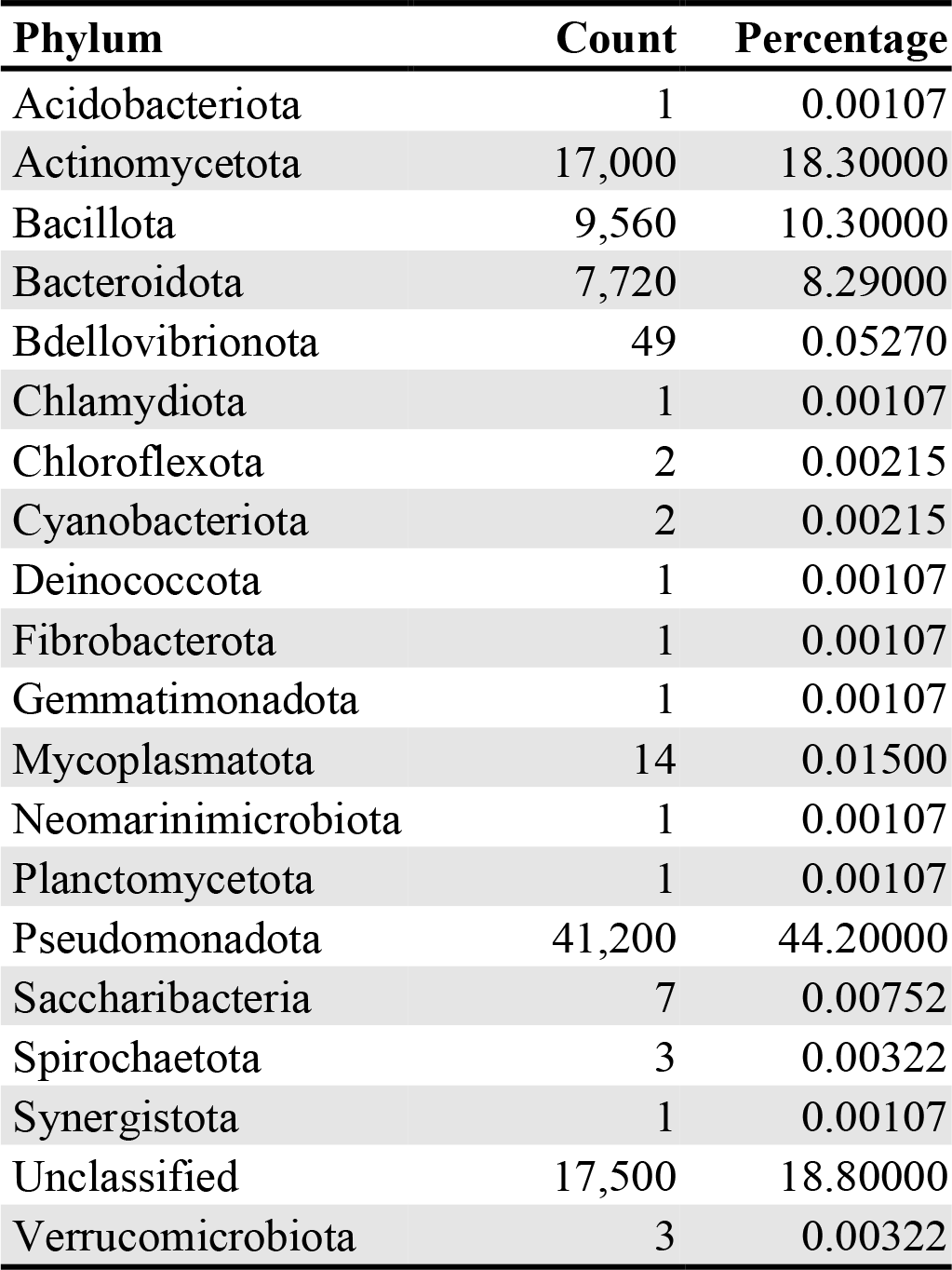
Taxonomic assignments to phyla for bacterial contigs in the BSFL midgut community. The counts and percentages of bacterial contigs assigned to each taxonomic phyla.

**Table 7.3:**
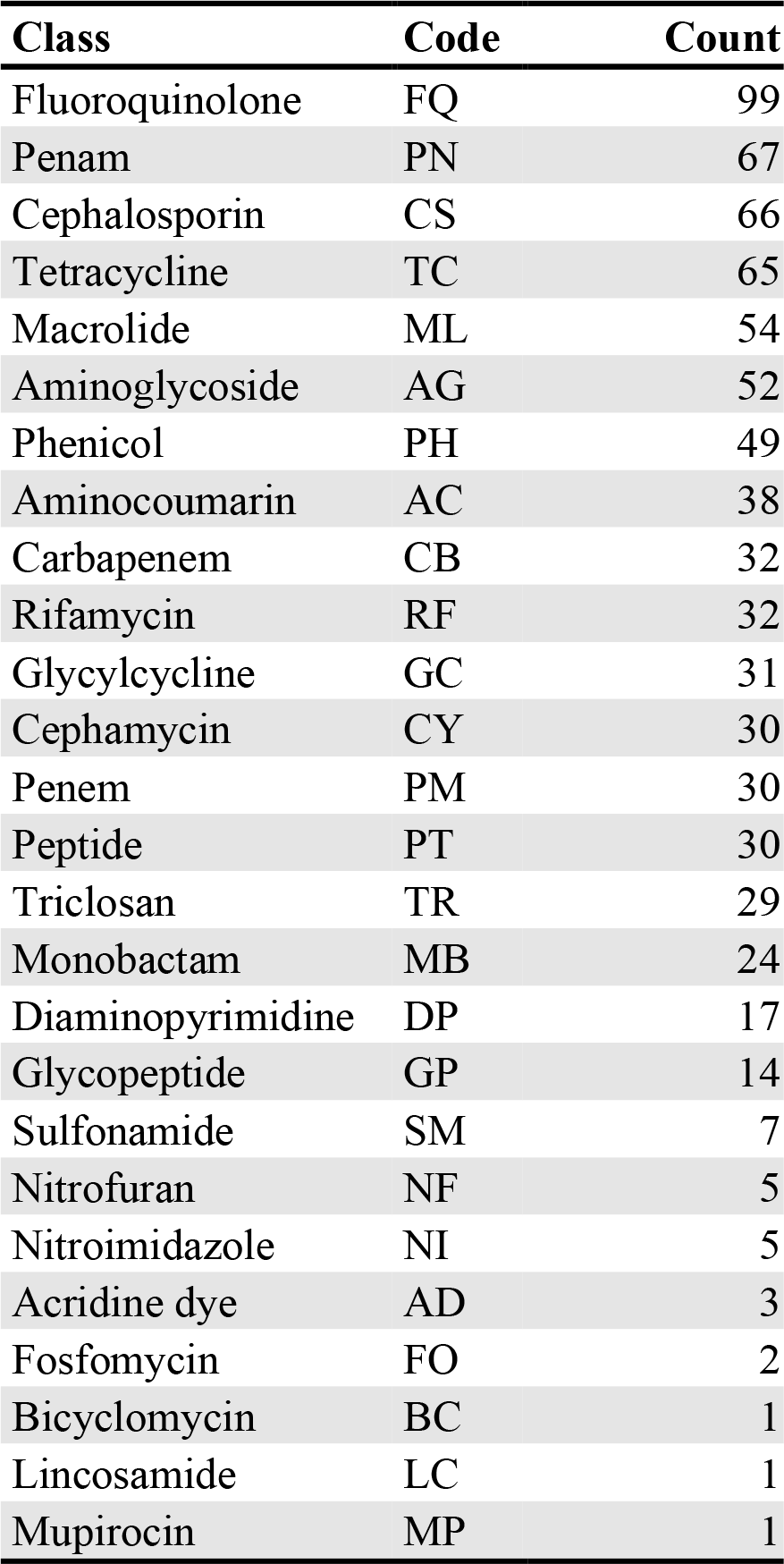
Encoded resistance to antibiotic classes in the BSFL midgut bacterial community. The number of incidences of resistance encoded against antibiotic classes by ARGs identified on bacterial contigs in the BSFL midgut community.

**Table 7.4:** Plasmid gene-and ARG-encoding contigs in the BSFL midgut community. Contigs of the coassembly predicted to encode both plasmid gene sequences and ARGs with identified plasmid accessions, ARG gene names, and CARD product description for the predicted ARG products.

| Contig | Plasmid accession | ARG name | CARD product description |
| --- | --- | --- | --- |
| k127_189896 | AF378372 | lnuA | lnuA is a plasmid-mediated nucleotidyltransferase found in *Staphylococcus chromogenes* |
| k127_399044 | JX514065 | QnrD2 | QnrD2 is a plasmid-mediated quinolone resistance protein found in *Salmonella enterica* |
| k127_113777 | AB290882 | ANT(4')-Ib | Kanamycin nucleotidyltransferase sequence from *Staphylococcus aureus* plasmid. Confers resistance to kanamycin neomycin and other aminoglycosides |

**Table 7.5:** VF-encoding plasmids of the BSFL midgut community. The accessions of plasmids identified in the coassembly with encoding VF sequences alongside the plasmid predicted mobility, predicted VF ID, the VF gene name, the VF category, and the product description.

| Accession | Mobility | ID | Name | Category | Product |
| --- | --- | --- | --- | --- | --- |
| JN420336 | Conjugative | VF015<br>9 | Hsp60 | Adherence | Hsp60, 60K heat shock protein HtpB |
| JN233704 | Conjugative | VF023<br>7 | Ibes | Invasion | Cu(+)/Ag(+) efflux RND transporter outer membrane channel CusC |

## 7.3 Supplementary figures

**Figure 7.1:**
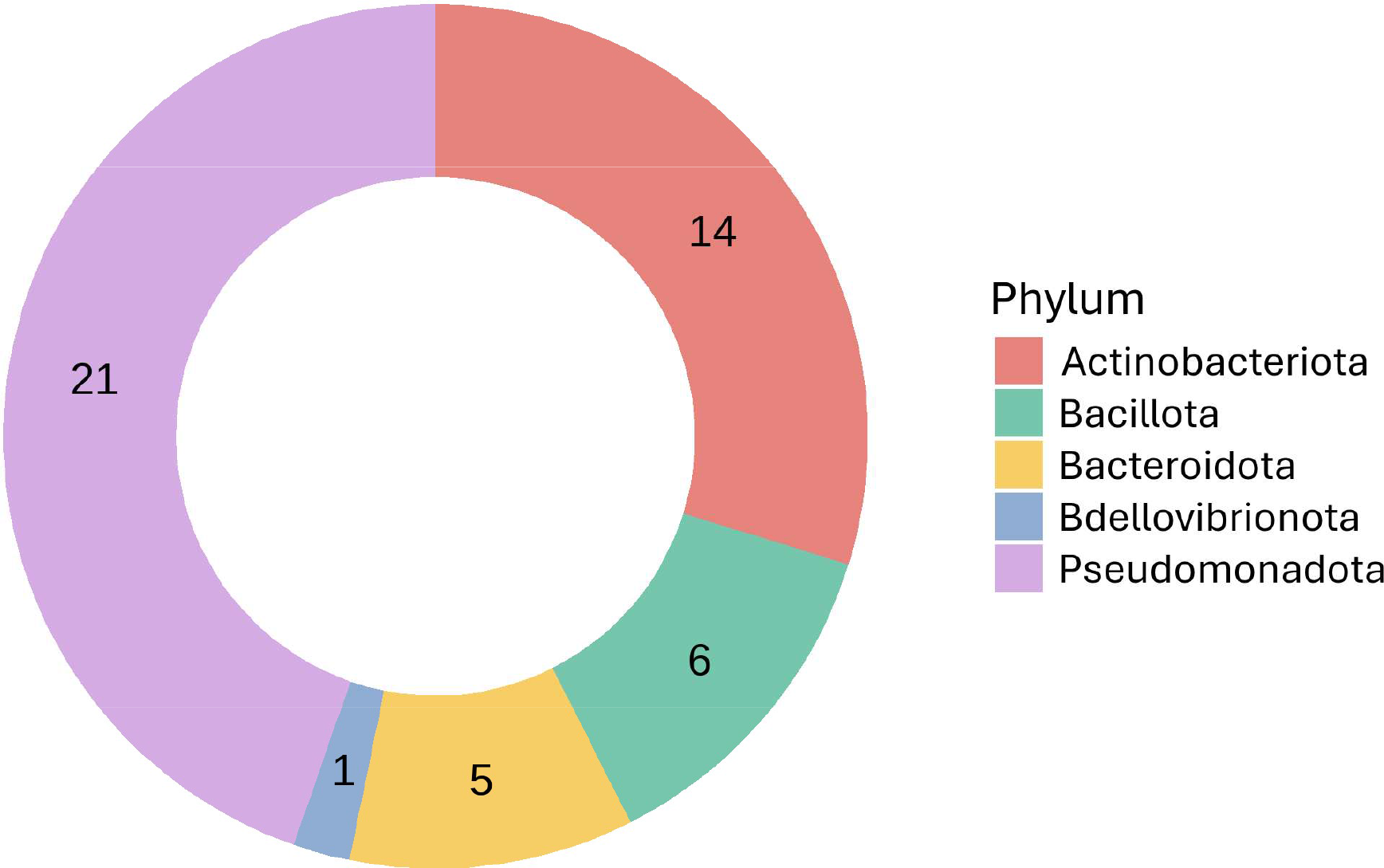
Taxonomic assignments to phyla of MAGs in the BSFL midgut community. The number of MAGs taxonomically assigned to each bacterial phyla through GTDB-Tk taxonomic classification.

