## Supplementary File 2 for "The gut bacterial community of black soldier fly larvae is a reservoir of antibiotic resistance and virulence genes": SF2_community_structure.html

Interactive taxonomic community structure of the BSFL midgut bacterial community


### Interactive taxonomic community structure of the BSFL midgut bacterial community

### 1 CAT classified interactive bacterial community structure

Figure 1.1: **CAT assigned kingdom-level community structure of the BSFL midgut bacterial community.** The total relative bacterial abundances of contigs assigned to bacterial kingdoms by CAT. Columns represent individual samples.

Figure 1.2: **CAT assigned phylum-level community structure of the BSFL midgut bacterial community.** The total relative bacterial abundances of contigs assigned to bacterial phyla by CAT. Phyla with total relative bacterial abundances of < 1 % were grouped into Other. Columns represent individual samples.

Figure 1.3: **CAT assigned class-, order-, and family-level community structure of the BSFL midgut bacterial community.** The total relative bacterial abundances of contigs belonging to taxonomic classifications of bacterial classes, orders, and families assigned by CAT were averaged across samples (n = 6). Node size and connection thickness is proportional to mean relative bacterial abundance.

Figure 1.4: **CAT assigned genus-level community structure of the BSFL midgut bacterial community.** The total relative bacterial abundances of contigs assigned to genera by CAT for each sample. Genera were ordered by descending mean relative bacterial abundance and mean relative bacterial abundances across samples were included in row labels (n = 6). Columns represent individual samples.

Figure 1.5: **CAT assigned species-level community structure of the BSFL midgut bacterial community.** The total relative bacterial abundances of contigs assigned to species by CAT for each sample. Species were ordered by descending mean relative bacterial abundance and mean relative bacterial abundances across samples were included in row labels (n = 6). Columns represent individual samples.

### 2 Kraken2 classified interactive bacterial community structure

Figure 2.1: **Kraken2 assigned kingdom-level community structure of the BSFL midgut bacterial community.** The total relative bacterial abundances of contigs assigned to bacterial kingdoms by kraken2. Columns represent individual samples.

Figure 2.2: **Kraken2 assigned phylum-level community structure of the BSFL midgut bacterial community.** The total relative bacterial abundances of contigs assigned to bacterial phyla by kraken2. Phyla with total relative bacterial abundances of < 1 % were grouped into Other. Columns represent individual samples.

Figure 2.3: **Kraken2 assigned class-, order-, and family-level community structure of the BSFL midgut bacterial community.** The total relative bacterial abundances of contigs belonging to taxonomic classifications of bacterial classes, orders, and families assigned by Kraken2 were averaged across samples (n = 6). Node size and connection thickness is proportional to mean relative bacterial abundance.

Figure 2.4: **Kraken2 assigned genus-level community structure of the BSFL midgut bacterial community.** The total relative bacterial abundances of contigs assigned to genera by Kraken2 for each sample. Genera were ordered by descending mean relative bacterial abundance and mean relative bacterial abundances across samples were included in row labels (n = 6). Columns represent individual samples.

Figure 2.5: **Kraken2 assigned species-level community structure of the BSFL midgut bacterial community.** The total relative bacterial abundances of contigs assigned to species by Kraken2 for each sample. Species were ordered by descending mean relative bacterial abundance and mean relative bacterial abundances across samples were included in row labels (n = 6). Columns represent individual samples.
