## Supplementary File 4 for "The gut bacterial community of black soldier fly larvae is a reservoir of antibiotic resistance and virulence genes": MAG_interactive_taxonomies.html

Interactive MAG taxonomic classifications of the BSFL midgut community


### Interactive MAG taxonomic classifications of the BSFL midgut community

Figure 1: **Hierarchical structure of MAG taxonomic classifications in the BSFL midgut community.** The number of MAGs assigned to each hierarchical level of the GTDB-Tk taxonomic classification from central phylum to class to order to family to genus moving outwards

Table 1: **Genus and species taxonomic classification of MAGs in the BSFL midgut community.** Genus and species GTDB-Tk taxonomic classifications for MAGs in the BSFL midgut community with FastANI reference genome and pplacer reference genome accessions.

| MAG | Genus | Species | FastANI reference | pplacer reference |
| --- | --- | --- | --- | --- |
| bin.10 | Morganella | Morganella morganii | GCF\_019243775.1 | GCF\_019243775.1 |
| bin.101 | Bacteriovoracaceae\_UBA4096 | NA | NA | NA |
| bin.102 | Enterococcus | Enterococcus phoeniculicola | GCF\_000407505.1 | GCF\_000407505.1 |
| bin.103 | Sphingobacterium | Sphingobacterium paucimobilis | GCF\_000416985.1 | GCF\_000416985.1 |
| bin.105 | Oxalicibacterium | NA | NA | GCF\_014635065.1 |
| bin.109 | Microbacterium | NA | NA | NA |
| bin.110 | Phytobacter | Phytobacter massiliensis | GCF\_000321045.1 | GCF\_000321045.1 |
| bin.12 | Enterococcus | Enterococcus nangangensis | GCF\_005405245.1 | GCF\_005405245.1 |
| bin.16 | Comamonas | NA | NA | GCF\_902829245.1 |
| bin.2 | Weeksella | Weeksella massiliensis | GCA\_000751595.1 | GCA\_000751595.1 |
| bin.22 | Scrofimicrobium | NA | NA | NA |
| bin.25 | Brevibacterium | NA | NA | NA |
| bin.27 | Rhizobiaceae\_63-22 | NA | NA | NA |
| bin.28 | Paenochrobactrum | NA | NA | GCF\_014205685.1 |
| bin.29 | Enterococcus | Enterococcus italicus | GCF\_000185365.1 | GCF\_000185365.1 |
| bin.3 | Scrofimicrobium | NA | NA | NA |
| bin.32 | Bordetella | Bordetella sp009763255 | GCF\_009763255.1 | GCF\_009763255.1 |
| bin.33 | Ochrobactrum | Ochrobactrum teleogrylli | GCF\_006376685.1 | GCF\_006376685.1 |
| bin.34 | Providencia | Providencia stuartii | GCA\_016618195.1 | GCA\_016618195.1 |
| bin.35 | Pauljensenia | Pauljensenia polynesiensis | GCF\_000820725.1 | GCF\_000820725.1 |
| bin.37 | Comamonas | NA | NA | GCF\_002158865.1 |
| bin.4 | Sphingobacterium | Sphingobacterium wenxiniae | GCF\_900116225.1 | GCF\_900116225.1 |
| bin.40 | Enterococcus | Enterococcus diestrammenae | GCF\_016908885.1 | GCF\_016908885.1 |
| bin.47 | Flavobacterium | Flavobacterium odoratimimum | GCF\_001485415.1 | GCF\_001485415.1 |
| bin.49 | Microbacterium | NA | NA | GCF\_005347485.1 |
| bin.5 | Enterococcus | Enterococcus gallinarum | GCF\_001544275.1 | GCF\_001544275.1 |
| bin.56 | Pelethocola | NA | NA | GCA\_018866245.1 |
| bin.57 | Ochrobactrum | NA | NA | NA |
| bin.58 | Cellulosimicrobium | Cellulosimicrobium funkei | GCF\_004519295.1 | GCF\_004519295.1 |
| bin.60 | Shinella | NA | NA | GCF\_003574625.1 |
| bin.61 | Providencia | Providencia rettgeri | GCF\_013255915.1 | GCF\_013255915.1 |
| bin.66 | Agromyces | Agromyces mediolanus | GCF\_014648575.1 | GCF\_014648575.1 |
| bin.67 | Microbacterium | NA | NA | GCF\_003327285.1 |
| bin.68 | Kosakonia | Kosakonia sp000410515 | GCF\_000410515.1 | GCF\_000410515.1 |
| bin.7 | Sphingobacterium | Sphingobacterium daejeonense | GCF\_901472535.1 | GCF\_901472535.1 |
| bin.72 | Proteus | Proteus mirabilis | GCF\_000160755.1 | GCF\_000160755.1 |
| bin.77 | Microbacterium | NA | NA | NA |
| bin.78 | Achromobacter | Achromobacter xylosoxidans | GCF\_001457475.1 | GCF\_001457475.1 |
| bin.82 | Microbacterium | NA | NA | NA |
| bin.85 | Comamonas | Comamonas sp002472915 | GCA\_002472915.1 | GCA\_002472915.1 |
| bin.87 | Pseudomonas | NA | NA | NA |
| bin.88 | Stenotrophomonas | NA | NA | NA |
| bin.9 | Kerstersia | Kerstersia gyiorum | GCF\_004216755.1 | GCF\_004216755.1 |
| bin.92 | Corynebacterium | Corynebacterium provencense | GCA\_900049755.1 | GCA\_900049755.1 |
| bin.93 | Scrofimicrobium | Scrofimicrobium canadense | GCF\_009696615.1 | GCF\_009696615.1 |
| bin.98 | Methylobacillus | NA | NA | GCF\_000013705.1 |
| bin.99 | Salana | NA | NA | GCF\_003751805.1 |
