## Supplementary File 5 for "The gut bacterial community of black soldier fly larvae is a reservoir of antibiotic resistance and virulence genes": SF5_MAG_network.html

Interactive network of ARG and VF genes in MAGs of the BSFL midgut community


### Interactive network of ARG and VF genes in MAGs of the BSFL midgut community

Figure 1: **ARG and VF encoding MAG taxa.** The ARG and VF genes encoded by taxa with MAGs. Unique taxa are represented by dark grey large nodes with ARG nodes in green, and VF nodes in orange. Grey links represent genes identified in the MAGs of a single taxon, red links represent genes identified in the MAGs of more than one taxon. Hovering over nodes provides additional information on drug classes that genes confer resistance to and the mechanism of resistance for ARG nodes, and virulence factor class for VF nodes.
