## Supplementary File 6 for "The gut bacterial community of black soldier fly larvae is a reservoir of antibiotic resistance and virulence genes": SF6_manuscript_code.Rmd

````
---
title: "The gut bacterial community of black soldier fly larvae is a reservoir of antibiotic resistance and virulence genes"
output: 
  bookdown::word_document2:
    reference_docx: "../Rmarkdown resources/word-styles-reference-manuscript.docx"
    number_sections: false
bibliography: 
  - "../Rmarkdown resources/resistome.bib"
  - "../Rmarkdown resources/bioinformatics.bib"
  - "../Rmarkdown resources/r_packages.bib"
csl: "../Rmarkdown resources/clean.csl"
---

Roma Davis^1,§^, Scott Conor JR^1,§^, Brilli Matteo^1^, Sequino Giuseppina^2^, Esposito Alessia^2^, De Filippis Francesca^2,3^, Tettamanti Gianluca^4,5^, Casartelli Morena^1,5,\*^, Caccia Silvia^1,3,\*^

^1^Department of Biosciences, University of Milan, Milan, Italy

^2^Department of Agricultural Sciences, University of Naples Federico II, Portici, Italy

^3^Task Force on Microbiome Studies, University of Naples Federico II, Naples, Italy

^4^Department of Biotechnology and Life Sciences, University of Insubria, Varese, Italy

^5^Interuniversity Centre for Studies on Bioinspired Agro-environmental Technology (BAT Centre), University of Naples Federico II, Portici, Italy

<br>

^§^ These authors have equally contributed to this work.


^\*^ Corresponding authors: 

Morena Casartelli, Department of Biosciences, University of Milan, via Celoria 26, 20133, Milan, Italy. email address:

Silvia Caccia, Department of Biosciences, University of Milan, via Celoria 26, 20133, Milan, Italy. email address:

\newpage

```{r setup, include=FALSE}
knitr::opts_chunk$set(echo = FALSE, warning = FALSE, message = FALSE)
set.seed(202507)
```

```{r packages}
library(knitr)
library(showtext)
library(dplyr)
library(tidyverse)
library(purrr)
library(magrittr)
library(tidyr)
library(stringr)
library(data.table)
library(flextable)
library(ggplot2)
library(patchwork)
library(ggraph)
library(grid)
library(gridExtra)
library(cowplot)
library(pheatmap)
library(stringdist)
library(ggVennDiagram)
library(igraph)
library(ggraph)
library(cowplot)
library(ggalluvial)
library(plotly)
library(htmlwidgets)
library(webshot2)
library(tidygraph)
library(viridis)
library(magick)
library(readxl)

font_add("Aptos", "../Rmarkdown resources/Aptos.ttf")
showtext.auto()
par(family = "Aptos")
options(scipen = 999)

wes_colours <- c("#D73027", "#1B9E77", "#F2AD00", "#4575B4", "#B775D1", "#E66100", "#A6761D", "#CAB2D6", "#FDC086", "#33A02C","#E7298A", "#1F78B4")
# Function to select the required number of colors
my_wes_palette <- function(n) {
  wes_colours[seq_len(n)]  # Extract first 'n' colors
}

# Function to add opacity (blend with white)
add_opacity <- function(color, alpha = 0.5) {
  rgb_col <- col2rgb(color)
  rgb(rgb_col[1, ], rgb_col[2, ], rgb_col[3, ], alpha = alpha * 255, maxColorValue = 255)
}

num_to_word <- function(x) {
  words <- c(
    "one", "two", "three", "four", "five",
    "six", "seven", "eight", "nine"
  )
  
  # If x is a whole number between 1 and 9, return word
  if (is.numeric(x) && x %% 1 == 0 && x >= 1 && x <= 9) {
    return(words[x])
  }
  
  # Otherwise return the number unchanged
  return(x)
}
```

```{r sup-files}
# Define supplementary file titles
supp_files <- c(
  "Shotgun metagenomic read quality report",
 "Interactive taxonomic community structures",
 "ARG, VF, and Plasmid database",
 "Interactive MAG taxonomic classification",
 "Interactive MAG network plot of ARG and VF genes",
 "R Markdown manuscript file"
)

# Create a function to return "Supplementary file X" for a given title
get_supp <- function(title) {
  index <- match(title, supp_files)
  if (!is.na(index)) {
    return(paste("Supplementary File", index))
  } else {
    stop("Title not found in supplementary files.")
  }
}
```

```{r tabsetup}
#set the table caption styling
set_flextable_defaults(font.family = "Aptos",
                       font.size = 11, 
                       digits = 0, 
                       border.color = "#000000",
                       padding.bottom = 1,
                       padding.top = 1,
                       padding.left = 3,
                       padding.right = 1)
style_flextable <- function(ft) {
  ft %>%
    bold(part = "header") %>%  # Make header row bold
    bg(i = seq(2, nrow(ft$body$dataset), by = 2), j = NULL, bg = "grey90", part = "body") # Alternating grey rows
}
```

# Abstract

Antimicrobial resistance (AMR) is a serious threat to global health. Agricultural practices that have contributed greatly to AMR spread urgently require innovation to address this issue, and more broadly challenges of sustainability and environmental concern. The larvae of black soldier fly (BSFL), *Hermetia illucens*, are considered a promising resource for advancing sustainable and circular agri-food systems given their ability to bioconvert organic waste streams into protein- and lipid-rich biomass suitable for feed applications and the use of the rearing residues as organic fertilisers. However, despite their emerging industrial applications, the risks of antibiotic resistance spread through their use remain underexplored. To elucidate this aspect, the profiles of antibiotic resistance genes (ARGs) and virulence factors (VFs), and their occurrence on plasmids were predicted from the midgut bacterial community of BSFL. Shotgun metagenomics revealed candidate resistance genes for 26 classes of antibiotics, and virulence via nine mechanisms, with taxa belonging to the Pseudomonadota phylum as the dominant contributors. Highly relevant to public health was the identification of genes encoding resistance to carbapenem class antibiotics in bacterial genomes and mobile plasmids. Reconstruction of metagenomes enabled more precise taxonomic resolution and revealed taxa harbouring multiple resistance and virulence genes, including a *Pseudomonas* species with 42 VFs and seven ARGs. Notably, for the first time antibiotic resistant bacterial species were isolated from the gut microbiota of BSFL, validating and complementing the results obtained *in silico*. Together, this work represents a comprehensive profile of the BSFL midgut bacterial resistome, while also providing relevant context on virulence and mobility. Importantly, it emphasises the urgent need to adopt strategies to mitigate potential risks associated with the development of emerging technologies associated to the use of BSFL and derived products.

\newpage

# Introduction

The spread of antimicrobial resistance genes (ARGs) has become a crucial global health threat, menacing the efficacy of the treatment and prevention of infections. Roughly 5 million deaths were associated with bacterial antimicrobial resistance (AMR) in 2019, and this number is projected to increase up to 10 million by 2050 [@ho2025]. Among other factors, agricultural practices and livestock production have greatly contributed to the spread of ARGs in the environment, with recent demands for a reduction in the use of antibiotics and the establishment of more sustainable processes [@wee2020; @ho2025]. ARG dissemination occurs via horizontal gene transfer of ARG-encoding DNA either through conjugation-based transfer of ARG-carrying mobile genetic elements [@wang2024; @jian2021; @munshi2025], transformation through the uptake of extracellular DNA [@bender2022], or other bacteriophage and membrane vesicle-based mechanisms [@tao2022]. Additionally, the horizontal transfer and dissemination of virulence factor (VF) genes through the same mechanisms is of growing concern as these VF genes might provide a selective advantage to ARG-carrying bacteria by improving growth, reproduction and resistance in stressful environments [@niu2013], while potentially enabling pathogenicity [@zhou2024; @fang2024].

The European Union's concern about this problem is highlighted by the One Health Action Plan against AMR which aims to improve the surveillance of ARGs and reduce antibiotic use by linking human health to ecosystem, crop, and livestock conditions [@EU2017_893]. However, as highlighted by the adoption of the circular economy action plan for sustainable growth by the European Union, the spread of AMR and pathogens are not the sole challenges facing the agricultural industry [@europeancommission2020]. The urgent need to innovate practices in the food system, while simultaneously improving the circularity of waste streams to reduce the overall waste generation, must also consider the AMR risks associated with any innovation. One such innovation is the development of insect-based biorefineries to support circular economy processes [@kim2025; @drewery2022; @mannaa2023].

Insects have emerged as agents in the renovation of agricultural practices and livestock production towards sustainability [@bruno2025; @kim2025; @mannaa2023]. Among them, the Black Soldier Fly (BSF) *Hermetia illucens* is increasingly recognised as a model for a wide range of biotechnological applications [@bruno2025; @tettamanti2024; @caccia2025]. Principally, scientific research into BSF larvae (BSFL) has focused on their capability to efficiently bioconvert organic wastes and byproducts into biomass rich in proteins and lipids that can be used in the feed and food sectors [@bonelli2020; @kim2025; @drewery2022]. Proteins and lipids from BSFL are also suitable for bioplastic and biodiesel production, respectively [@setti2020; @bruno2025; @jung2022]. Additionally, the frass (i.e., the residues at the end of insect rearing) is suitable as fertiliser [@jenkins2023; @lomonaco2024]. Therefore, bioconversion processes driven by BSFL represent a powerful means for creating circular-economy value chains in which waste is transformed into valuable resources.

The production of BSF biomass from waste streams is, however, strictly regulated in the European Union. For example, the use of BSF meals in the feed sector is restricted to insects reared on substrates of vegetable origin or specifically allowed materials of animal origin considered as microbiologically and chemically safe [@EU2017_893; @lahteenmaki-uutela2021]. In other countries, regulation of insect rearing, production processes, and applications may vary considerably [@lahteenmaki-uutela2021]. The capability of BSFL to efficiently bioconvert a wide range of organic materials, including nutritionally poor substrates, is derived from their saprophagous feeding habits and the high functional plasticity of their gut and the associated microbiota [@bonelli2020; @defilippis2023; @bruno2025; @filippis2024]. The complex gut microbial community structure is also a reservoir of ARGs, VFs, and plasmids with the profiles of these genes influenced by rearing conditions and thus a point of concern for safety [@defilippis2023; @bohm2024; @bruno2019; @rong2024; @vandeweyer2023]. This aspect must be clarified to establish safe and informed BSFL-mediated processes, especially in low- and middle-income countries, where the AMR-related morbidity or mortality are particularly severe and regulation on insect rearing may be lenient [@ho2025; @lahteenmaki-uutela2021].

Despite the growing relevance of the topic, only a few studies have investigated the diversity and distribution of ARGs in the BSFL microbiome [@zhao2023; @zhao2023a; @rong2024; @schokker2025; @bohm2024]. These studies, based on metagenomic analyses, provide important insight into the fate of ARGs across different waste streams including chicken and swine manure, residuals from the dairy and potato industries, and sludge. All investigations showed diet-dependent changes in both microbial community structure and resistome profiles with AMR bacteria still present in the final products of bioconversion processes, which may pose a risk in their use [@zhao2023; @zhao2023a; @rong2024]. Indeed, putatively pathogenic taxa with multiple ARGs were found in the midgut metagenomes of larvae reared on dairy wastes and sewage sludge, with the authors highlighting their role in ARG spread [@schokker2025]. However, these studies did not fully characterise the co-occurrence of VFs and ARGs in microorganisms of the BSFL midgut community. Most importantly, *in silico* predictions were not supported by *in vivo* microbiological evidence.

Therefore, effectively profiling the midgut resistome and virulome of BSFL is not only valuable to increase basic knowledge on this topic, but will also provide an assessment of the potential environmental and public health risks associated with the biotechnological applications based on BSFL. Herein, remaining knowledge gaps were addressed by establishing a comprehensive platform of knowledge was established based on metagenomic analysis of the ARGs and VFs, and the mobile plasmids encoded within the BSFL midgut. The investigation of antibiotic resistance was extended by the isolation and species-level identification of antibiotic-resistant strains from the BSFL midgut. Through these analyses, we present a comprehensive profile of the BSFL midgut resistome together with virulome and plasmid profiles, to provide a valuable resource for the BSFL research community for the informed use of these larvae in agricultural processes.

\newpage

# Materials and methods

```{r contig-import, eval=FALSE}
# Read in the named contig taxonomic classifications from CAT annotation tool
contig_tax <- readLines("Bioinformatics/contig_classification/CAT_output.contig2classification.named.txt") %>%
  { 
    header <- .[1] %>% gsub("^#\\s*", "", .) %>% strsplit("\t") %>% .[[1]]
    data_lines <- .[-1]

    split_lines <- data_lines %>% map(~ strsplit(.x, "\t")[[1]])
    max_cols <- split_lines %>% map_int(length) %>% max()
    padded <- split_lines %>% map(~ c(.x, rep(NA, max_cols - length(.x))))

    df <- do.call(rbind, padded) %>% as.data.frame(stringsAsFactors = FALSE)
    colnames(df) <- c(header, paste0("V", seq_len(ncol(df) - length(header))))
    df
  } %>% 
  select(-c("lineage", "lineage scores (f: 0.5)", "full lineage names"))

# Read in the contig depth information
contig_depth <- read.delim("Bioinformatics/coassembly_coverage/depth.txt", header = T, sep = "\t") %>% 
  rename(contig = contigName, length = contigLen, Sum_Avg_depth = totalAvgDepth, STD_1_Avg = STD_1.sorted, STD_1_Var = STD_1.sorted.var, STD_2_Avg = STD_2.sorted, STD_2_Var = STD_2.sorted.var, STD_4_Avg = STD_4.sorted, STD_4_Var = STD_4.sorted.var, STD_5_Avg = STD_5.sorted, STD_5_Var = STD_5.sorted.var, STD_7_Avg = STD_7.sorted, STD_7_Var = STD_7.sorted.var, STD_8_Avg = STD_8.sorted, STD_8_Var = STD_8.sorted.var)
```

```{r contig-tax-parse, eval=FALSE}
# Define non-valid ranks
non_valid_ranks <- c(
  "2020",
  "L",
  "2022",
  "in: enterobacteria",
  "Halichondria",
  "in: firmicutes",
  "in: high G+C Gram-positive bacteria",
  "in: a-proteobacteria"
)
# Step 1: Add row_id and extract taxonomic data
long_tax <- contig_tax %>%
  mutate(row_id = row_number()) %>%
  pivot_longer(
    cols = starts_with("V"),
    names_to = "column",
    values_to = "tax_string"
  ) %>%
  filter(!is.na(tax_string)) %>%
  mutate(
    rank = str_extract(tax_string, "(?<=\\().+?(?=\\))"),
    name = str_remove(tax_string, " \\(.+\\): .*"),
    conf = str_extract(tax_string, "\\d+\\.\\d+$")
  ) %>%
  filter(!is.na(rank)) %>%
  filter(!rank %in% non_valid_ranks) %>%   # <--- exclude these ranks here
  group_by(row_id, rank) %>%
  summarise(
    taxon = paste(name, collapse = "; "),
    confidence = paste(conf, collapse = "; "),
    .groups = "drop"
  ) %>%
  ungroup()

# Step 2: Pivot taxon and confidence info separately
taxon_part <- long_tax %>%
  select(row_id, rank, taxon) %>%
  pivot_wider(
    names_from = rank,
    values_from = taxon,
    names_glue = "{rank}"
  )

conf_part <- long_tax %>%
  select(row_id, rank, confidence) %>%
  pivot_wider(
    names_from = rank,
    values_from = confidence,
    names_glue = "{rank}_conf"
  )

# Step 3: Join everything back
cleaned_tax_table <- contig_tax %>%
  select(contig, classification, reason) %>%
  mutate(row_id = row_number()) %>%
  left_join(taxon_part, by = "row_id") %>%
  left_join(conf_part, by = "row_id") %>%
  select(contig, classification, reason, everything(), -row_id) %>% 
  rename(no_rank = "no rank", species_group = "species group", species_subgroup = "species subgroup", no_rank_conf = "no rank_conf", species_group_conf = "species group_conf", species_subgroup_conf = "species subgroup_conf")
```

```{r abricate-card-import, eval=FALSE}
# Import the outputs from the Abricate predictions of Antibiotic resistance genes from the CARD database, also rename to make more accessible
contig_card <- read.delim("Bioinformatics/abricate/abricate_card_results_STD_contigs.tab", sep = "\t", header = TRUE) %>% 
  rename(contig = SEQUENCE, start = START, end = END, strand = STRAND, gene = GENE, coverage = COVERAGE, coverage_map = COVERAGE_MAP, gaps = GAPS, coverage_percentage = X.COVERAGE, pident = X.IDENTITY, accession = ACCESSION, product = PRODUCT, resistance = RESISTANCE) %>% 
  select(contig, start, end, strand, gene, coverage, coverage_map, gaps, coverage_percentage, pident, accession, product, resistance)

# Desired order of columns per ARG
arg_fields <- c("start", "end", "strand", "gene",
                "coverage", "coverage_map", "gaps",
                "coverage_percentage", "pident", "accession",
                "product", "resistance")

# Get all ARG indices
arg_indices <- contig_card %>%
  count(contig) %>%
  pull(n) %>%
  max()
# Get desired order of columns
desired_arg_order <- c("contig")
for (i in 1:arg_indices) {
  desired_arg_order <- c(desired_arg_order, paste0("arg", i, "_", arg_fields))
}

# Number the args as some contigs have multiple (up to 5 here)
contig_card_wide <- contig_card %>%
  group_by(contig) %>%
  mutate(arg_index = row_number()) %>%
  ungroup() %>%
  pivot_wider(
    id_cols = contig,
    names_from = arg_index,
    values_from = c(
      start, end, strand, gene,
      coverage, coverage_map, gaps,
      coverage_percentage, pident, accession,
      product, resistance
    ),
    names_glue = "arg{arg_index}_{.value}"
  ) %>% 
  select(any_of(desired_arg_order))
```

```{r abricate-vfdb-import, eval=FALSE}
# Import the outputs from the Abricate predictions of Antibiotic resistance genes from the VFDB database, also rename to make more accessible
contig_vfdb <- read.delim("Bioinformatics/abricate/abricate_VFDB_results_STD_contigs.tab", sep = "\t", header = TRUE) %>% 
  rename(contig = SEQUENCE, start = START, end = END, strand = STRAND, gene = GENE, coverage = COVERAGE, coverage_map = COVERAGE_MAP, gaps = GAPS, coverage_percentage = X.COVERAGE, pident = X.IDENTITY, accession = ACCESSION, product = PRODUCT) %>% 
  select(contig, start, end, strand, gene, coverage, coverage_map, gaps, coverage_percentage, pident, accession, product)

# Desired order of columns per ARG
vf_fields <- c("start", "end", "strand", "gene",
                "coverage", "coverage_map", "gaps",
                "coverage_percentage", "pident", "accession",
                "product")

# Get all ARG indices
vf_indices <- contig_vfdb %>%
  count(contig) %>%
  pull(n) %>%
  max()
# Get desired order of columns
desired_vf_order <- c("contig")
for (i in 1:vf_indices) {
  desired_vf_order <- c(desired_vf_order, paste0("vf", i, "_", vf_fields))
}

# Number the args as some contigs have multiple (up to 5 here)
contig_vfdb_wide <- contig_vfdb %>%
  group_by(contig) %>%
  mutate(vf_index = row_number()) %>%
  ungroup() %>%
  pivot_wider(
    id_cols = contig,
    names_from = vf_index,
    values_from = c(
      start, end, strand, gene,
      coverage, coverage_map, gaps,
      coverage_percentage, pident, accession,
      product
    ),
    names_glue = "vf{vf_index}_{.value}"
  ) %>% 
  select(any_of(desired_vf_order))
```

```{r abricate-plasmid-import, eval=FALSE}
# Read in the plasmid reports and modify columns, also remove the duplicate assignments for a single contigs as these are just artifacts of the plasmid finder database
contig_plasmids <- read.delim("Bioinformatics/abricate/abricate_PlasmidF_results_STD_contigs.tab", header = TRUE) %>% 
  rename(contig = SEQUENCE, PF_start = START, PF_end = END, PF_strand = STRAND, PF_gene = GENE, PF_coverage = COVERAGE, PF_coverage_map = COVERAGE_MAP, PF_gaps = GAPS, PF_coverage_percentage = X.COVERAGE, PF_pident = X.IDENTITY, PF_accession = ACCESSION, PF_product = PRODUCT) %>% 
  select(contig, PF_start, PF_end, PF_strand, PF_gene, PF_coverage, PF_coverage_map, PF_gaps, PF_coverage_percentage, PF_pident, PF_accession, PF_product) %>% 
  group_by(contig) %>% 
  slice_max(order_by = PF_pident, with_ties = FALSE) %>% 
  ungroup()
```

```{r contig-merge, eval=FALSE}
# Merge the taxonomic info and the depth data together for each contig. There are 89 more lines in the depth file as 89 contigs could not be assigned a taxonomic classification so we will populate these rows with NAs. Then will clean up the order so that it makes sense
contig_merged <- contig_depth %>%
  left_join(cleaned_tax_table, by = "contig") %>% 
  select("contig", "length", "Sum_Avg_depth", "STD_1_Avg", "STD_1_Var", "STD_2_Avg", "STD_2_Var", "STD_4_Avg", "STD_4_Var", "STD_5_Avg", "STD_5_Var", "STD_7_Avg", "STD_7_Var", "STD_8_Avg", "STD_8_Var", "classification", "reason", "no_rank", "clade", "superkingdom", "kingdom", "subkingdom", "phylum", "subphylum", "superclass", "class", "subclass", "infraclass", "cohort", "superorder", "order", "suborder", "infraorder", "parvorder", "superfamily", "family", "subfamily", "tribe", "subtribe", "genus", "species_group", "species_subgroup", "species", "subspecies", "varietas", "strain", "no_rank_conf", "clade_conf", "superkingdom_conf", "kingdom_conf", "subkingdom_conf",
  "phylum_conf", "subphylum_conf", "superclass_conf", "class_conf", "subclass_conf",
  "infraclass_conf", "cohort_conf", "superorder_conf", "order_conf", "suborder_conf",
  "infraorder_conf", "parvorder_conf", "superfamily_conf", "family_conf", "subfamily_conf",
  "tribe_conf", "subtribe_conf", "genus_conf", "species_group_conf", "species_subgroup_conf",
  "species_conf", "subspecies_conf", "varietas_conf", "strain_conf")

# Also merge the Abricate CARD and VFDB results
contig_db_arg_vf <- contig_merged %>% 
  left_join(contig_card_wide, by = "contig") %>% 
  left_join(contig_vfdb_wide, by = "contig") %>% 
  left_join(contig_plasmids, by = "contig")

# This is all the information that is now ready to filter in anyway for further analysis. Let's write this out to a .csv file so that we can save time reading files in for the future
write.table(contig_db_arg_vf, "Bioinformatics/contig_classification/contig_tax_depth_arg_vf_plasmids_db.txt", quote = FALSE, row.names = FALSE, sep = "\t")
```

```{r read-contig-db, eval=FALSE}
# Read in the .txt file of the database
contig_db <- read.delim("Bioinformatics/contig_classification/contig_tax_depth_arg_vf_plasmids_db.txt", header = T, sep = "\t")
```

```{r contig-tax-clean, eval=FALSE}
# Okay, the database has everything, but we want to clean the taxonomic ranks to those with a confidence score of >= 0.5, which is actually the default for CAT automatically so no taxonomic classifications will have a score less than this. Each rank has it's own column and corresponding confidence column, however some columns have multiple entries
filter_taxonomy_by_conf_dt <- function(dt, threshold = 0.5) {
  # Ensure data is a data.table
  setDT(dt)

  # Taxonomic columns with _conf columns
  ranks <- names(dt)[!str_ends(names(dt), "_conf") & str_c(names(dt), "_conf") %in% names(dt)]

  for (rank in ranks) {
    conf_col <- paste0(rank, "_conf")

    dt[, c(rank, conf_col) := {
      # Coerce to character before splitting
      tax_split <- strsplit(as.character(get(rank)), "; ")
      conf_split <- strsplit(as.character(get(conf_col)), "; ")

      result <- mapply(function(t, c) {
        c <- as.numeric(c)
        if (length(c) != length(t)) return(c(NA_character_, NA_character_))
        keep <- which(c >= threshold)
        if (length(keep) == 0) return(c(NA_character_, NA_character_))
        c(paste(t[keep], collapse = "; "), paste(c[keep], collapse = "; "))
      }, tax_split, conf_split, SIMPLIFY = TRUE)

      list(result[1, ], result[2, ])
    }]
  }

  return(dt)
}

# Apply function to clean taxonomy columns of database
contig_db_cleaned <- filter_taxonomy_by_conf_dt(as.data.table(contig_db), threshold = 0.5)

# Write this to a .txt so I don't have to do this again
write.table(contig_db_cleaned, "Bioinformatics/contig_classification/contig_tax_depth_arg_vf_db_clean.txt", quote = FALSE, row.names = FALSE, sep = "\t")
```

```{r read-contig-db-clean}
# Read in the contig database with the cleaned taxonomies
contig_db_clean <- read.delim("Bioinformatics/contig_classification/contig_tax_depth_arg_vf_db_clean.txt", header = T, sep = "\t")

# Firstly define the sample columns, which will be useful later on also
sample_cols <- c("STD_1_Avg", "STD_2_Avg", "STD_4_Avg", "STD_5_Avg", "STD_7_Avg", "STD_8_Avg")

# Normalise the relative abundances for future use
contig_db_norm <- contig_db_clean %>% 
  mutate(across(all_of(sample_cols),
                ~ (.x / sum(.x)) * 100,
                .names = "norm_{.col}"))
```

```{r contig-bacteria}
# For profiling the resistome, we want to focus on the bacterial community, and therefore we should filter to bacteria
# Firstly, can filter based on the 'superkingdom' column to Bacteria
contig_bacteria <- contig_db_clean %>% 
  filter(superkingdom == "Bacteria")

# Calculate the number and percentage of contigs assigned to the superkingdom bacteria
contigs_bacteria_num <- nrow(contig_bacteria)
contigs_bacteria_percentage <- (nrow(contig_bacteria) / nrow(contig_db_clean)) * 100

# Calculate proportions of bacterial read depths
bacteria_proportions <- contig_db_clean %>%
  summarise(across(all_of(sample_cols), sum, .names = "total_{.col}")) %>%
  bind_cols(
    contig_db_clean %>%
      filter(superkingdom == "Bacteria") %>%
      summarise(across(all_of(sample_cols), sum, .names = "bacteria_{.col}"))
  ) %>%
  mutate(across(starts_with("bacteria_"), 
                ~ .x / get(sub("bacteria_", "total_", cur_column())) * 100, 
                .names = "prop_{.col}")) %>%
  select(starts_with("prop_")) %>%
  rename_with(~ gsub("prop_bacteria_", "", .x)) %>%
  pivot_longer(everything(), names_to = "sample", values_to = "bacteria_percent") %>% 
  as.data.frame()

# Calculate the mean
bacteria_proportion_mean <- mean(bacteria_proportions$bacteria_percent)
bacteria_proportion_sd <- sd(bacteria_proportions$bacteria_percent)
```

```{r bacteria-norm}
# Let's now normalise the read depths for bacterial contigs
contig_bacteria_norm <- contig_bacteria %>% 
  mutate(across(all_of(sample_cols),
                ~ (.x / sum(.x)) * 100,
                .names = "norm_{.col}"))
```

```{r contig-bacteria-tax}
# Okay firstly I want to just investigate the taxonomic profile of the bacteria we have identified
# So let's take only the columns I am interested in for plotting the taxonomic profiles from the normalised results also let's remove the asterisks (these happen when e.g. there is only one genus within a family, so it can only be assigned to one thing)
contig_bacteria_tax <- contig_bacteria_norm %>% 
  select(c(contig, norm_STD_1_Avg, norm_STD_2_Avg, norm_STD_4_Avg, norm_STD_5_Avg, norm_STD_7_Avg, norm_STD_8_Avg, superkingdom, kingdom, phylum, class, order, family, genus, species)) %>% 
  rename(Superkingdom = superkingdom, Kingdom = kingdom, Phylum = phylum, Class = class, Order = order, Family = family, Genus = genus, Species = species) %>% 
  mutate(across(c(Superkingdom, Kingdom, Phylum, Class, Order, Family, Genus, Species), 
                ~ gsub("\\*$", "", .)
                )
         ) %>% 
  mutate(across(c(Superkingdom, Kingdom, Phylum, Class, Order, Family, Genus, Species), 
                ~ gsub("Candidatus ", "", .)
                )
         )

# Now I want to see ow many contigs are assigned to each taxonomic level of interest and how many unique classifications there are at each taxonomic level
# Firstly let's define the taxonomy levels
taxonomy_levels <- c("Kingdom", "Phylum", "Class", "Order", "Family", "Genus", "Species")

# Summarise the taxonomies
contig_taxon_sum <- contig_bacteria_tax %>%
  select(all_of(taxonomy_levels)) %>% 
  summarize(across(everything(), 
                   list(
                     Contig_count = ~ sum(!is.na(.)),
                     Unique_Taxa = ~ n_distinct(.[!is.na(.)])
                   ))) %>%
  pivot_longer(cols = everything(), 
               names_to = c("Taxonomic level", "Metric"),
               names_sep = "_") %>% 
  pivot_wider(names_from = Metric, values_from = value) %>% 
  rename("Contigs assigned" = Contig, "Unique classifications" = Unique)
# Convert the taxonomy dataframe to a list for inline referencing
contig_taxon_stats <- contig_taxon_sum %>%
  split(.$`Taxonomic level`) %>%
  lapply(function(x) list(Contigs = x$`Contigs assigned`, Unique = x$`Unique classifications`))
```

```{r bacteria-kingdom}
# Let's see the ditribution of the different bacterial kingdoms that the contigs are assigned to
bacteria_kingdom <- contig_bacteria_tax %>% 
  mutate(Kingdom = replace_na(Kingdom, "Unclassified")) %>% 
  count(Kingdom, name = "Count") %>% 
  mutate(Percentage = (Count / sum(Count)) * 100)
# Pull out the counts and percentage into a list for inline referencing
bacteria_kingdom_stats <- bacteria_kingdom %>%
  split(.$Kingdom) %>%
  lapply(function(x) list(Count = x$Count, Percentage = x$Percentage))

# I also want to sum the relative abundances for plotting the abundance profile of kingdoms
kingdom <- contig_bacteria_tax %>% 
  mutate(Kingdom = replace_na(Kingdom, "Unclassified")) %>% 
  select(c(contig, norm_STD_1_Avg, norm_STD_2_Avg, norm_STD_4_Avg, norm_STD_5_Avg, norm_STD_7_Avg, norm_STD_8_Avg, Kingdom)) %>% 
  pivot_longer(
    cols = -c(contig, Kingdom),  # Keep contig and Kingdom as fixed columns
    names_to = "Sample",     # Convert sample names to a single column
    values_to = "Abundance"  # Convert abundance values to a single column
  ) %>% 
  # Group contigs of the same Kingdom together
  group_by(Sample, Kingdom) %>%
  summarise(Abundance = sum(Abundance), .groups = "drop")

# Summarise the average relative abundances of each phylum
kingdom_sum <- kingdom %>% 
  group_by(Kingdom) %>%
  summarise(mean_abundance = mean(Abundance), .groups = "drop")
# Convert to a list for inline referencing
kingdom_sum_stats <- kingdom_sum %>%
  deframe()
```

```{r bacteria-phylum}
# Let's see the distribution of the different bacterial phyla that the contigs are assigned to
bacteria_phylum <- contig_bacteria_tax %>% 
  mutate(Phylum = replace_na(Phylum, "Unclassified")) %>% 
  count(Phylum, name = "Count") %>% 
  mutate(Percentage = (Count / sum(Count)) * 100)
# Pull out the contig counts and percentage into a list for inline referencing
bacteria_phylum_stats <- bacteria_phylum %>%
  split(.$Phylum) %>%
  lapply(function(x) list(Count = x$Count, Percentage = x$Percentage))

# I also want to sum the relative abundances for plotting the abundance profile of phyla
phylum <- contig_bacteria_tax %>% 
  mutate(Phylum = replace_na(Phylum, "Unclassified")) %>% 
  select(c(contig, norm_STD_1_Avg, norm_STD_2_Avg, norm_STD_4_Avg, norm_STD_5_Avg, norm_STD_7_Avg, norm_STD_8_Avg, Phylum)) %>% 
  pivot_longer(
    cols = -c(contig, Phylum),  # Keep contig and Phylum as fixed columns
    names_to = "Sample",     # Convert sample names to a single column
    values_to = "Abundance"  # Convert abundance values to a single column
  ) %>% 
  # Group contigs of the same Kingdom together
  group_by(Sample, Phylum) %>%
  summarise(Abundance = sum(Abundance), .groups = "drop")

# I want to change the low-abundance phyla contributing less than 1 % on average to "Other"
# Calculate mean abundance per Phylum
low_abundance_phyla <- phylum %>%
  group_by(Phylum) %>%
  summarise(mean_abundance = mean(Abundance), .groups = "drop") %>%
  filter(mean_abundance < 1) %>%
  pull(Phylum)

# Replace those phyla with "Other" and aggregate
phylum_clean <- phylum %>%
  mutate(Phylum = ifelse(Phylum %in% low_abundance_phyla, "Other", Phylum)) %>% 
  group_by(Sample, Phylum) %>%
  summarise(Abundance = sum(Abundance), .groups = "drop") %>%
  mutate(
    Phylum = factor(
      Phylum,
      levels = c(
        sort(setdiff(unique(Phylum), c("Other", "Unclassified"))),
        "Other",
        "Unclassified"
      )
    )
  )

# Summarise the average relative abundances of each phylum
phylum_sum <- phylum_clean %>% 
  group_by(Phylum) %>%
  summarise(mean_abundance = mean(Abundance), .groups = "drop")
# Convert to a list for inline referencing
phylum_sum_stats <- phylum_sum %>%
  deframe()

# Also calculate proportions of numbers of contigs for only contigs classified to the phylum level. This will be used later when comparing these proportions to contigs with ARGs or VFs
bacteria_phylum_classified <- contig_bacteria_tax %>% 
  filter(!is.na(Phylum)) %>% 
  count(Phylum, name = "Count") %>% 
  mutate(Percentage = (Count / sum(Count)) * 100)
```

```{r heatmap-genus-make}
# Take the abundances of different genus level classifications, ignoring the contigs which are unclassified at the genus level
genus_abundance <- contig_bacteria_tax %>% 
  mutate(Genus = ifelse(is.na(Genus), "Unclassified", Genus)) %>% 
  group_by(Genus) %>% 
  summarise(across(starts_with("norm_STD"), sum)) %>% 
  ungroup() %>% 
  rename("STD1" = norm_STD_1_Avg, "STD2" = norm_STD_2_Avg, 
         "STD4" = norm_STD_4_Avg, "STD5" = norm_STD_5_Avg, 
         "STD7" = norm_STD_7_Avg, "STD8" = norm_STD_8_Avg)

# Get top genera with abundance greater 0.1 %
top_genera <- genus_abundance %>%
  mutate(mean_abundance = rowMeans(across(starts_with("STD")))) %>%
  filter(mean_abundance > 0.1)

top_genera_mat <- top_genera %>% 
  select(-mean_abundance)

# Convert to matrix
genus_matrix <- as.matrix(top_genera[ , -1])  # drop Genus column
rownames(genus_matrix) <- top_genera$Genus   # use Genus as row names

# Calculate means of relative abundance and add to the genus rownames
mean_annotation <- signif(rowMeans(genus_matrix), digits = 3)

# Order rows by mean abundance:
genus_matrix_ordered <- genus_matrix[order(rowMeans(genus_matrix), decreasing = TRUE), ]

# Calculate pseudocount and log-transform ordered matrix:
min_val <- min(genus_matrix_ordered[genus_matrix_ordered > 0])
pseudo <- min_val / 10
log_genus_matrix <- log10(genus_matrix_ordered + pseudo)

# Round original (non-log) abundances for display
genus_labels <- format(round(genus_matrix_ordered, digits = 2), nsmall = 2)

# Now let's check the contigs which were unclassified at the genus level and make sure there are contigs classified to all phyla
genus_unclassified <- contig_bacteria_tax %>% 
  filter(is.na(Genus)) %>% 
  mutate(Phylum = ifelse(is.na(Phylum), "Unclassified", Phylum)) %>% 
  count(Phylum, name = "Count")
```

```{r community-plot-make, eval=FALSE}
# Set font
par(family = "Aptos")

# Sort the phyla alphabetically, excluding Unclassified
phylum_sorted <- sort(setdiff(levels(phylum_clean$Phylum), c("Other", "Unclassified")))
# Build color vector
phylum_colors <- setNames(
  c(my_wes_palette(length(phylum_sorted)), "grey10", "grey80"),
  c(phylum_sorted, "Other", "Unclassified")
)

phylum_plot <- ggplot(phylum_clean
                            , aes(x = Sample, y = Abundance, fill = Phylum)) +
  geom_bar(stat = "identity", position = "stack", size = 0.3, colour = "black", alpha = 0.5) +
  scale_fill_manual(values = phylum_colors) +
  theme_minimal(base_family = "Aptos") +
  theme(
    panel.grid.major = element_blank(),
    panel.grid.minor = element_blank(),
    axis.title.x = element_blank(),
    axis.title.y = element_text(size = 50, color = "black", lineheight = 0.3, margin = margin(r = 5)),
    axis.text.y = element_text(size = 44, color = "black"),
    axis.text.x = element_blank(),
    axis.line.y = element_line(color = "black", linewidth = 0.5),
    axis.line.x = element_blank(),
    axis.ticks.y = element_line(color = "black", linewidth = 0.5),
    axis.ticks.x = element_blank(),
    strip.text = element_text(size = 50),
    legend.position = "bottom",
    legend.title = element_text(size = 50, color = "black"),
    legend.text = element_text(size = 44, color = "black")
  ) +
  guides(
    fill = guide_legend(order = 1, nrow = 6, byrow = TRUE)
    ) +
  labs(
    y = "Relative bacterial abundance (%)",
    fill = "Phylum"
  )

ggsave(
  filename = "plots/phylum_barplot.png",
  plot = phylum_plot + theme(text = element_text(family = "Aptos")),
  width = 1200 / 300,   # convert pixels to inches
  height = 2100 / 300,  # convert pixels to inches
  dpi = 300,
  units = "in",
  bg = "white"
)

# Plot the heatmap
genus_heatmap <- pheatmap(log_genus_matrix,
                          cluster_rows = FALSE,
                          cluster_cols = FALSE,
                          scale = "none",
                          color = colorRampPalette(c("white", "#4575B4"))(100),
                          fontsize = 10,
                          main = "",
                          border_color = "white",
                          display_numbers = genus_labels,
                          number_color = "black",
                          fontsize_number = 9,
                          show_colnames = FALSE,
                          legend = FALSE,
                          silent = TRUE)

png(
  filename = "plots/genus_pheatmap.png",
  width = 1200,
  height = 2200,
  res = 300
)

# Draw the silent pheatmap manually
grid::grid.draw(genus_heatmap$gtable)

dev.off()

# Read images
phylum_img <- image_read("plots/phylum_barplot.png")
genus_img <- image_read("plots/genus_pheatmap.png")

# Annotate with labels
phylum_img <- image_annotate(phylum_img, "A", size = 100, font = "Aptos",
                       gravity = "northwest", color = "black", location = "+10+-20")
genus_img <- image_annotate(genus_img, "B", size = 100, font = "Aptos",
                       gravity = "northwest", color = "black", location = "+10+-20")

# Combine side by side
combined_community_img <- image_append(c(phylum_img, genus_img))

# Save final combined PNG
image_write(combined_community_img, path = "plots/combined_community_plot.png", format = "png")
```

```{r kraken-import, eval=FALSE}
# Import the kraken contig classification results where the confidence threshold was set to 0.2 and separate into different levels of taxonomic lineage
kraken <- read.delim("Bioinformatics/kraken/conf0.2_contig_lineage.tsv", sep = "\t", header = FALSE, stringsAsFactors = FALSE) %>% 
  rename(contig = V1, lineage = V3) %>% 
  separate_wider_delim(
    lineage,
    delim = ";",
    names_sep = "_",
    too_few = "align_start"
  )


# Import depths
# Read in the contig depth information
contig_depth <- read.delim("Bioinformatics/coassembly_coverage/depth.txt", header = T, sep = "\t") %>% 
  rename(contig = contigName, length = contigLen, Sum_Avg_depth = totalAvgDepth, STD_1_Avg = STD_1.sorted, STD_1_Var = STD_1.sorted.var, STD_2_Avg = STD_2.sorted, STD_2_Var = STD_2.sorted.var, STD_4_Avg = STD_4.sorted, STD_4_Var = STD_4.sorted.var, STD_5_Avg = STD_5.sorted, STD_5_Var = STD_5.sorted.var, STD_7_Avg = STD_7.sorted, STD_7_Var = STD_7.sorted.var, STD_8_Avg = STD_8.sorted, STD_8_Var = STD_8.sorted.var)
```

```{r kraken-depth-merge, eval=FALSE}
# Merge kraken taxonomic info to the depths for each contig, and then add the card and vfdb results
kraken_db_full <- contig_depth %>%
  left_join(kraken, by = "contig") %>% 
  left_join(contig_card_wide, by = "contig") %>% 
  left_join(contig_vfdb_wide, by = "contig") %>% 
  left_join(contig_plasmids, by = "contig")

# Write this db to a .txt to save for future use
write.table(kraken_db_full, "Bioinformatics/kraken/kraken_db_full.txt", quote = FALSE, row.names = FALSE, sep = "\t")
```

```{r kraken-db-import, eval=FALSE}
# Import the entire database
kraken_db_full <- read.delim("Bioinformatics/kraken/kraken_db_full.txt", header = TRUE, sep = "\t")

# Normalise the relative abundances for future use
kraken_db_norm <- kraken_db_full %>% 
  mutate(across(all_of(sample_cols),
                ~ (.x / sum(.x)) * 100,
                .names = "norm_{.col}"))
```

```{r full-db-make, eval=FALSE}
full_community_db <- contig_db_norm %>% 
  rename_with(~ paste0("CAT_", .), c("classification",
                "reason",
                "no_rank",
                "clade",
                "superkingdom",
                "kingdom",
                "subkingdom",
                "phylum",
                "subphylum",
                "superclass",
                "class",
                "subclass",
                "infraclass",
                "cohort",
                "superorder",
                "order",
                "suborder",
                "infraorder",
                "parvorder",
                "superfamily",
                "family",
                "subfamily",
                "tribe",
                "subtribe",
                "genus",
                "species_group",
                "species_subgroup",
                "species",
                "subspecies",
                "varietas",
                "strain",
                "no_rank_conf",
                "clade_conf",
                "superkingdom_conf",
                "kingdom_conf",
                "subkingdom_conf",
                "phylum_conf",
                "subphylum_conf",
                "superclass_conf",
                "class_conf",
                "subclass_conf",
                "infraclass_conf",
                "cohort_conf",
                "superorder_conf",
                "order_conf",
                "suborder_conf",
                "infraorder_conf",
                "parvorder_conf",
                "superfamily_conf",
                "family_conf",
                "subfamily_conf",
                "tribe_conf",
                "subtribe_conf",
                "genus_conf",
                "species_group_conf",
                "species_subgroup_conf",
                "species_conf",
                "subspecies_conf",
                "varietas_conf",
                "strain_conf")) %>% 
  left_join(kraken_db_norm %>% 
              select(contig,
                     lineage_1,
                     lineage_2,
                     lineage_3,
                     lineage_4,
                     lineage_5,
                     lineage_6,
                     lineage_7,
                     lineage_8), by = "contig") %>% 
  rename_with(~ paste0("kraken2_", .),
              c("lineage_1",
                "lineage_2",
                "lineage_3",
                "lineage_4",
                "lineage_5",
                "lineage_6",
                "lineage_7",
                "lineage_8"))

# Write this full database to a .txt file
write.table(full_community_db, "Bioinformatics/full_community_db.txt", quote = FALSE, row.names = FALSE, sep = "\t")
```

## Shotgun metagenomics of the BSFL gut microbiota

All *in silico* analyses were performed using shotgun metagenomic data of the midgut community of BSFL reared on standard diet (STD: 50 % (w/w) wheat bran; 20 % (w/w) alfalfa meal; 30 % (w/w) corn meal, mixed with water in 1:1 (w/w) ratio) obtained previously in @defilippis2023. Peritrophic matrix enclosed midguts were pooled into six independent samples of 15-30 midguts for DNA extraction, purification and library preparation with Illumina NovaSeq paired-end read sequencing. Host reads contamination was removed mapping reads to the *H. illucens* genome (NCBI Accession Number: PRJEB37575) by using BMTagger [@bmtagger_ncbi_2011] as reported previously [@defilippis2023]. The raw sequence reads generated previously were deposited in the Sequence Read Archive (SRA) of the NCBI under accession number PRJNA864640.

## Coassembly and read mapping

Shotgun metagenomic reads were quality-filtered using PRINSEQ 0.20.4 [@schmieder2011], to keep only reads with a Phred score $\geq$ 20 and length $\geq$ 75 bp previously [@defilippis2023]. FastQC 0.11.5 was used to assess read quality prior to coassembly [@andrews2010] and quality control results were compiled with MultiQC 1.16 into an interactive report for confirmation of sufficient read quality in all samples [@ewels2016] (**`r get_supp("Shotgun metagenomic read quality report")`**). Coassembly of contigs using merged forward and reverse reads from all six biological replicates was performed using MEGAHIT 1.2.9 with a minimum contig length of 1,000 bp, and the 'meta-large' preset enabled for high biodiversity samples [@li2015]. The resulting coassembly consisted of 112,981 contigs with a total length of 535,307,860 bp, with contigs ranging from 1,000 bp to 1,615,041 bp in size, and with an N50 value of 12,625 bp. The coassembly was indexed and reads for each sample were mapped to the coassembly with Bowtie2 2.3.4.1 [@langmead2012], with success rates across samples ranging from 89.0 to 91.7 %. Samtools 1.7 was used to sort and index the alignment file for downstream analysis [@li2009]. The 'jgi_summarize_bam_contig_depths' function of MetaBAT2 2.2.17 was used to generate average read depth values and standard deviation of read depth values for coassembly contigs independently for each sample [@kang2019].

## Contig taxonomic assignment

For contig-level taxonomic assignment, the coassembly was filtered to retain 112,892 contigs with a minimum contig length of 1,000 bp and a minimum average read depth of 2 based on MetaBAT2 2.2.17 depth summary results. Prodigal 2.6.3 was used to predict 579,379 open reading frames (ORFs) representing 475,501,254 bp out of the 535,209,308 bp of the filtered metagenomic contigs [@hyatt2010]. The resulting predicted amino acid sequences were then used for taxonomic classification of contigs with the Contig Annotation Tool (CAT) 6.0.1 due to the higher precision of CAT on bacterial benchmarks compared to other tools resulting from the ORF-based strategy that made it a suitable choice for the classification of longer contigs compared to shorter reads [@mirdita2021]. CAT leverages DIAMOND 2.1.12 for protein sequence alignment of ORFs against a pre-constructed NCBI non-redundant protein database obtained from the CAT repository [@buchfink2021; @vonmeijenfeldt2019]. For each predicted ORF, DIAMOND hits with a bitscore $\geq$ 50 % of the top-scoring hit were retained, and a weighted consensus across these hits was used to assign taxonomy to the contig. Of the 112,892 contigs retained after filtering, `r contigs_bacteria_num` (`r signif(contigs_bacteria_percentage, 3)` %) were assigned to the superkingdom bacteria by CAT. The relative abundance of bacterial contigs was similar across samples with an average of `r signif(bacteria_proportion_mean, 3)` % and a standard deviation of ± `r signif(bacteria_proportion_sd, 3)` %. Classification success and unique classifications at each taxonomic rank can be seen in (**Supplementary table \@ref(tab:bacteria-taxon-table)**). For additional context, Kraken2 2.1.6 was also used to predict taxonomic IDs of contigs using the pre-built July 2025 standard Kraken2 database and with a default confidence threshold of 0.2 [@wood2019]. TaxonKit 0.20.0 was used to obtain full taxonomic lineages for Kraken2 assigned taxonomic IDs in combination with the NCBI Taxonomy database [@sayers2022]. Due to the previously reported increased false positive rate associated with the used of Kraken2 [@wood2019], Kraken2 was used solely to provide alternative taxonomic classifications for readers.

## Contig binning and MAG taxonomic classification 

For contig binning into metagenome-assembled genomes (MAGs), the coassembly was filtered to retain 37,454 contigs with a minimum length of 2,000 bp, a minimum average read depth of 5 based on MetaBAT2 2.2.17 depth summary results, and a maximum average coverage variance of 50. Binning of contigs to MAGs was performed using MetaBAT2 2.2.17 [@kang2019], achieving binning of 94.0 % of the filtered contigs representing 307,506,758 bp across 110 bins. The completeness and contamination percentages of MAGs were then estimated with CheckM 1.2.4 [@parks2015]. Then, 48 'medium-quality MAGs' were retained by filtering MAGs to a minimum completeness of 50 % and a maximum contamination of 5 %. Taxonomic assignment of MAGs was performed with GTDB-Tk 2.3.2, referencing the GTDB release r207 v2 [@chaumeil2020]. The 'classify_wf' workflow was run with default parameters, skipping any dereplication or redundancy filtering with the 'skip_ani_screen' argument. MAGs were classified to species using FastANI 1.32 where average nucleotide identity (ANI) and alignment fraction (AF) to the closest reference genome met the GTDB thresholds of 95 % and 65 %, respectively [@jain2018]. Otherwise, taxonomic classification was assigned based on the alignment of MAGs to marker genes in the GTDB-Tk alignment pipeline and placement into the bacterial reference tree using pplacer 1.1 [@matsen2010]. GTDB-Tk taxonomic classification was successful for 47 MAGs.

## ARG, VF, and plasmid sequence prediction

Contig fasta sequences were analysed using the ABRicate software 1.0.1  [@seemann2025]. The Comprehensive Antibiotic Resistance Database (CARD) [@alcock2023], the Virulence Factor Database (VFDB) [@zhou2024], and PlasmidFinder [@carattoli2014] were utilised to search for ARG, VF, and plasmid sequences, respectively. All databases used with ABRicate were updated in January 2025. For additional ARG sequence metadata, "CARD data" 4.0.1 was downloaded from CARD and metadata extracted for identified ARG sequences using the predicted gene name. Predicted VF sequences were classified to 14 base categories according to the VFDB database based on VF product descriptions and gene names [@liu2022]. Unique plasmid accessions identified in the BSFL midgut microbiota were searched within the "ALL Plasmid Meta Data", "ALL Plasmid Antimicrobial Resistance Gene Meta Data", and "ALL Plasmid Virulent Factor Meta Data" databases available for download from the PlasmidScope database for predictions of plasmid mobility, ARG genes, and VF genes, respectively [@li2024].

## Isolation of antibiotic-resistant bacteria

BSF eggs were collected from a colony established in 2015 at the University of Insubria (Varese, Italy) and maintained as reported in @bruno2019. The eggs were laid on a Petri dish (9 x 1.5 cm) with STD for fly larvae rearing [@hogsette1992] and maintained in a humid chamber at 27 °C until hatching. Nipagin (methyl 4-hydroxybenzoate) was added to the diet to avoid mold growth. After emergence, BSF larvae were reared on this diet for 4 days and then placed in plastics boxes (16 x 16 x 9 cm and 12 x 12 x 8 cm) containing STD without nipagin. Actively feeding last instar larvae were collected, rinsed with tap water and then dried and washed with 70 % (v/v) ethanol. For each sample, the midguts from five larvae were isolated in sterile PBS in a sterile Petri dish (5.5 x 1.3 cm) using tweezers and scissors previously washed with 70 % (v/v) ethanol. The midgut content enclosed in the peritrophic matrix was separated from the epithelium and collected in 1.5 mL tubes [@bruno2025a]. Then, 10 µL of sterile PBS for each mg of midgut content  was added and resulting suspensions were homogenised with a plastic pestle. Concomitantly with the sampling of BSFL, approximately 100 mg of the rearing diet were also collected in 1.5 mL tubes, with 3 µL of sterile PBS added for each mg of diet and the resulting suspensions were crushed with a plastic pestle and vortexed. Five 10-fold dilutions of midgut content and diet samples were prepared in 1.5 mL tubes using sterile PBS. Luria Bertani (LB: 10 g/L tryptone, 5 g/L yeast extract, 10 g/L NaCl, 10 g/L agar) plates were prepared adding 8 different antibiotics (100 µg/mL ampicillin, 10 µg/mL tetracycline, 20 µg/mL chloramphenicol, 100 µg/mL streptomycin, 50 µg/mL kanamycin, 10 µg/mL gentamicin, 10 µg/mL ciprofloxacin and 10 µg/mL vancomycin). Antibiotic concentrations were chosen as common working concentration for laboratory plate preparation. Similar concentrations were used in previous works on antimicrobial resistance bacteria from environmental samples [@armalyte2019; @perelomov2023]. The last three 10-fold dilutions (1/1,000, 1/10,000, 1/100,000) of midgut content suspensions and diet sample suspensions were plated onto LB plates with antibiotics and were grown overnight at 37 °C. Resulting colonies were then picked and streaked onto another LB plate containing the corresponding antibiotic and grown overnight at 37 °C to isolate unique taxa. For each isolate plate, a single colony was picked with an inoculation loop and dispersed into a solution of 50 % (v/v) glycerol in liquid LB medium within cryovials which were subsequently stored at -80 °C. Samples of freshly prepared STD were also prepared in the same way and plated, however no growth was observed on antibiotic plates.

## MALDI-TOF species identification of resistant isolates

Resistant isolates were streaked on fresh plates of LB agar and incubated overnight at 37 °C for 16 hours. Identification was performed using a MALDI-TOF Biotyper Sirius Biotyper (Matrix-Assisted Laser Desorption/Ionization Time-Of-Flight; Bruker, USA), according to the manufacturer's standard protocol. Briefly, 1 µL of 70 % (v/v) formic acid solution in water and 1 µL of α-cyano-4-hydrocinnamic acid matrix solution (Bruker, USA) were added directly onto the colonies and allowed to air dry. Spectra were analyzed with Bruker Biotyper Compass HT RUO software 5.1.3 with the MBT Compass Library 12.0. Calibration was performed for each target plate using Bruker Bacterial Test Standard (BTS). To ensure confidence in the identification of bacterial taxa from samples, a log score of spectral similarity above the 1.7 threshold was considered.

## Data analysis

Read depths for each contig were normalised to relative abundances based on the total read depth per sample. To investigate the bacterial community structure, contigs were filtered to those predicted as "Bacteria" by CAT and Kraken2 separately and the read depths re-normalised for these subsets of contigs to obtain 'relative bacterial abundances'. All data analysis and visualisation was performed in RStudio 2025.09.0+387 "Cucumberleaf Sunflower" release built on R 4.5.1 [@positteam2025]. Phylum-level community structures for the entire bacterial community, the counts of VFs belonging to the VF product classes, and MAG phylum-level classification numbers were plotted with the 'ggplot2' package [@wickham2016]. Heatmaps of antibiotic resistance mechanism against drug class, phylum level-contributions to ARGs and VFs of bacterial contigs, and genus-level contributions of bacterial MAGs to ARGs and VFs were plotted with the 'pheatmap' package [@kolde2025]. Plasmid mobility and contributions to antibiotic resistance, and antibiotic resistance of isolates were visualised woth alluvial plots using the 'ggalluvial' package [@brunson2020]. To visualise MAG-gene associations for ARG and VF genes, an edge-table was generated from links between MAGs and genes based on contig IDs of predicted genes and binned contigs. The 'igraph' package was used to generate a graph object distinguishing Taxon, ARG, and VF nodes based on size and colour, and edges were distinguished based on single taxon or multiple taxon connections [@csardi2025]. The network was plotted using the 'ggraph' package utilising a Fruchterman-Reingold layout [@pedersen2024]. Multi-panel figures of ggplot objects were combined with the 'patchwork' package [@pedersen2025]. Multiple pheatmap plots were combined with the 'gridExtra' package [@auguie2017]. Combinations of ggplot and pheatmap objects for figures were made by saving objects as individual image files and combining with the 'magick' package [@ooms2025].

Interactive kingdom and phylum bacterial relative abundance plots were created using the 'ggplot' and 'plotly' packages [@wickham2016; @sievert2020]. Interactive sankey diagrams of the bacterial relative abundance ordered hierarchically from Class to Order to Family taxonomic classifications were constructed using the 'sankeyNetwork' function of the 'networkD3' package, and were rendered with customisation using the 'htmlwidgets' package [@allaire2025; @vaidyanathan2023]. Interactive heatmaps of genus- and species-level community structures across samples were visualised using the 'heatmaply' package [@galili2017]. Interactive community figures for both CAT- and Kraken2-classified bacterial contig communities can be seen in **`r get_supp("Interactive taxonomic community structures")`**. The interactive sunburst plot depicting the hierarchical structure of MAG taxonomic classifications was plotted using the 'plotly' package [@sievert2020]. MAG genus and species classifications by GTDB-Tk alongside FastANI and pplacer reference genome accessions were presented in table format using the 'kableExtra' package [@zhu2024]. The interactive MAG taxonomic classifications are visible in **`r get_supp("Interactive MAG taxonomic classification")`**.

The submitted manuscript was generated in Word .docx format and the interactive figures were rendered in .html supplementary files from 'rmarkdown' using the 'knitr' and 'bookdown' packages [@xie2020; @xie2014; @xie2016]. Tables in the .docx manuscript were generated using the 'flextable' package [@gohel2025]. The .Rmd file used to generate this manuscript containing all text and code for analysis and plotting is available in **`r get_supp("R Markdown manuscript file")`**.

\newpage

# Results

## Bacterial community structure of the BSFL midgut microbiota

```{r community-plot, fig.cap="**Bacterial community structure of the BSFL midgut.** (**A**) The relative bacterial abundances of bacterial phyla for each sample. Phyla with relative bacterial abundances < 1 % were grouped into Other. (**B**) Genera with average relative bacterial abundances > 0.1 % across the six samples. Bacterial relative abundance values for genera are presented within cells for each sample, with genera ordered in descending mean relative bacterial abundance. Columns represent individual samples.", out.width="100%"}
knitr::include_graphics("plots/combined_community_plot.png")
```

The BSFL bacterial midgut community was revealed to be similar to previous assessments based on 16S rRNA gene amplicon sequencing or marker-gene based assignment of shotgun metagenomic contigs for larvae reared on the standard diet [@defilippis2023]. Bacteroidota was the most abundant phylum at `r signif(phylum_sum_stats["Bacteroidota"], 3)` % on average (**Figure \@ref(fig:community-plot)**), while Pseudomonadota contributed on average to `r signif(phylum_sum_stats["Pseudomonadota"], 3)` % of the relative bacterial abundance, less than Bacteroidota despite the larger number of unique contigs assigned to the Pseudomonadota phylum (**Supplementary table \@ref(tab:sup-bacteria-phylum-table)**). Actinomycetota and Bacillota phyla constituted on average `r signif(phylum_sum_stats["Actinomycetota"], 3)` % and `r signif(phylum_sum_stats["Bacillota"], 3)` %, respectively. The remaining classified phyla together contributed only `r signif(phylum_sum_stats["Other"], 3)` %. The remaining `r signif(phylum_sum_stats["Unclassified"], 3)` % of the relative bacterial abundance of the community structure was derived from contigs that were unclassified at phylum level by CAT. About `r signif(top_genera$mean_abundance[top_genera$Genus == "Unclassified"], 3)` % of the relative abundance of bacterial contigs was attributed to taxa unclassified at the genus level by CAT. Contigs unclassified at genus level were distributed across `r length(genus_unclassified$Phylum)` phyla including Actinomycetota, Bacillota, Bacteroidota, and Pseudomonadota. The *Sphingobacterium* genus was observed with an average bacterial relative abundance of `r signif(top_genera$mean_abundance[top_genera$Genus == "Sphingobacterium"], 3)` %, compared to the second most abundant taxa classified to the genus-level of *Sinirhodobacter* at `r signif(top_genera$mean_abundance[top_genera$Genus == "Sinirhodobacter"], 3)` %. The community structure from kingdom to species taxonomic ranks can be explored interactively in more detail for CAT and Kraken2 predicted communities in **`r get_supp("Interactive taxonomic community structures")`**.

\newpage

## Antibiotic resistance and virulence in the BSFL midgut microbiota

```{r CAT-arg-sum}
# Filter to contigs with at least 1 predicted antibiotic resistance gene and pivot longer
CAT_arg_contigs <- contig_db_norm %>% 
  filter(!is.na(arg1_start)) %>% 
  pivot_longer(
    cols = matches("^arg\\d+_"),
    names_to = c("arg_id", ".value"),
    names_pattern = "(arg\\d+)_(.*)"
  ) %>% 
  filter(!is.na(start))

# Count the number of rows which is equivalent to the number of predicted ARGs for all contigs
CAT_total_args <- nrow(CAT_arg_contigs)

# Filter to the Bacterial superkingdom
CAT_bacteria_arg <- CAT_arg_contigs %>% 
  filter(superkingdom == "Bacteria")

# Count the number of ARGs on contigs predicted to be bacterial by CAT
CAT_bacteria_arg_count <- nrow(CAT_bacteria_arg)
# Count the number of unique contigs
CAT_bacteria_arg_contigs <- length(unique(CAT_bacteria_arg$contig))
# Calculate the percentage of contigs with ARGs predicted to be bacterial
CAT_bacteria_arg_contigs_percentage <- (CAT_bacteria_arg_contigs / length(unique(CAT_arg_contigs$contig))) * 100

# Lastly sum the average across the relative abundance columns to calculate the total relative abundance of ARG-carrying contigs
total_arg_relabun <- contig_db_norm %>% 
  filter(!is.na(arg1_start)) %>% 
  rowwise() %>% 
  mutate(mean_rel_abun = mean(c_across(starts_with("norm_")), na.rm = TRUE)) %>% 
  ungroup() %>% 
  { sum(.$mean_rel_abun) }

# Now sum the average across the relative abundance columns only for contigs assigned as Bacterial by CAT
CAT_bacteria_arg_relabun <- contig_db_norm %>% 
  filter(superkingdom == "Bacteria") %>% 
  filter(!is.na(arg1_start)) %>% 
  rowwise() %>% 
  mutate(mean_rel_abun = mean(c_across(starts_with("norm_")), na.rm = TRUE)) %>% 
  ungroup() %>% 
  { sum(.$mean_rel_abun) }
```

```{r card-meta}
# Read in the card Antibiotic Resistance Ontology (ARO) index which contains the metadata for each ARG prediction
card_meta <- read.delim("Bioinformatics/abricate/aro_index.tsv", header = TRUE)

# Make a map of shorter names for the antibiotics
drug_code_map <- c(
  "acridine_dye" = "AD",
  "aminocoumarin" = "AC",
  "aminoglycoside" = "AG",
  "bicyclomycin" = "BC",
  "carbapenem" = "CB",
  "cephalosporin" = "CS",
  "cephamycin" = "CY",
  "diaminopyrimidine" = "DP",
  "fluoroquinolone" = "FQ",
  "fosfomycin" = "FO",
  "glycylcycline" = "GC",
  "glycopeptide" = "GP",
  "lincosamide" = "LC",
  "macrolide" = "ML",
  "monobactam" = "MB",
  "mupirocin" = "MP",
  "nitroimidazole" = "NI",
  "nitrofuran" = "NF",
  "peptide" = "PT",
  "penam" = "PN",
  "penem" = "PM",
  "phenicol" = "PH",
  "rifamycin" = "RF",
  "sulfonamide" = "SM",
  "tetracycline" = "TC",
  "triclosan" = "TR"
)

# Make a function for matching the name of antibiotic with a shorter code allowing fuzzy matching (not-exact matching)
convert_to_codes_fuzzy <- function(class_string, max_dist = 2) {
  if (is.na(class_string) || class_string == "") return(NA)
  
  class_list <- unlist(strsplit(class_string, ";"))
  class_list <- trimws(class_list)
  
  code_list <- sapply(class_list, function(x) {
    # Compute string distance to all known drug classes
    distances <- stringdist::stringdist(tolower(x), tolower(names(drug_code_map)))
    min_dist <- min(distances)
    
    # Accept only if the distance is below threshold
    if (min_dist <= max_dist) {
      closest <- names(drug_code_map)[which.min(distances)]
      return(drug_code_map[closest])
    } else {
      return("UNK")  # Unknown class if too far
    }
  })
  
  return(paste(unique(code_list), collapse = ";"))
}

# Make a table of the shortened antibiotic codes
drug_class_legend <- data.frame(
  Code = unname(drug_code_map),
  Drug_Class = names(drug_code_map)
) %>% 
  rename(Class = Drug_Class) %>% 
  mutate(Class = str_to_title(Class),
         Class = gsub("Acridine_dye", "Acridine dye", Class))
```

```{r arg-classification-make, eval=FALSE}
# I want to see for each arg it's classifications from CAT and Kraken2, and then I want to merge on the read depths across samples for a supplementary .csv file
arg_classifications <- CAT_arg_contigs %>% 
  select(c("contig", "length", "superkingdom", "kingdom", "phylum", "class", "order", "family", "genus", "species", "arg_id", "start", "end", "strand", "gene", "coverage", "coverage_map", "gaps", "coverage_percentage", "pident", "accession", "product", "resistance")) %>% 
  rename(CAT_superkingdom = superkingdom, CAT_kingdom = kingdom, CAT_phylum = phylum, CAT_class = class, CAT_order = order, CAT_family = family, CAT_genus = genus, CAT_species = species) %>% 
  left_join(kraken_arg_contigs %>% 
              select(c("contig", "arg_id", "lineage_1", "lineage_2", "lineage_3", "lineage_4", "lineage_5", "lineage_6", "lineage_7", "lineage_8")) %>% 
              rename(kraken2_superkingdom = lineage_1, kraken2_kingdom = lineage_2, kraken2_phylum = lineage_3, kraken2_class = lineage_4, kraken2_order = lineage_5, kraken2_family = lineage_6, kraken2_genus = lineage_7, kraken2_species = lineage_8), by = c("contig", "arg_id")) %>% 
  mutate(kraken2_superkingdom = case_when(
    kraken2_kingdom %in% c("Pseudomonadati", "Bacillati", "unclassified Bacteria kingdom") ~ "Bacteria",
    TRUE ~ "Unclassified"
  ),
  CAT_superkingdom = ifelse(is.na(CAT_superkingdom), "Unclassified", CAT_superkingdom)) %>%
  mutate(across(kraken2_kingdom:kraken2_species, ~ ifelse(grepl("^unclassified", .), NA, .))) %>% 
  left_join(card_meta %>% rename(gene = ARO.Name))

# Convert antibiotic classes to the shorter codes
arg_classifications$drug_code <- sapply(arg_classifications$resistance, convert_to_codes_fuzzy)

# Merge the read depth to the dataframe for a supplementary table
arg_classifications <- arg_classifications %>% 
  left_join(contig_db_norm %>% select(c("contig", "norm_STD_1_Avg", "norm_STD_2_Avg", "norm_STD_4_Avg", "norm_STD_5_Avg", "norm_STD_7_Avg", "norm_STD_8_Avg"))) %>% 
  rename(STD_1_RelAbun = norm_STD_1_Avg, STD_2_RelAbun = norm_STD_2_Avg, STD_4_RelAbun = norm_STD_4_Avg, STD_5_RelAbun = norm_STD_5_Avg, STD_7_RelAbun = norm_STD_7_Avg, STD_8_RelAbun = norm_STD_8_Avg, ARG_start = start, ARG_end = end, ARG_strand = strand, ARG_gene = gene, ARG_coverage = coverage, ARG_coverage_map = coverage_map, ARG_gaps = gaps, ARG_coverage_percentage = coverage_percentage, ARG_pident = pident, ARG_accession = accession, ARG_product = product, ARG_resistance = resistance, ARO_accession = ARO.Accession, ARO_cv_term = CVTERM.ID, ARO_model_seqID = Model.Sequence.ID, ARO_model_ID = Model.ID, ARO_model_name = Model.Name, ARO_protein_accession = Protein.Accession, ARO_DNA_accession = DNA.Accession, ARO_gene_family = AMR.Gene.Family, ARO_drug_class = Drug.Class, ARO_resistance_mechanism = Resistance.Mechanism, ARO_drug_code = drug_code) %>% 
  select(c("contig", "length", "ARG_start", "ARG_end", "ARG_strand", "ARG_gene", "ARG_coverage", "ARG_coverage_map", "ARG_gaps", "ARG_coverage_percentage", "ARG_pident", "ARG_accession", "ARG_product", "ARG_resistance", "ARO_accession", "ARO_cv_term", "ARO_model_seqID", "ARO_model_ID", "ARO_model_name", "ARO_protein_accession", "ARO_DNA_accession", "ARO_gene_family", "ARO_drug_class", "ARO_resistance_mechanism", "ARO_drug_code", "CAT_superkingdom", "CAT_kingdom", "CAT_phylum", "CAT_class", "CAT_order", "CAT_family", "CAT_genus", "CAT_species", "kraken2_superkingdom", "kraken2_kingdom", "kraken2_phylum", "kraken2_class", "kraken2_order", "kraken2_family", "kraken2_genus", "kraken2_species", "STD_1_RelAbun", "STD_2_RelAbun", "STD_4_RelAbun", "STD_5_RelAbun", "STD_7_RelAbun", "STD_8_RelAbun"))
```

```{r arg_db_sup, eval=FALSE}
# Make a .csv file with a leading line explaining each column for a supplementary file
write.csv(arg_classifications, "Bioinformatics/abricate/ARG_db.csv", row.names = FALSE)
```

```{r arg-classification-read}
# Read in the arg_classification database
arg_classifications <- read.csv("Bioinformatics/abricate/ARG_db.csv", header = TRUE)
```

```{r CAT-vf-sum}
# Filter to contigs with at least 1 predicted virulence factor and pivot longer
CAT_vf_contigs <- contig_db_norm %>% 
  filter(!is.na(vf1_start)) %>% 
  pivot_longer(
    cols = matches("^vf\\d+_"),
    names_to = c("vf_id", ".value"),
    names_pattern = "(vf\\d+)_(.*)"
  ) %>% 
  filter(!is.na(start))

# Count the number of rows which is equivalent to the number of predicted VFs for all contigs
CAT_total_vfs <- nrow(CAT_vf_contigs)

# Filter to the Bacterial superkingdom
CAT_bacteria_vf <- CAT_vf_contigs %>% 
  filter(superkingdom == "Bacteria")

# Count the number of VFs on contigs predicted to be bacterial by CAT
CAT_bacteria_vf_count <- nrow(CAT_bacteria_vf)
# Count the number of unique contigs
CAT_bacteria_vf_contigs <- length(unique(CAT_bacteria_vf$contig))
# Calculate the percentage of contigs with VFs predicted to be bacterial
CAT_bacteria_vf_contigs_percentage <- (CAT_bacteria_vf_contigs / length(unique(CAT_vf_contigs$contig))) * 100

# Lastly sum the average across the relative abundance columns to calculate the total relative abundance of VF-carrying contigs
total_vf_relabun <- contig_db_norm %>% 
  filter(!is.na(vf1_start)) %>% 
  rowwise() %>% 
  mutate(mean_rel_abun = mean(c_across(starts_with("norm_")), na.rm = TRUE)) %>% 
  ungroup() %>% 
  { sum(.$mean_rel_abun) }

# Now sum the average across the relative abundance columns only for contigs assigned as Bacterial by CAT
CAT_bacteria_vf_relabun <- contig_db_norm %>% 
  filter(superkingdom == "Bacteria") %>% 
  filter(!is.na(vf1_start)) %>% 
  rowwise() %>% 
  mutate(mean_rel_abun = mean(c_across(starts_with("norm_")), na.rm = TRUE)) %>% 
  ungroup() %>% 
  { sum(.$mean_rel_abun) }
```

```{r vf-classification-make, eval=FALSE}
# I want to see for each vf it's classifications from CAT and Kraken2, and then I want to merge on the read depths across samples for a supplementary .csv file
vf_classifications <- CAT_vf_contigs %>% 
  select(c("contig", "length", "superkingdom", "kingdom", "phylum", "class", "order", "family", "genus", "species", "vf_id", "start", "end", "strand", "gene", "coverage", "coverage_map", "gaps", "coverage_percentage", "pident", "accession", "product")) %>% 
  rename(CAT_superkingdom = superkingdom, CAT_kingdom = kingdom, CAT_phylum = phylum, CAT_class = class, CAT_order = order, CAT_family = family, CAT_genus = genus, CAT_species = species) %>% 
  left_join(kraken_vf_contigs %>% 
              select(c("contig", "vf_id", "lineage_1", "lineage_2", "lineage_3", "lineage_4", "lineage_5", "lineage_6", "lineage_7", "lineage_8")) %>% 
              rename(kraken2_superkingdom = lineage_1, kraken2_kingdom = lineage_2, kraken2_phylum = lineage_3, kraken2_class = lineage_4, kraken2_order = lineage_5, kraken2_family = lineage_6, kraken2_genus = lineage_7, kraken2_species = lineage_8), by = c("contig", "vf_id")) %>% 
  mutate(kraken2_superkingdom = case_when(
    kraken2_kingdom %in% c("Pseudomonadati", "Bacillati", "unclassified Bacteria kingdom") ~ "Bacteria",
    TRUE ~ "Unclassified"
  ),
  CAT_superkingdom = ifelse(is.na(CAT_superkingdom), "Unclassified", CAT_superkingdom)) %>%
  mutate(across(kraken2_kingdom:kraken2_species, ~ ifelse(grepl("^unclassified", .), NA, .))) %>% 
  left_join(contig_db_norm %>% select(c("contig", "norm_STD_1_Avg", "norm_STD_2_Avg", "norm_STD_4_Avg", "norm_STD_5_Avg", "norm_STD_7_Avg", "norm_STD_8_Avg")), by = "contig") %>% 
  rename(STD_1_RelAbun = norm_STD_1_Avg, STD_2_RelAbun = norm_STD_2_Avg, STD_4_RelAbun = norm_STD_4_Avg, STD_5_RelAbun = norm_STD_5_Avg, STD_7_RelAbun = norm_STD_7_Avg, STD_8_RelAbun = norm_STD_8_Avg, VF_start = start, VF_end = end, VF_strand = strand, VF_gene = gene, VF_coverage = coverage, VF_coverage_map = coverage_map, VF_gaps = gaps, VF_coverage_percentage = coverage_percentage, VF_pident = pident, VF_accession = accession, VF_product = product) %>% 
  select(c("contig", "length", "VF_start", "VF_end", "VF_strand", "VF_gene", "VF_coverage", "VF_coverage_map", "VF_gaps", "VF_coverage_percentage", "VF_pident", "VF_accession", "VF_product", "CAT_superkingdom", "CAT_kingdom", "CAT_phylum", "CAT_class", "CAT_order", "CAT_family", "CAT_genus", "CAT_species", "kraken2_superkingdom", "kraken2_kingdom", "kraken2_phylum", "kraken2_class", "kraken2_order", "kraken2_family", "kraken2_genus", "kraken2_species", "STD_1_RelAbun", "STD_2_RelAbun", "STD_4_RelAbun", "STD_5_RelAbun", "STD_7_RelAbun", "STD_8_RelAbun"))
```

```{r vf_db_sup, eval=FALSE}
# Make a .csv file with a leading line explaining each column for a supplementary file
write.csv(vf_classifications, "Bioinformatics/abricate/VF_db.csv", row.names = FALSE)
```

``` {r vf-classification-read}
# Read in the VF classification db
vf_classifications <- read.csv("Bioinformatics/abricate/VF_db.csv", header = TRUE)
```

```{r ARG-class-count}
# Repeat for the genes predicted to be bacteria by CAT
CAT_ARG_class_counts <- arg_classifications %>%
  filter(CAT_superkingdom == "Bacteria") %>% 
  separate_rows(ARO_drug_code, sep = ";") %>%  # each code becomes a row
  group_by(ARO_drug_code) %>%
  summarise(Count = n()) %>%
  arrange(desc(Count))

# Count the number of classes
ARG_class_num <- nrow(CAT_ARG_class_counts)

# Sum the total number of hits to confer resistance to these classes from CAT classified bacterial contigs
CAT_ARG_class_sum <- sum(CAT_ARG_class_counts$Count)

# Merge drug codes
antibiotic_class_count <- CAT_ARG_class_counts %>% 
  rename(Code = ARO_drug_code) %>% 
  left_join(drug_class_legend, by = "Code") %>% 
  select(c("Class", "Code", "Count"))

# Make a list of the antibiotic names with codes in brackets that have over 10 counts of resistance
antibiotic_class_list <- antibiotic_class_count %>% 
  filter(Count > 10) %>% 
  filter(Code != "FQ") %>% 
  mutate(Name = paste0(Class, " (", Code, ")"))

antibiotic_class_list_single <- antibiotic_class_count %>% 
  filter(Count == 1) %>%  
  mutate(Name = paste0(Class, " (", Code, ")"))
```

```{r mechanism}
# Isolate just the contigs, arg_ids, and their resistance mechanisms
CAT_mechanism <- arg_classifications %>% 
  filter(CAT_superkingdom == "Bacteria") %>% 
  select(c(contig, ARO_resistance_mechanism, STD_1_RelAbun, STD_2_RelAbun, STD_4_RelAbun, STD_5_RelAbun, STD_7_RelAbun, STD_8_RelAbun)) %>% 
  mutate(ARO_resistance_mechanism = ifelse(is.na(ARO_resistance_mechanism), "Unclassified", ARO_resistance_mechanism)) %>% 
  rename(Mechanism = ARO_resistance_mechanism) %>% 
  mutate(Mechanism = gsub("antibiotic efflux;antibiotic target alteration", "Efflux", Mechanism),
         Mechanism = gsub("antibiotic efflux;reduced permeability to antibiotic", "Efflux", Mechanism),
         Mechanism = gsub("antibiotic inactivation", "Inactivation", Mechanism),
         Mechanism = gsub("antibiotic target alteration", "Target alteration", Mechanism),
         Mechanism = gsub("antibiotic target protection", "Target protection", Mechanism),
         Mechanism = gsub("antibiotic target replacement", "Target replacement", Mechanism),
         Mechanism = gsub("antibiotic efflux", "Efflux", Mechanism))

# Count the number of rows e.g. ARGs for each type of mechanism
CAT_mechanism_count <- CAT_mechanism %>% 
  group_by(Mechanism) %>% 
  summarise("Count" = length(Mechanism))

# Merge together and add a new column to signify it's all contigs
mechanism_df <- CAT_mechanism_count %>% 
  mutate(across(everything(), ~replace_na(.x, 0)))
```

```{r class-vs-mechanism}
# CAT classified bacterial contigs
CAT_bacteria_mech_drug_df <- arg_classifications %>%
  filter(CAT_superkingdom == "Bacteria") %>% 
  select(ARO_drug_code, ARO_resistance_mechanism) %>% 
  mutate(ARO_resistance_mechanism = ifelse(is.na(ARO_resistance_mechanism), "Unclassified", ARO_resistance_mechanism)) %>% 
  separate_rows(ARO_drug_code, sep = ";") %>%
  mutate(
    ARO_resistance_mechanism = ifelse(
      str_detect(tolower(ARO_resistance_mechanism), "antibiotic efflux"),
      "antibiotic efflux",  # Merge all efflux types
      ARO_resistance_mechanism
    )
  ) %>%
  group_by(ARO_resistance_mechanism, ARO_drug_code) %>%
  summarise(Count = n(), .groups = "drop") %>%
  pivot_wider(names_from = ARO_resistance_mechanism, values_from = Count, values_fill = 0) %>% 
  left_join(antibiotic_class_count %>% select(c(Code, Class)) %>% rename(ARO_drug_code = Code), by = "ARO_drug_code") %>% 
  select(-ARO_drug_code) %>% 
  column_to_rownames("Class") %>% 
  rename(Efflux = "antibiotic efflux", Inactivation = "antibiotic inactivation", "Target alteration" = "antibiotic target alteration", "Target protection" = "antibiotic target protection") %>% 
  mutate(Target_replacement = 0) %>% 
  rename("Target replacement" = Target_replacement) %>% 
  select(c("Efflux", "Inactivation", "Target alteration", "Target protection", "Target replacement", "Unclassified")) 

CAT_bacteria_mech_drug_df_sum <- as.data.frame(colSums(CAT_bacteria_mech_drug_df)) %>% 
  rename(total_count = "colSums(CAT_bacteria_mech_drug_df)") %>% 
  rownames_to_column("mechanism")

CAT_bacteria_mech_class_df_sum <- as.data.frame(colSums(CAT_bacteria_mech_drug_df > 0)) %>% 
  rename(total_classes = "colSums(CAT_bacteria_mech_drug_df > 0)") %>% 
  rownames_to_column("mechanism")

CAT_bacteria_mech_drug_mat <- CAT_bacteria_mech_drug_df %>% 
  as.matrix() %>% 
  log1p()

# define common breaks across all matrices
drug_vs_mech_breaks <- seq(min(CAT_bacteria_mech_drug_mat), max(CAT_bacteria_mech_drug_mat), length.out = 101)
# define a common color palette
drug_vs_mech_colors <- colorRampPalette(c("white", "#4575B4"))(100)

# Convert the original count dataframe to a numeric matrix
drug_vs_mech_labels <- as.matrix(CAT_bacteria_mech_drug_df)
```

```{r vf-product-parse}
# I want to reduce all the unique products from VFs into some more succint classes. I can do this sequentially by first reducing anything that is a Type I, II, III etc. secretion system into these products, and then see what I am left with and move onto the next pattern I can spot.
vf_classifications_tidy <- vf_classifications %>%
  mutate(
    VF_product_class = if_else(
      str_detect(VF_product, regex("type III secretion system", ignore_case = TRUE)),
      "Effector delivery system",
      VF_product
    )
  ) %>%
  mutate(
    VF_product_class = if_else(
      str_detect(VF_product_class, regex("type IV secret", ignore_case = TRUE)),
      "Effector delivery system",
      VF_product_class
    )
  ) %>%
  mutate(
    VF_product_class = if_else(
      str_detect(VF_product_class, regex("type VI secretion", ignore_case = TRUE)),
      "Effector delivery system",
      VF_product_class
    )
  ) %>%
  mutate(
    VF_product_class = if_else(
      str_detect(VF_product_class, regex("type six secretion", ignore_case = TRUE)),
      "Effector delivery system",
      VF_product_class
    )
  ) %>%
  mutate(
    VF_product_class = if_else(
      str_detect(VF_product_class, regex(" secretion ", ignore_case = TRUE)),
      "Effector delivery system",
      VF_product_class
    )
  ) %>%
  mutate(
    VF_product_class = if_else(
      str_detect(VF_product_class, regex("flagellar", ignore_case = TRUE)),
      "Motility",
      VF_product_class
    )
  ) %>% 
  mutate(
    VF_product_class = if_else(
      str_detect(VF_product_class, regex("flagellum", ignore_case = TRUE)),
      "Motility",
      VF_product_class
    )
  ) %>% 
  mutate(
    VF_product_class = if_else(
      str_detect(VF_product_class, regex("curli", ignore_case = TRUE)),
      "Adherence",
      VF_product_class
    )
  ) %>% 
    mutate(
    VF_product_class = if_else(
      str_detect(VF_product_class, regex("twitching motility protein", ignore_case = TRUE)),
      "Motility",
      VF_product_class
    )
  ) %>% 
  mutate(
    VF_product_class = if_else(
      str_detect(VF_product_class, regex("pilus", ignore_case = TRUE)),
      "Motility",
      VF_product_class
    )
  ) %>% 
  mutate(
    VF_product_class = if_else(
      str_detect(VF_product_class, regex("pili ", ignore_case = TRUE)),
      "Motility",
      VF_product_class
    )
  ) %>% 
  mutate(
    VF_product_class = if_else(
      str_detect(VF_product_class, regex("polymerized tip adhesin", ignore_case = TRUE)),
      "Motility",
      VF_product_class
    )
  ) %>%
  mutate(
    VF_product_class = if_else(
      str_detect(VF_product_class, regex("chemotaxis", ignore_case = TRUE)),
      "Motility",
      VF_product_class
    )
  ) %>% 
  mutate(
    VF_product_class = if_else(
      str_detect(VF_product_class, regex("regulator protein EcpR", ignore_case = TRUE)),
      "Regulation",
      VF_product_class
    )
  ) %>% 
  mutate(
    VF_product_class = if_else(
      str_detect(VF_product_class, regex("fimbrial", ignore_case = TRUE)),
      "Adherence",
      VF_product_class
    )
  ) %>% 
  mutate(
    VF_product_class = if_else(
      str_detect(VF_product_class, regex("outer membrane", ignore_case = TRUE)),
      "Biofilm",
      VF_product_class
    )
  ) %>% 
  mutate(
    VF_product_class = if_else(
      str_detect(VF_product_class, regex("B-band O-antigen", ignore_case = TRUE)),
      "Adherence",
      VF_product_class
    )
  ) %>%
  mutate(
    VF_product_class = if_else(
      str_detect(VF_product_class, regex("Collagen binding", ignore_case = TRUE)),
      "Adherence",
      VF_product_class
    )
  ) %>% 
  mutate(
    VF_product_class = if_else(
      str_detect(VF_product_class, regex("serum resistance protein", ignore_case = TRUE)),
      "Immune modulation",
      VF_product_class
    )
  ) %>% 
  mutate(
    VF_product_class = if_else(
      str_detect(VF_product_class, regex("lipoprotein", ignore_case = TRUE)),
      "Effector delivery system",
      VF_product_class
    )
  ) %>% 
  mutate(
    VF_product_class = if_else(
      str_detect(VF_product_class, regex("fibrinogen binding protein", ignore_case = TRUE)),
      "Adherence",
      VF_product_class
    )
  ) %>% 
  mutate(
    VF_product_class = if_else(
      str_detect(VF_product_class, regex("exotoxin ", ignore_case = TRUE)),
      "Exotoxin",
      VF_product_class
    )
  ) %>% 
  mutate(
    VF_product_class = if_else(
      str_detect(VF_product_class, regex("enterotoxin ", ignore_case = TRUE)),
      "Exotoxin",
      VF_product_class
    )
  ) %>% 
  mutate(
    VF_product_class = if_else(
      str_detect(VF_product_class, regex("transcriptional ", ignore_case = TRUE)),
      "Regulation",
      VF_product_class
    )
  ) %>% 
  mutate(
    VF_product_class = if_else(
      str_detect(VF_product_class, regex("sigma-70", ignore_case = TRUE)),
      "Regulation",
      VF_product_class
    )
  ) %>% 
  mutate(
    VF_product_class = if_else(
      str_detect(VF_product_class, regex("receptor", ignore_case = TRUE)),
      "Nutritional/Metabolic factor",
      VF_product_class
    )
  ) %>% 
  mutate(
    VF_product_class = if_else(
      str_detect(VF_product_class, regex("ABC", ignore_case = TRUE)),
      "Nutritional/Metabolic factor",
      VF_product_class
    )
  ) %>% 
  mutate(
    VF_product_class = if_else(
      str_detect(VF_product_class, regex("envelope", ignore_case = TRUE)),
      "Regulation",
      VF_product_class
    )
  ) %>% 
  mutate(
    VF_product_class = if_else(
      str_detect(VF_product_class, regex("signal", ignore_case = TRUE)),
      "Regulation",
      VF_product_class
    )
  ) %>% 
  mutate(
    VF_product_class = if_else(
      str_detect(VF_product_class, regex("two-component response regulator", ignore_case = TRUE)),
      "Regulation",
      VF_product_class
    )
  ) %>% 
  mutate(
    VF_product_class = if_else(
      str_detect(VF_product_class, regex("Rab2", ignore_case = TRUE)),
      "Immune modulation",
      VF_product_class
    )
  ) %>% 
  mutate(
    VF_product_class = if_else(
      str_detect(VF_product_class, regex("protease", ignore_case = TRUE)),
      "Exoenzyme",
      VF_product_class
    )
  ) %>% 
  mutate(
    VF_product_class = if_else(
      str_detect(VF_product_class, regex("lyase", ignore_case = TRUE)),
      "Nutritional/Metabolic factor",
      VF_product_class
    )
  ) %>% 
  mutate(
    VF_product_class = if_else(
      str_detect(VF_product_class, regex("dipeptidase", ignore_case = TRUE)),
      "Nutritional/Metabolic factor",
      VF_product_class
    )
  ) %>%
  mutate(
    VF_product_class = if_else(
      str_detect(VF_product_class, regex("phenazine", ignore_case = TRUE)),
      "Nutritional/Metabolic factor",
      VF_product_class
    )
  ) %>% 
  mutate(
    VF_product_class = if_else(
      str_detect(VF_product_class, regex("alginate biosynthesis protein Alg8", ignore_case = TRUE)),
      "Biofilm",
      VF_product_class
    )
  ) %>% 
  mutate(
    VF_product_class = if_else(
      str_detect(VF_product_class, regex("alginate biosynthesis", ignore_case = TRUE)),
      "Nutritional/Metabolic factor",
      VF_product_class
    )
  ) %>% 
  mutate(
    VF_product_class = if_else(
      str_detect(VF_product_class, regex("salicylate biosynthesis", ignore_case = TRUE)),
      "Nutritional/Metabolic factor",
      VF_product_class
    )
  ) %>% 
  mutate(
    VF_product_class = if_else(
      str_detect(VF_product_class, regex("pyoverdine biosynthesis", ignore_case = TRUE)),
      "Nutritional/Metabolic factor",
      VF_product_class
    )
  ) %>% 
  mutate(
    VF_product_class = if_else(
      str_detect(VF_product_class, regex("lipid A biosynthesis lauroyl", ignore_case = TRUE)),
      "Biofilm",
      VF_product_class
    )
  ) %>%
  mutate(
    VF_product_class = if_else(
      str_detect(VF_product_class, regex("alginate biosynthesis protein AlgZ/FimS", ignore_case = TRUE)),
      "Regulation",
      VF_product_class
    )
  ) %>%
  mutate(
    VF_product_class = if_else(
      str_detect(VF_product_class, regex("Lipopolysaccharide core", ignore_case = TRUE)),
      "Biofilm",
      VF_product_class
    )
  ) %>%
  mutate(
    VF_product_class = if_else(
      str_detect(VF_product_class, regex("negative regulator for alginate biosynthesis", ignore_case = TRUE)),
      "Regulation",
      VF_product_class
    )
  ) %>%
  mutate(
    VF_product_class = if_else(
      str_detect(VF_product_class, regex("alginate biosynthesis regulatory protein AlgR", ignore_case = TRUE)),
      "Regulation",
      VF_product_class
    )
  ) %>%
  mutate(
    VF_product_class = if_else(
      str_detect(VF_product_class, regex("paerucumarin biosynthesis", ignore_case = TRUE)),
      "Nutritional/Metabolic factor",
      VF_product_class
    )
  ) %>%
  mutate(
    VF_product_class = if_else(
      str_detect(VF_product_class, regex("acyl carrier", ignore_case = TRUE)),
      "Biofilm",
      VF_product_class
    )
  ) %>% 
  mutate(
    VF_product_class = if_else(
      str_detect(VF_product_class, regex("AlgW protein", ignore_case = TRUE)),
      "Exoenzyme",
      VF_product_class
    )
  ) %>% 
  mutate(
    VF_product_class = if_else(
      str_detect(VF_product_class, regex("MbtH-like", ignore_case = TRUE)),
      "Regulation",
      VF_product_class
    )
  ) %>% 
  mutate(
    VF_product_class = if_else(
      str_detect(VF_product_class, regex("autoinducer synthesis", ignore_case = TRUE)),
      "Regulation",
      VF_product_class
    )
  ) %>% 
  mutate(
    VF_product_class = if_else(
      str_detect(VF_product_class, regex("exporter", ignore_case = TRUE)),
      "Nutritional/Metabolic factor",
      VF_product_class
    )
  )%>%
  mutate(
    VF_product_class = if_else(
      str_detect(VF_product_class, regex("cyclic beta 1-2 glucan synthetase", ignore_case = TRUE)),
      "Biofilm",
      VF_product_class
    )
  ) %>%
  mutate(
    VF_product_class = if_else(
      str_detect(VF_product_class, regex("phosphatidate cytidylyltransferase", ignore_case = TRUE)),
      "Biofilm",
      VF_product_class
    )
  ) %>%
  mutate(
    VF_product_class = if_else(
      str_detect(VF_product_class, regex("undecaprenyl diphosphate synthase", ignore_case = TRUE)),
      "Biofilm",
      VF_product_class
    )
  ) %>%
  mutate(
    VF_product_class = if_else(
      str_detect(VF_product_class, regex("phosphoglucomutase", ignore_case = TRUE)),
      "Biofilm",
      VF_product_class
    )
  ) %>%
  mutate(
    VF_product_class = if_else(
      str_detect(VF_product_class, regex("\\bentB\\b|isochorismatase", ignore_case = TRUE)),
      "Nutritional/Metabolic factor",
      VF_product_class
    )
  ) %>%
  mutate(
    VF_product_class = if_else(
      str_detect(VF_product_class, regex("\\bentA\\b|dihydroxybenzoate dehydrogenase", ignore_case = TRUE)),
      "Nutritional/Metabolic factor",
      VF_product_class
    )
  ) %>%
  mutate(
    VF_product_class = if_else(
      str_detect(VF_product_class, regex("\\bfabZ\\b|hydroxymyristoyl ACP dehydratase", ignore_case = TRUE)),
      "Biofilm",
      VF_product_class
    )
  ) %>%
  mutate(
    VF_product_class = if_else(
      str_detect(VF_product_class, regex("\\blpxA\\b|acyltransferase", ignore_case = TRUE)),
      "Biofilm",
      VF_product_class
    )
  ) %>%
  mutate(
    VF_product_class = if_else(
      str_detect(VF_product_class, regex("\\blpxD\\b|glucosamine N-acyltransferase", ignore_case = TRUE)),
      "Biofilm",
      VF_product_class
    )
  ) %>%
  mutate(
    VF_product_class = if_else(
      str_detect(VF_product_class, regex("\\bkdsA\\b|phosphooctonate aldolase", ignore_case = TRUE)),
      "Biofilm",
      VF_product_class
    )
  ) %>%
  mutate(
    VF_product_class = if_else(
      str_detect(VF_product_class, regex("\\bkdsB\\b|octulosonate cytidylyltransferase", ignore_case = TRUE)),
      "Biofilm",
      VF_product_class
    )
  ) %>%
  mutate(
    VF_product_class = if_else(
      str_detect(VF_product_class, regex("\\bpvdH\\b|diaminobutyrate", ignore_case = TRUE)),
      "Nutritional/Metabolic factor",
      VF_product_class
    )
  ) %>%
  mutate(
    VF_product_class = if_else(
      str_detect(VF_product_class, regex("\\bwaa[PFG]\\b|heptosyltransferase|glucosyltransferase", ignore_case = TRUE)),
      "Biofilm",
      VF_product_class
    )
  ) %>%
  mutate(
    VF_product_class = if_else(
      str_detect(VF_product_class, regex("\\balg8\\b|alginate-c5-mannuronan-epimerase", ignore_case = TRUE)),
      "Biofilm",
      VF_product_class
    )
  ) %>%
  mutate(
    VF_product_class = if_else(
      str_detect(VF_product_class, regex("\\balgD\\b|GDP-mannose 6-dehydrogenase", ignore_case = TRUE)),
      "Biofilm",
      VF_product_class
    )
  ) %>%
  mutate(
    VF_product_class = if_else(
      str_detect(VF_product_class, regex("\\bbplB\\b|acetyltransferase", ignore_case = TRUE)),
      "Exoenzyme",
      VF_product_class
    )
  ) %>%
  mutate(
    VF_product_class = if_else(
      str_detect(VF_product_class, regex("\\bentC\\b|isochorismate synthase", ignore_case = TRUE)),
      "Nutritional/Metabolic factor",
      VF_product_class
    )
  ) %>%
  mutate(
    VF_product_class = if_else(
      str_detect(VF_product_class, regex("\\bentE\\b|AMP ligase", ignore_case = TRUE)),
      "Nutritional/Metabolic factor",
      VF_product_class
    )
  ) %>%
  mutate(
    VF_product_class = if_else(
      str_detect(VF_product_class, regex("\\bpmm\\b|phosphomannomutase", ignore_case = TRUE)),
      "Biofilm",
      VF_product_class
    )
  ) %>%
  mutate(
    VF_product_class = if_else(
      str_detect(VF_product_class, regex("mannose-1-phosphate guanylyltransferase", ignore_case = TRUE)),
      "Biofilm",
      VF_product_class
    )
  ) %>%
  mutate(
    VF_product_class = if_else(
      str_detect(VF_product_class, regex("mannose-6-phosphate isomerase", ignore_case = TRUE)),
      "Biofilm",
      VF_product_class
    )
  ) %>%
  mutate(
    VF_product_class = if_else(
      str_detect(VF_product_class, regex("mannosyltransferase C", ignore_case = TRUE)),
      "Biofilm",
      VF_product_class
    )
  ) %>%
  mutate(
    VF_product_class = if_else(
      str_detect(VF_product_class, regex("protein phosphatase PppB|HSI-I", ignore_case = TRUE)),
      "Effector delivery system",
      VF_product_class
    )
  ) %>%
  mutate(
    VF_product_class = if_else(
      str_detect(VF_product_class, regex("phosphomannose isomerase|GDP-mannose pyrophosphorylase", ignore_case = TRUE)),
      "Biofilm",
      VF_product_class
    )
  ) %>%
  mutate(
    VF_product_class = if_else(
      str_detect(VF_product_class, regex("alginate o-acetyltransferase AlgI", ignore_case = TRUE)),
      "Biofilm",
      VF_product_class
    )
  )

```

```{r vf-class-count}
# Count the number of VFs belonging to each class
vf_class_count <- vf_classifications_tidy %>% 
  filter(CAT_superkingdom == "Bacteria") %>% 
  select(VF_product_class) %>% 
  group_by(VF_product_class) %>%
  summarise(GeneCount = n(),
            .groups = "drop") %>% 
  mutate(VF_product_class = gsub("Effector delivery system", "Effector delivery", VF_product_class),
         VF_product_class = gsub("Immune modulation", "Immune", VF_product_class),
         VF_product_class = gsub("Nutritional/Metabolic factor", "Nutritional", VF_product_class)) %>% 
  mutate(VF_product_class = reorder(VF_product_class, -GeneCount))

# Count the number of VF product classes
vf_class <- length(unique(vf_class_count$VF_product_class))
```

```{r ARG-VF-class-plot-make, eval=FALSE}
# Set font
par(family = "Aptos")

# Plot the CAT bacteria vs mechanism pheatmap
CAT_bacteria_class_vs_mechanism_pheatmap <- pheatmap(CAT_bacteria_mech_drug_mat,
                                scale = "none",
                                cluster_rows = FALSE,
                                cluster_cols = FALSE,
                                color = drug_vs_mech_colors,
                                breaks = drug_vs_mech_breaks,
                                show_rownames = TRUE,
                                fontsize = 12,
                                main = "",
                                border_color = "grey",
                                legend = FALSE,
                                silent = TRUE,
                                display_numbers = drug_vs_mech_labels,
                                fontsize_number = 10,
                                number_color = "black")

png(
  filename = "plots/ARG_class_mech_pheatmap.png",
  width = 1200,
  height = 2200,
  res = 300
)

# Draw the silent pheatmap manually
grid::grid.draw(CAT_bacteria_class_vs_mechanism_pheatmap$gtable)

dev.off()

# Plot the VF class count column chart
CAT_bacteria_VF_class <- ggplot(vf_class_count, aes(x = VF_product_class, y = GeneCount)) +
  geom_bar(stat = "identity", colour = "black", fill = "#1B9E77", alpha = 0.4) +
  theme_minimal(base_family = "Aptos") +
  scale_y_continuous(expand = expansion(mult = c(0, 0)), limits = c(0, 100), breaks = seq(0, 100, by = 20)) +
  theme(
    panel.grid.major = element_blank(),
    panel.grid.minor = element_blank(),
    axis.title.y = element_text(size = 65, color = "black", lineheight = 0.3, margin = margin(r = 5)),
    axis.title.x = element_blank(),
    axis.text.y = element_text(size = 50, color = "black"),
    axis.text.x = element_text(size = 50, color = "black", angle = 90, hjust = 1, vjust = 0.5),
    axis.line = element_line(color = "black", linewidth = 0.5),
    axis.ticks = element_line(color = "black", linewidth = 0.5),
    plot.margin = margin(t = 50, r = 10, b = 10, l = 30), 
  ) +
    labs(x = "",
       y = "Gene count")

ggsave(
  filename = "plots/ARG_class_VF_barplot.png",
  plot = CAT_bacteria_VF_class + theme(text = element_text(family = "Aptos")),
  width = 1200 / 300,   # convert pixels to inches
  height = 2100 / 300,  # convert pixels to inches
  dpi = 300,
  units = "in",
  bg = "white"
)

# Read images
ARG_class_img <- image_read("plots/ARG_class_mech_pheatmap.png")
VF_class_img <- image_read("plots/ARG_class_VF_barplot.png")

# Annotate with labels
ARG_class_img <- image_annotate(ARG_class_img, "A", size = 100, font = "Aptos",
                       gravity = "northwest", color = "black", location = "+30+-20")
VF_class_img <- image_annotate(VF_class_img, "B", size = 100, font = "Aptos",
                       gravity = "northwest", color = "black", location = "+30+-20")

# Combine side by side
combined_ARG_VF_class_img <- image_append(c(ARG_class_img, VF_class_img))

# Save final combined PNG
image_write(combined_ARG_VF_class_img, path = "plots/combined_ARG_VF_class_plot.png", format = "png")
```

```{r ARG-VF-class-plot, fig.cap="**Antibiotic resistance and virulence genes in the BSFL bacterial midgut community.** (**A**) The incidences of resistance predicted to be conferred to each of the antibiotic classes by the ARGs of each resistance mechanism for ARGs identified on bacterial contigs. (**B**) The number of predicted VF genes on bacterial contigs belonging to each VF class.", out.width="100%"}
knitr::include_graphics("plots/combined_ARG_VF_class_plot.png")
```

In total, `r CAT_total_args` ARGs were predicted across all contigs, of which `r CAT_bacteria_arg_count` genes were identified on `r CAT_bacteria_arg_contigs` unique bacterial contigs. The remaining ARG-carrying contigs were not assigned taxonomically. The relative abundance of bacterial contigs carrying ARGs accounted for `r signif(CAT_bacteria_arg_relabun, 3)` % of the total bacterial contig abundance. In comparison, `r CAT_total_vfs` VFs were predicted across all contigs, of which `r CAT_bacteria_vf_count` were identified on `r CAT_bacteria_vf_contigs` bacterial contigs. Again, all other VF-carrying contigs were unassigned. The bacterial VF-carrying contigs accounted for `r signif(CAT_bacteria_vf_relabun, 3)` % of the total bacterial contig abundance. The entire database of all predicted ARGs and VFs alongside relative abundances across samples and contig taxonomic classifications can be found in **`r get_supp("ARG, VF, and Plasmid database")`**.

In total, `r CAT_ARG_class_sum` incidences of resistance were predicted to be conferred to `r ARG_class_num` classes of antibiotic compound by ARGs predicted on bacterial contigs, where each gene may confer an incidence of resistance to multiple antibiotic classes. (**Supplementary table \@ref(tab:antibiotic-class-count-tab)**). The fluoroquinolone class of antibiotics had the highest number of incidences of resistance conferred against it with a total of `r antibiotic_class_count$Count[antibiotic_class_count$Class == "Fluoroquinolone"]` incidences of resistance. Antibiotic efflux was the most abundant encoded mechanism of resistance with `r mechanism_df$Count[mechanism_df$Mechanism == "Efflux"]` genes which conferred `r CAT_bacteria_mech_drug_df_sum$total_count[CAT_bacteria_mech_drug_df_sum$mechanism == "Efflux"]` incidences of resistance against `r CAT_bacteria_mech_class_df_sum$total_classes[CAT_bacteria_mech_class_df_sum$mechanism == "Efflux"]` classes of antibiotics (**Figure \@ref(fig:ARG-VF-class-plot)A**). Smaller numbers were predicted to be conferred by any other mechanism. Additionally, `r mechanism_df$Count[mechanism_df$Mechanism == "Unclassified"]` ARGs were unclassified in their mechanism of action and conferred `r CAT_bacteria_mech_drug_df_sum$total_count[CAT_bacteria_mech_drug_df_sum$mechanism == "Unclassified"]` incidences of resistance against `r CAT_bacteria_mech_class_df_sum$total_classes[CAT_bacteria_mech_class_df_sum$mechanism == "Unclassified"]` classes of antibiotics. 

Regarding virulence, motility class VFs were the most common with `r vf_class_count$GeneCount[vf_class_count$VF_product_class == "Motility"]` VF genes predicted for this class, followed by `r vf_class_count$GeneCount[vf_class_count$VF_product_class == "Biofilm"]` for biofilm class VFs (**Figure \@ref(fig:ARG-VF-class-plot)B**). In total, `r num_to_word(vf_class)` unique modes of virulence of the 14 possible VF product classes in the VFDB database were encoded by genes in the bacterial community of the BSFL midgut.

```{r phylum-ARG}
# Now I want to link the phyla to the predicted ARGs with an alluvial plot
# We will use the CAT classifications as a conservative look at linking phyla to ARGs
phylum_class_arg_df <- arg_classifications %>% 
  filter(CAT_superkingdom == "Bacteria") %>% 
  select(c(contig, CAT_phylum, ARO_drug_code)) %>% 
  mutate(CAT_phylum = ifelse(is.na(CAT_phylum), "Unclassified", CAT_phylum)) %>% 
  separate_rows(ARO_drug_code, sep = ";") %>% 
  group_by(CAT_phylum, ARO_drug_code) %>%
  summarise(GeneCount = n(),
            .groups = "drop")

# Sum the counts for each phyla for inline referencing
phylum_class_arg_sum <- phylum_class_arg_df %>% 
  group_by(CAT_phylum) %>% 
  summarise(TotalClasses = n(),
            TotalCount = sum(GeneCount))

# Make a matrix for a heatmap
phylum_class_arg_mat <- phylum_class_arg_df %>% 
  left_join(antibiotic_class_count %>% select(c(Code, Class)) %>% rename(ARO_drug_code = Code), by = "ARO_drug_code") %>% 
  select(-ARO_drug_code) %>% 
  pivot_wider(names_from = CAT_phylum, values_from = GeneCount, values_fill = 0) %>%
  column_to_rownames("Class")

# Log the values for the colour with a +1 pseudocount
phylum_class_arg_mat_log <- phylum_class_arg_mat %>% 
  as.matrix() %>% 
  log1p()

# Convert the original count dataframe to a numeric matrix
phylum_class_arg_mat_labels <- phylum_class_arg_mat
```

```{r phylum-VF}
# Now I want to link the phyla to the predicted VFs based on the product class with an alluvial plot
# We will use the CAT classifications as a conservative look at linking phyla to VFs
phylum_class_vf_df <- vf_classifications_tidy %>% 
  filter(CAT_superkingdom == "Bacteria") %>% 
  select(c(contig, CAT_phylum, VF_product_class)) %>% 
  mutate(CAT_phylum = ifelse(is.na(CAT_phylum), "Unclassified", CAT_phylum)) %>%
  group_by(CAT_phylum, VF_product_class) %>%
  summarise(GeneCount = n(),
            .groups = "drop") %>% 
  mutate(VF_product_class = gsub("Effector delivery system", "Effector delivery", VF_product_class),
         VF_product_class = gsub("Immune modulation", "Immune", VF_product_class),
         VF_product_class = gsub("Nutritional/Metabolic factor", "Nutritional", VF_product_class))

# Sum the counts for each phyla for inline referencing
phylum_class_vf_sum <- phylum_class_vf_df %>% 
  group_by(CAT_phylum) %>% 
  summarise(TotalClasses = n(),
            TotalCount = sum(GeneCount))

# Make a matrix for a heatmap
phylum_class_vf_mat <- phylum_class_vf_df %>% 
  pivot_wider(names_from = CAT_phylum, values_from = GeneCount, values_fill = 0) %>% 
  column_to_rownames("VF_product_class")

# Log the values for the colour with a +1 pseudocount
phylum_class_vf_mat_log <- phylum_class_vf_mat %>% 
  as.matrix() %>% 
  log1p()

# Convert the original count dataframe to a numeric matrix
phylum_class_vf_mat_labels <- phylum_class_vf_mat
```

```{r ARG-phyla}
# I want to sum the relative abundances for plotting the abundance profile of phyla
arg_phyla_abun <- arg_classifications %>% 
  filter(CAT_superkingdom == "Bacteria") %>% 
  select(c(contig, CAT_phylum, STD_1_RelAbun, STD_2_RelAbun, STD_4_RelAbun, STD_5_RelAbun, STD_7_RelAbun, STD_8_RelAbun)) %>% 
  distinct() %>% 
  select(-contig) %>% 
  mutate(CAT_phylum = replace_na(CAT_phylum, "Unclassified")) %>% 
  pivot_longer(
    cols = -c(CAT_phylum),
    names_to = "Sample",
    values_to = "Abundance"
  ) %>% 
  group_by(Sample, CAT_phylum) %>%
  summarise(Abundance = sum(Abundance), .groups = "drop") %>% 
  mutate(Sample = gsub("_RelAbun", "", Sample)) %>% 
  group_by(Sample) %>% 
  mutate(Proportion = (Abundance / sum(Abundance)) * 100) %>% 
  ungroup()

# Let's merge the Pseudomonadota proportions for each sample to the general community proportions
Pseudo_prop <- arg_phyla_abun %>% 
  filter(CAT_phylum == "Pseudomonadota") %>% 
  select(-c(Abundance, CAT_phylum)) %>% 
  left_join(phylum_clean %>% 
              filter(Phylum == "Pseudomonadota") %>% 
              mutate(Sample = gsub("norm_", "", Sample),
                     Sample = gsub("_Avg", "", Sample)) %>% 
              select(-Phylum),
            by = "Sample") %>% 
  rename(ARG_prop = Proportion, Bacteria_prop = Abundance)

# Store the average proportions of ARG carrying Pseudo and general Psuedo community
arg_pseudo_prop <- mean(Pseudo_prop$ARG_prop)
bacteria_pseudo_prop <- mean(Pseudo_prop$Bacteria_prop)

# Perform a t.test
pseudo_ttest <- t.test(Pseudo_prop$Bacteria_prop, Pseudo_prop$ARG_prop, paired = TRUE)

# For proportions of contig numbers remove the unclassified at phylum level as the increased rate of unclassified taxa makes the dominance of Pseudomonadota appear smaller
arg_phylum_classified <- arg_classifications %>% 
  filter(!is.na(CAT_phylum)) %>% 
  select(contig, CAT_phylum) %>% 
  distinct() %>% 
  count(CAT_phylum, name = "Count") %>% 
  mutate(Percentage = (Count / sum(Count)) * 100)
```

```{r VF-phyla}
# I want to sum the relative abundances for plotting the abundance profile of phyla
vf_phyla_abun <- vf_classifications %>% 
  filter (CAT_superkingdom == "Bacteria") %>% 
  select(c(contig, CAT_phylum, STD_1_RelAbun, STD_2_RelAbun, STD_4_RelAbun, STD_5_RelAbun, STD_7_RelAbun, STD_8_RelAbun)) %>% 
  distinct() %>% 
  select(-contig) %>%
  mutate(CAT_phylum = replace_na(CAT_phylum, "Unclassified")) %>% 
  pivot_longer(
    cols = -c(CAT_phylum),
    names_to = "Sample",
    values_to = "Abundance"
  ) %>% 
  group_by(Sample, CAT_phylum) %>%
  summarise(Abundance = sum(Abundance), .groups = "drop") %>% 
  mutate(Sample = gsub("_RelAbun", "", Sample)) %>% 
  group_by(Sample) %>% 
  mutate(Proportion = (Abundance / sum(Abundance)) * 100) %>% 
  ungroup()

# Let's merge the Pseudomonadota proportions for each sample to the general community proportions
vf_pseudo_prop <- vf_phyla_abun %>% 
  filter(CAT_phylum == "Pseudomonadota") %>% 
  select(-c(Abundance, CAT_phylum)) %>% 
  left_join(phylum_clean %>% 
              filter(Phylum == "Pseudomonadota") %>% 
              mutate(Sample = gsub("norm_", "", Sample),
                     Sample = gsub("_Avg", "", Sample)) %>% 
              select(-Phylum),
            by = "Sample") %>% 
  rename(VF_prop = Proportion, Bacteria_prop = Abundance)

# Store the average proportions of VF carrying Pseudo and general Psuedo community
vf_community_pseudo_prop <- mean(vf_pseudo_prop$VF_prop)
bacteria_pseudo_prop <- mean(vf_pseudo_prop$Bacteria_prop)

# Perform a t.test
vf_pseudo_ttest <- t.test(vf_pseudo_prop$Bacteria_prop, vf_pseudo_prop$VF_prop, paired = TRUE)

# For proportions of contig numbers remove the unclassified at phylum level as the increased rate of unclassified taxa makes the dominance of Pseudomonadota appear smaller
vf_phylum_classified <- vf_classifications %>% 
  filter(!is.na(CAT_phylum)) %>% 
  select(contig, CAT_phylum) %>% 
  distinct() %>% 
  count(CAT_phylum, name = "Count") %>% 
  mutate(Percentage = (Count / sum(Count)) * 100)
```

```{r phylum-ARG-VF-plot, fig.cap="**Resistant and virulent phyla in the BSFL bacterial midgut community.** The number of incidences of resistance conferred by ARGs to the antibiotic classes and the counts of VF genes belonging to each product class, for genes identified on bacterial contigs classified to each bacterial phylum.", fig.width=6, fig.height=6, dpi=300}
# define common breaks across all matrices
phylum_class_all_vals <- c(phylum_class_arg_mat_log, phylum_class_vf_mat_log)
phylum_class_breaks <- seq(min(phylum_class_all_vals), max(phylum_class_all_vals), length.out = 101)
# define a common color palette
phylum_class_colors <- colorRampPalette(c("white", "#4575B4"))(100)

# Set font
par(family = "Aptos")
# Heatmap for phylum vs args
phylum_class_arg_pheatmap <- pheatmap(phylum_class_arg_mat_log,
                                scale = "none",
                                cluster_rows = FALSE,
                                cluster_cols = FALSE,
                                color = phylum_class_colors,
                                breaks = phylum_class_breaks,
                                show_rownames = TRUE,
                                fontsize = 30,
                                main = "ARGs",
                                border_color = "grey",
                                legend = FALSE,
                                silent = TRUE,
                                display_numbers = phylum_class_arg_mat_labels,
                                fontsize_number = 26,
                                number_color = "black")

phylum_class_vf_pheatmap <- pheatmap(phylum_class_vf_mat_log,
                                scale = "none",
                                cluster_rows = FALSE,
                                cluster_cols = FALSE,
                                color = phylum_class_colors,
                                breaks = phylum_class_breaks,
                                show_rownames = TRUE,
                                fontsize = 30,
                                main = "VFs",
                                border_color = "grey",
                                legend = FALSE,
                                silent = TRUE,
                                display_numbers = phylum_class_vf_mat_labels,
                                fontsize_number = 26,
                                number_color = "black")

grid.arrange(
    phylum_class_arg_pheatmap[[4]],
    phylum_class_vf_pheatmap[[4]],
    ncol = 2,
    widths = c(1.4, 1.1)
  )
```

Pseudomonadota phylum taxa contributed the most to antibiotic resistance (**Figure \@ref(fig:phylum-ARG-VF-plot)**), with `r phylum_class_arg_sum$TotalCount[phylum_class_arg_sum$CAT_phylum == "Pseudomonadota"]` total incidences of resistance conferred against `r phylum_class_arg_sum$TotalClasses[phylum_class_arg_sum$CAT_phylum == "Pseudomonadota"]` classes of antibiotics. However, `r phylum_class_arg_sum$TotalCount[phylum_class_arg_sum$CAT_phylum == "Unclassified"]` incidences of resistance against `r phylum_class_arg_sum$TotalClasses[phylum_class_arg_sum$CAT_phylum == "Unclassified"]` classes resulted from ARG genes on bacterial contigs unclassified to the phylum level. Of the total number of contigs in the bacterial community that were classified to phylum, `r signif(bacteria_phylum_classified$Percentage[bacteria_phylum_classified$Phylum == "Pseudomonadota"], 3)` % belonged to the Pseudomonadota phylum. This increased to `r signif(arg_phylum_classified$Percentage[arg_phylum_classified$CAT_phylum == "Pseudomonadota"], 3)` % when examining the bacterial ARG-carrying contigs classified to phylum.

Pseudomonadota phylum taxa contributed to all `r num_to_word(phylum_class_vf_sum$TotalClasses[phylum_class_vf_sum$CAT_phylum == "Pseudomonadota"])` VF classes identified here via `r phylum_class_vf_sum$TotalCount[phylum_class_vf_sum$CAT_phylum == "Pseudomonadota"]` predicted VFs on contigs assigned to this phylum (**Figure \@ref(fig:phylum-ARG-VF-plot)**). Only `r num_to_word(phylum_class_vf_sum$TotalCount[phylum_class_vf_sum$CAT_phylum == "Bacillota"])` VF was identified on a Bacillota assigned contig of the membrane class of VFs. The remaining `r phylum_class_vf_sum$TotalCount[phylum_class_vf_sum$CAT_phylum == "Unclassified"]` predicted VFs, spanning `r num_to_word(phylum_class_vf_sum$TotalClasses[phylum_class_vf_sum$CAT_phylum == "Unclassified"])` VF product classes, were identified on bacterial contigs unassigned to phylum. The `r signif(bacteria_phylum_classified$Percentage[bacteria_phylum_classified$Phylum == "Pseudomonadota"], 3)` % of the number of bacterial phylum-classified contigs assigned to the Pseudomonadota phylum increased to `r signif(vf_phylum_classified$Percentage[vf_phylum_classified$CAT_phylum == "Pseudomonadota"], 3)` % when investigating the bacterial VF-carrying contigs classified to phylum.

\newpage

## Plasmids in the BSFL midgut microbiota

```{r CAT-plasmid-sum}
# Filter to contigs predicted to be part of a plasmid
CAT_plasmid_contigs <- contig_db_norm %>% 
  filter(!is.na(PF_start))

# Count the number of rows which is equivalent to the number of contigs predicted to be part of plasmids
CAT_total_plasmid_contigs <- nrow(CAT_plasmid_contigs)

# Count the number of unique plasmids based on accession numbers
CAT_unique_plasmids <- length(unique(CAT_plasmid_contigs$PF_accession))

# Lastly sum the average across the relative abundance columns to calculate the total relative abundance of VF-carrying contigs
total_plasmid_relabun <- contig_db_norm %>% 
  filter(!is.na(PF_start)) %>% 
  rowwise() %>% 
  mutate(mean_rel_abun = mean(c_across(starts_with("norm_")), na.rm = TRUE)) %>% 
  ungroup() %>% 
  { sum(.$mean_rel_abun) }
```

```{r plasmid-classification-make, eval=FALSE}
# I want to see for each plasmid if it's classifications from CAT and Kraken2, and then I want to merge on the read depths across samples for a supplementary .csv file
plasmid_classifications <- CAT_plasmid_contigs %>% 
  select(c("contig", "length", "superkingdom", "kingdom", "phylum", "class", "order", "family", "genus", "species", "PF_start", "PF_end", "PF_strand", "PF_gene", "PF_coverage", "PF_coverage_map", "PF_gaps", "PF_coverage_percentage", "PF_pident", "PF_accession", "PF_product")) %>% 
  rename(CAT_superkingdom = superkingdom, CAT_kingdom = kingdom, CAT_phylum = phylum, CAT_class = class, CAT_order = order, CAT_family = family, CAT_genus = genus, CAT_species = species) %>% 
  left_join(kraken_plasmid_contigs %>% 
              select(c("contig", "lineage_1", "lineage_2", "lineage_3", "lineage_4", "lineage_5", "lineage_6", "lineage_7", "lineage_8")) %>% 
              rename(kraken2_superkingdom = lineage_1, kraken2_kingdom = lineage_2, kraken2_phylum = lineage_3, kraken2_class = lineage_4, kraken2_order = lineage_5, kraken2_family = lineage_6, kraken2_genus = lineage_7, kraken2_species = lineage_8), by = c("contig")) %>% 
  mutate(kraken2_superkingdom = case_when(
    kraken2_kingdom %in% c("Pseudomonadati", "Bacillati", "unclassified Bacteria kingdom") ~ "Bacteria",
    TRUE ~ "Unclassified"
  ),
  CAT_superkingdom = ifelse(is.na(CAT_superkingdom), "Unclassified", CAT_superkingdom)) %>%
  mutate(across(kraken2_kingdom:kraken2_species, ~ ifelse(grepl("^unclassified", .), NA, .))) %>% 
  left_join(contig_db_norm %>% select(c("contig", "norm_STD_1_Avg", "norm_STD_2_Avg", "norm_STD_4_Avg", "norm_STD_5_Avg", "norm_STD_7_Avg", "norm_STD_8_Avg")), by = "contig") %>% 
  rename(STD_1_RelAbun = norm_STD_1_Avg, STD_2_RelAbun = norm_STD_2_Avg, STD_4_RelAbun = norm_STD_4_Avg, STD_5_RelAbun = norm_STD_5_Avg, STD_7_RelAbun = norm_STD_7_Avg, STD_8_RelAbun = norm_STD_8_Avg) %>% 
  select(c("contig", "length", "PF_start", "PF_end", "PF_strand", "PF_gene", "PF_coverage", "PF_coverage_map", "PF_gaps", "PF_coverage_percentage", "PF_pident", "PF_accession", "PF_product", "CAT_superkingdom", "CAT_kingdom", "CAT_phylum", "CAT_class", "CAT_order", "CAT_family", "CAT_genus", "CAT_species", "kraken2_superkingdom", "kraken2_kingdom", "kraken2_phylum", "kraken2_class", "kraken2_order", "kraken2_family", "kraken2_genus", "kraken2_species", "STD_1_RelAbun", "STD_2_RelAbun", "STD_4_RelAbun", "STD_5_RelAbun", "STD_7_RelAbun", "STD_8_RelAbun"))
```

```{r plasmid_db_sup, eval=FALSE}
# Make a .csv file with a leading line explaining each column for a supplementary file
write.csv(plasmid_classifications, "Bioinformatics/abricate/Plasmid_db.csv", row.names = FALSE)
```

```{r plasmid-classification-read}
plasmid_classifications <- read.csv("Bioinformatics/abricate/Plasmid_db.csv", header = TRUE)
```

```{r plasmid-ARG}
# I want to find if there is any co-occurence between plasmids and ARGs to see if these ARGs are potentially mobile
plasmid_ARGs <- plasmid_classifications %>% 
  select(-c(STD_1_RelAbun, STD_2_RelAbun, STD_4_RelAbun, STD_5_RelAbun, STD_7_RelAbun, STD_8_RelAbun)) %>% 
  left_join(arg_classifications %>% select(-c(STD_1_RelAbun, STD_2_RelAbun, STD_4_RelAbun, STD_5_RelAbun, STD_7_RelAbun, STD_8_RelAbun, CAT_superkingdom, CAT_kingdom, CAT_phylum, CAT_class, CAT_order, CAT_family, CAT_genus, CAT_species, kraken2_superkingdom, kraken2_kingdom, kraken2_phylum, kraken2_class, kraken2_order, kraken2_family, kraken2_genus, kraken2_species)), by = "contig") %>% 
  filter(!is.na(ARG_gene))

# Count the number of contigs with hits against both plasmids and ARGs
co_plasmid_args <- nrow(plasmid_ARGs)

# Make a table for output
co_plasmid_args_df <- plasmid_ARGs %>% 
  select(contig, PF_accession, ARG_gene, ARG_product) %>% 
  rename(Contig = contig,"Plasmid accession" = PF_accession, "ARG name" = ARG_gene, "CARD product description" = ARG_product)
```

```{r plasmid-VF}
# I want to find if there is any co-occurence between plasmids and VFs to see if these VFs are potentially mobile
plasmid_VFs <- plasmid_classifications %>% 
  select(-c(STD_1_RelAbun, STD_2_RelAbun, STD_4_RelAbun, STD_5_RelAbun, STD_7_RelAbun, STD_8_RelAbun)) %>% 
  left_join(vf_classifications %>% select(-c(STD_1_RelAbun, STD_2_RelAbun, STD_4_RelAbun, STD_5_RelAbun, STD_7_RelAbun, STD_8_RelAbun, CAT_superkingdom, CAT_kingdom, CAT_phylum, CAT_class, CAT_order, CAT_family, CAT_genus, CAT_species, kraken2_superkingdom, kraken2_kingdom, kraken2_phylum, kraken2_class, kraken2_order, kraken2_family, kraken2_genus, kraken2_species)), by = "contig") %>% 
  filter(!is.na(VF_gene))

# Count the number of contigs with hits against both plasmids and VFs
co_plasmid_vfs <- nrow(plasmid_VFs)
```

```{r plasmid-metadata}
# I took the 24 unique plasmid accessions and searched for their entries in the PlasmidScope database to get metadata on them
# Firstly general metadata
plasmid_meta <- read.delim("Bioinformatics/abricate/plasmid_metadata.tsv", header = TRUE, sep = "\t")
```

```{r plasmid-ARG-metadata}
# Then read in the Plasmid Scope ARG metadata
plasmid_arg_meta <- read.delim("Bioinformatics/abricate/plasmid_arg_data.tsv", header = TRUE, sep = "\t")

# Now I want to investigate the resistance predicted by PlasmidScope
# Remove tricky streptogramin antibiotic IDs which cause trouble, filter out the loose predictions of ARGs as I don't want to overinflate the predictions of mobile resistance
plasmid_arg_meta_clean <- plasmid_arg_meta %>% 
  mutate(Drug.Class = gsub("; streptogramin antibiotic; streptogramin A antibiotic; streptogramin B antibiotic", "", Drug.Class)) %>% 
  filter(Cut_Off != "Loose")

# Get the number of unique accessions where the Drug.Class column is not empty e.g. resistance encoded
plasmid_arg_meta_unique <- plasmid_arg_meta_clean %>% 
  filter(Drug.Class != "") %>% 
  distinct(Accession)

# Count the number of these unique PlasmidScope resistant plasmids
plasmid_arg_meta_unique_plasmids <- nrow(plasmid_arg_meta_unique)

# Count the number of unique accessions predicted to encode resistance by PlasmidScope metadata and also by coccurrence of a predicted ARG on the contig identified as the plasmid fragment
double_resistant_plasmids <- length(intersect(plasmid_ARGs$PF_accession, unique(plasmid_arg_meta_unique$Accession)))

# Sum the total number of resistance encoding plasmids
total_resistant_plasmids <- co_plasmid_args + plasmid_arg_meta_unique_plasmids - double_resistant_plasmids       
```

Abricate prediction of plasmids identified `r CAT_total_plasmid_contigs` contigs that encoded sequences with similarity to plasmid genes in the PlasmidFinder database, with `r CAT_unique_plasmids` unique plasmid accessions identified overall. Only `r num_to_word(co_plasmid_args)` contigs were predicted to encode both a plasmid and ARG sequences, and these ARG sequences were described as plasmid-mediated (**Supplementary table \@ref(tab:plasmid-ARG-tab)**). No contigs were predicted to encode both a plasmid gene and a VF. Searching the unique plasmid accession numbers in the PlasmidScope database of ARG metadata identified `r num_to_word(plasmid_arg_meta_unique_plasmids)` of these unique plasmids recorded as encoding resistance, with `r num_to_word(double_resistant_plasmids)` one of them also identified from one of the three contigs encoding both a plasmid and an ARG sequence. Therefore, `r num_to_word(total_resistant_plasmids)` unique plasmids were identified in total either from contigs also encoding an ARG sequence, or with accessions of ARG-carrying plasmids according to the PlasmidScope database.

```{r plasmid-mobility-class}
# Now I want to look at the mobility of these 6 plasmids and the antibiotic classes they encode resistance against

# Make a slightly modified function to convert drug codes to shorter codes, as the PlasmidScope database has some differences compared to CARD:
convert_to_codes_fuzzy2 <- function(class_strings, max_dist = 2) {
  if (!requireNamespace("stringdist", quietly = TRUE)) {
    stop("Please install the 'stringdist' package: install.packages('stringdist')")
  }
  keys <- names(drug_code_map)
  keys_lower <- tolower(keys)

  vapply(class_strings, FUN.VALUE = character(1), function(class_string) {
    if (is.na(class_string) || trimws(class_string) == "") return(NA_character_)

    parts <- unlist(strsplit(class_string, ";", fixed = TRUE))
    parts <- trimws(parts)
    codes_found <- character(0)

    for (part in parts) {
      if (part == "") next
      x <- tolower(part)

      # Normalize: remove the word "antibiotic", punctuation, collapse spaces
      x <- gsub("\\bantibiotic\\b", "", x)
      x <- gsub("[[:punct:]]", " ", x)
      x <- gsub("\\s+", " ", x)
      x <- trimws(x)
      if (x == "") next

      # 1) direct substring match of any known key (fast & robust)
      exact_idx <- which(vapply(keys_lower,
                                FUN = function(k) grepl(k, x, fixed = TRUE),
                                logical(1)))
      if (length(exact_idx) > 0) {
        codes_found <- c(codes_found, drug_code_map[[ keys[exact_idx[1]] ]])
        next
      }

      # 2) match by token (split the cleaned phrase into words and try each)
      tokens <- unlist(strsplit(x, " ", fixed = TRUE))
      tokens <- tokens[nzchar(tokens)]
      token_matched <- FALSE
      for (t in tokens) {
        idx <- which(vapply(keys_lower,
                            FUN = function(k) grepl(t, k, fixed = TRUE),
                            logical(1)))
        if (length(idx) > 0) {
          codes_found <- c(codes_found, drug_code_map[[ keys[idx[1]] ]])
          token_matched <- TRUE
          break
        }
      }
      if (token_matched) next

      # 3) fuzzy fallback (compare cleaned phrase to keys)
      dists <- stringdist::stringdist(x, keys_lower, method = "osa")
      if (min(dists) <= max_dist) {
        idx <- which.min(dists)
        codes_found <- c(codes_found, drug_code_map[[ keys[idx] ]])
      } else {
        codes_found <- c(codes_found, "UNK")
      }
    } # end for parts

    if (length(codes_found) == 0) return(NA_character_)
    paste(unique(codes_found), collapse = ";")
  })
}
# Convert antibiotic classes to the shorter codes
plasmid_arg_meta_clean$Codes <- convert_to_codes_fuzzy2(plasmid_arg_meta_clean$Drug.Class)

# Take the 3 predicted ARGs from plasmids predicted before
plasmid_contig_ARGs <- plasmid_ARGs %>% 
  select(PF_accession, ARG_resistance)
# Convert antibiotic classes to the shorter codes
plasmid_contig_ARGs$drug_code <- sapply(plasmid_contig_ARGs$ARG_resistance, convert_to_codes_fuzzy)
# Tidy names and columns  
plasmid_contig_ARGs_clean <- plasmid_contig_ARGs %>% 
  select(-ARG_resistance) %>% 
  rename(Accession = PF_accession, Codes = drug_code)
  
# Row bind these resistance codes to those predicted by PlasmidScope, then collapse the codes into a single string for each accession
plasmid_arg_class <- plasmid_arg_meta_clean %>% 
  select(c(Accession, Codes)) %>% 
  group_by(Accession) %>%
  rbind(., plasmid_contig_ARGs_clean) %>% 
  summarise(
    Codes = Codes %>%
      na.omit() %>%                                   # drop NAs
      str_split(";") %>%                              # split into lists of codes
      unlist() %>%                                    # flatten to single vector
      str_trim() %>%                                  # trim spaces
      discard(~ .x == "") %>%                         # drop empty tokens
      paste(collapse = ";"),                          # rejoin with ;
    .groups = "drop"
  ) %>% 
  mutate(Codes = gsub("UNK;", "", Codes))

# Now take the plasmid accessions and names and mobility and merge to the Drug codes. Then separate by class and count the number of times resistance is encoded against each class by each plasmid
plasmid_mobility <- plasmid_meta %>%
  select(Accession, Predicted_Mobility) %>% 
  rename(Mobility = Predicted_Mobility) %>% 
  left_join(plasmid_arg_class, by = "Accession") %>% 
  filter(!is.na(Codes)) %>% 
  filter(Codes != "") %>% 
  separate_rows(Codes, sep = ";") %>% 
  left_join(antibiotic_class_count %>% select(c(Code, Class)) %>% rename(Codes = Code), by = "Codes") %>% 
  select(-Codes) %>% 
  group_by(Accession, Mobility, Class) %>% 
  summarise(Count = n()) %>% 
  mutate(Mobility = gsub("non-mobilizable", "Non-mobilisable", Mobility),
                         Mobility = gsub("mobilizable", "Mobilisable", Mobility),
         Mobility = gsub("conjugative", "Conjugative", Mobility)
  )

# Count the number of plasmids which are mobile, non-mobile, conjugative along with the counts of encoded resistance they confer
arg_plasmids_nonmobile <- plasmid_mobility %>% 
  filter(Mobility == "Non-mobilisable") %>% 
  distinct(Accession) %>% 
  nrow()
arg_plasmids_nonmobile_count <- plasmid_mobility %>% 
  filter(Mobility == "Non-mobilisable") %>% 
  pull(Count) %>% 
  sum()

arg_plasmids_mobile <- plasmid_mobility %>% 
  filter(Mobility == "Mobilisable") %>% 
  distinct(Accession) %>% 
  nrow()
arg_plasmids_mobile_count <- plasmid_mobility %>% 
  filter(Mobility == "Mobilisable") %>% 
  pull(Count) %>% 
  sum()

arg_plasmids_conjugative <- plasmid_mobility %>% 
  filter(Mobility == "Conjugative") %>% 
  distinct(Accession) %>% 
  nrow()
arg_plasmids_conjugative_count <- plasmid_mobility %>% 
  filter(Mobility == "Conjugative") %>% 
  pull(Count) %>% 
  sum()

arg_total_mobile <- arg_plasmids_mobile + arg_plasmids_conjugative
arg_total_mobile_count <- arg_plasmids_mobile_count + arg_plasmids_conjugative_count

# Also sum the total count of resistance encoded against classes
plasmid_class_resistance_count <- sum(plasmid_mobility$Count)
# And the number of unique classes
plasmid_class_resistance_unique <- length(unique(plasmid_mobility$Class))
```

```{r plasmid-ARGs-plot, fig.cap="**Plasmid-encoded resistance in the BSFL midgut community.** The predicted mobility and the number of incidences of resistance conferred against antibiotic classes encoded by ARGs for unique plasmids identified in the BSFL midgut community.", fig.height=4.5, fig.width=6, dpi=300}
# Static Sankey plot
plasmid_ARG_plot <- ggplot(plasmid_mobility,
                            aes(axis1 = Accession, axis2 = Mobility, axis3 = Class, y = Count)) + 
  geom_alluvium(aes(fill = Accession), width = 1/3) +
  geom_stratum(width = 1/3, fill = "lightgrey", color = "black", size = 0.1) +
  geom_text(stat = "stratum", aes(label = after_stat(stratum)), size = 6.9, fontface = "bold") +
  scale_x_discrete(limits = c("Accession", "Mobility", "Class"), expand = c(0.05, 0.05)) +
  scale_y_continuous(expand = c(0, 0)) +
  theme_minimal() +
  theme(axis.title = element_blank(),
        axis.text = element_blank(),
        axis.ticks = element_blank(),
        panel.grid = element_blank(),
        legend.position = "none") +
  scale_fill_viridis_d()

plasmid_ARG_plot
```

```{r plasmid-VF-metadata}
# Then read in the Plasmid Scope VF metadata
plasmid_vf_meta <- read.delim("Bioinformatics/abricate/plasmid_vf_data.tsv", header = TRUE, sep = "\t")

# Get the number of unique accessions where the VFSeqID column is not empty e.g. virulence encoded
plasmid_vf_meta_unique <- plasmid_vf_meta %>% 
  filter(VFSeq_ID != "") %>% 
  distinct(Accession)

# Count the number of these unique PlasmidScope virulent plasmids
plasmid_vf_meta_unique_plasmids <- nrow(plasmid_vf_meta_unique)

# Merge the plasmid mobility
plasmid_vf_mobility <- plasmid_meta %>%
  select(Accession, Predicted_Mobility) %>% 
  rename(Mobility = Predicted_Mobility) %>% 
  left_join(plasmid_vf_meta %>% 
              filter(VFSeq_ID != "") %>% 
              select(Accession, VFID, VF_Name, Vfcategory, Product), by = "Accession") %>% 
  filter(!is.na(VFID)) %>% 
  mutate(Mobility = gsub("conjugative", "Conjugative", Mobility)) %>% 
  rename(ID = VFID, Name = VF_Name, Category = Vfcategory)

# Count the rows that are mobile
plasmid_vf_mobile <- plasmid_vf_mobility %>% 
  filter(Mobility == "Conjugative") %>% 
  nrow()
```

```{r plasmid-arg-vf}
# Get the unique accessions of ARG-carrying plasmids
unique_plasmid_arg <- unique(plasmid_mobility$Accession)
# Get the unique accessions of VF-carrying plasmids
unique_plasmid_vf <- unique(plasmid_vf_mobility$Accession)

# Find the intersect
plasmid_arg_vf_intersect <- intersect(unique_plasmid_arg, unique_plasmid_vf)
# Get the length of the intersect
plasmid_arg_vf_intersect_length <- length(plasmid_arg_vf_intersect)
```

Based on PlasmidScope metadata of the unique plasmid accessions identified in the contig coassembly by PlasmidFinder, `r num_to_word(arg_plasmids_nonmobile)` plasmids were predicted to be non-mobilisable, and `r num_to_word(arg_total_mobile)` to be mobilisable, with `r num_to_word(arg_plasmids_conjugative)` of them through means of conjugation. In total, `r plasmid_class_resistance_count` incidences of resistance were conferred against `r plasmid_class_resistance_unique` unique classes of antibiotic compounds by the ARG genes predicted to occur on the plasmids identified in the BSFL midgut community (**Figure \@ref(fig:plasmid-ARGs-plot)**). Furthermore, `r arg_total_mobile_count` of the incidences of resistance were encoded by the `r num_to_word(arg_total_mobile)` mobilisable plasmids. 

Searching the accessions of the plasmids against the PlasmidScope VF metadata identified `r num_to_word(plasmid_vf_meta_unique_plasmids)` unique plasmids predicted to encode VF sequences. Both plasmids (`r plasmid_arg_vf_intersect[1]` and `r plasmid_arg_vf_intersect[2]`) were predicted to be mobile (**Supplementary table \@ref(tab:plasmid-VF-tab)**) and identified as containing ARG sequences (**Figure \@ref(fig:plasmid-ARGs-plot)**). The entire database of predicted plasmids alongside relative abundances across samples and contig taxonomic classifications can be seen in **`r get_supp("ARG, VF, and Plasmid database")`**.

\newpage

## Reconstruction of bacterial MAGs in the BSFL gut community

```{r MAG-import}
# Import the taxonomic classifications of medium-quality-bins
MAG_tax_medium <- read.delim("Bioinformatics/MAG_classification/medium_quality_bins/gtdbtk.bac120.summary.tsv", header = T, sep = "\t")

# Extract unique contigs from arg_classification and vf_classification database
arg_contigs <- unique(arg_classifications$contig)
vf_contigs <- unique(vf_classifications$contig)
```

```{r MAG-posthoc-filter}
# I want to remove the bins (MAGs) where a FastANI reference is not achieved, and the msa_percent value from tree placement is less than 50 %
MAG_tax_filtered <- MAG_tax_medium %>% 
  filter(!(fastani_reference == "N/A" & msa_percent < 50))
```

```{r MAG-tax}
# Extract the taxonomies and separate into separate columns, then update the old names of Firmicutes and Pseudomonadota, and remove the provisional split names with _A etc. suffixes. Also make the unofficaily named genera clearer by including the family name
MAG_tax <- MAG_tax_filtered %>% 
  select(c(user_genome, classification)) %>% 
  rename(MAG = user_genome) %>% 
  separate(
    col = classification,
    into = c("Kingdom", "Phylum", "Class", "Order", "Family", "Genus", "Species"),
    sep = ";",
    fill = "right"
  ) %>% 
  mutate(across(c(Kingdom, Phylum, Class, Order, Family, Genus, Species), ~ substr(., 4, nchar(.)))) %>% 
  mutate(Phylum = gsub("Firmicutes", "Bacillota", Phylum),
         Phylum = gsub("Proteobacteria", "Pseudomonadota", Phylum)) %>% 
  mutate(across(where(is.character),
                ~ gsub("_[A-Z]+", "", .))) %>% 
  mutate(Genus = gsub("UBA4096", "Bacteriovoracaceae_UBA4096", Genus),
         Genus = gsub("63-22", "Rhizobiaceae_63-22", Genus))

# Count the number of MAGs in total which is also the number identified to genus
MAG_num <- nrow(MAG_tax)
# Count the numer of MAGs with species level of assignment
MAG_species_num <- nrow(MAG_tax %>% filter(Species != ""))
```

```{r MAG-tax-db, eval=FALSE}
# Write the MAG taxonomic assignment results to a .csv file for supplementary
write.csv(MAG_tax_medium, "Bioinformatics/MAG_classification/medium_quality_bins/MAG_taxonomies.csv", row.names=FALSE)
```


```{r MAG-phylum}
# Count the number of MAGs belonging to each phylum and then calculate proportions and data for a donut chart
MAG_phylum <- MAG_tax %>% 
  group_by(Phylum) %>% 
  summarise(Count = n(), .groups = "drop") %>% 
  mutate(Fraction = Count / sum(Count),
         ymax = cumsum(Fraction),
         ymin = c(0, head(ymax, n = -1)),
         label_position = (ymax + ymin) / 2,
         label = paste0(Count))
```

```{r MAG-ARGs}
# I already have list of the unique args which I can use to track ARGs into the bins
# List all .fasta files in the folder containing the medium-quality bins
medium_bin_fastas <- list.files("Bioinformatics/medium_quality_bins", pattern = "\\.(fa|fasta)$", full.names = TRUE)

# Initialize result list (create a list to be filled by the loop)
MAG_args <- data.frame(contig = character(), MAG = character(), stringsAsFactors = FALSE)

# Loop through each fasta file and search headers for the contig IDS I am interested in
for (file in medium_bin_fastas) {
  # Read fasta lines
  fasta_lines <- readLines(file)
  
  # Extract only headers (lines starting with ">")
  headers <- grep("^>", fasta_lines, value = TRUE)
  
  # Clean headers to extract contig IDs (remove ">")
  contigs_in_bin <- sub("^>(\\S+).*", "\\1", headers)
  
  # Find matching contigs
  matched_contigs <- contigs_in_bin[contigs_in_bin %in% arg_contigs]
  
  # If matches found, record them
  if (length(matched_contigs) > 0) {
    temp <- data.frame(
      contig = matched_contigs,
      MAG = sub("\\.fa$", "", basename(file)),
      stringsAsFactors = FALSE
    )
    MAG_args <- rbind(MAG_args, temp)
  }
}

# Now I want to merge the arg information for contigs to the bins, I will actually merge the contig to bin info to the args, because the arg dataframe has multiple rows per contig so a left_join works. Then I will add the taxonomic classification of bins based on the bin name
MAG_args_df <- arg_classifications %>% 
  select(contig, ARG_gene, ARG_resistance, ARO_drug_class, ARO_resistance_mechanism, ARO_drug_code) %>% 
  left_join(MAG_args, by = "contig") %>% 
  filter(!is.na(MAG)) %>% 
  left_join(MAG_tax, by = "MAG")

# Get the unique number of MAGs containing contigs with ARGs predicted
resistant_MAGs <- length(unique(MAG_args_df$MAG))
# Get the number of unique contigs carrying resistance that made it into the MAGs
resistant_MAG_contigs <- length(unique(MAG_args_df$contig))
```

```{r MAG-genus-ARG}
# Now with the MAGs I can explore the taxa contributions to antibiotic classes with more specificity
# But I also want to link the genus to phyla so that I can compare with contigs, which I explored mainly at the phylum level
MAG_genus_arg <- MAG_args_df %>% 
  select(Phylum, Genus, ARO_drug_code) %>% 
  separate_rows(ARO_drug_code, sep = ";") %>% 
  group_by(Phylum, Genus, ARO_drug_code) %>%
  summarise(GeneCount = n(),
            .groups = "drop")

# Sum the counts for each phyla for inline referencing
MAG_phylum_sum <- MAG_genus_arg %>% 
  group_by(Phylum) %>% 
  summarise(TotalClasses = length(unique(ARO_drug_code)),
            TotalCount = sum(GeneCount))

# Get the total counts overall
MAG_arg_class_counts <- sum(MAG_phylum_sum$TotalCount)
# Get the number of unqiue classes
MAG_arg_classes <- length(unique(MAG_genus_arg$ARO_drug_code))

# Sum the counts for each genus for inline referencing
MAG_genus_sum <- MAG_genus_arg %>% 
  group_by(Genus) %>% 
  summarise(TotalClasses = n(),
            TotalCount = sum(GeneCount))

# Make a matrix for a heatmap
MAG_genus_arg_mat <- MAG_genus_arg %>% 
  select(-Phylum) %>% 
  pivot_wider(names_from = Genus, values_from = GeneCount, values_fill = 0) %>% 
  left_join(antibiotic_class_count %>% select(c(Code, Class)) %>% rename(ARO_drug_code = Code), by = "ARO_drug_code") %>% 
  select(-ARO_drug_code) %>% 
  column_to_rownames("Class")

# Log the values for the colour with a +1 pseudocount
MAG_genus_arg_mat_log <- MAG_genus_arg_mat %>% 
  as.matrix() %>% 
  log1p()

# Convert the original count dataframe to a numeric matrix
MAG_genus_arg_mat_labels <- MAG_genus_arg_mat

# Annotation for ARGs
MAG_genus_arg_ann <- MAG_genus_arg %>%
  distinct(Genus, Phylum) %>%
  tibble::column_to_rownames("Genus")
```

```{r MAG-VFs}
# I already have list of the unique vfs which I can use to track VFs into the bins
# Initialize result list (create a list to be filled by the loop)
MAG_vfs <- data.frame(contig = character(), MAG = character(), stringsAsFactors = FALSE)

# Loop through each fasta file and search headers for the contig IDS I am interested in
for (file in medium_bin_fastas) {
  # Read fasta lines
  fasta_lines <- readLines(file)
  
  # Extract only headers (lines starting with ">")
  headers <- grep("^>", fasta_lines, value = TRUE)
  
  # Clean headers to extract contig IDs (remove ">")
  contigs_in_bin <- sub("^>(\\S+).*", "\\1", headers)
  
  # Find matching contigs
  matched_contigs <- contigs_in_bin[contigs_in_bin %in% vf_contigs]
  
  # If matches found, record them
  if (length(matched_contigs) > 0) {
    temp <- data.frame(
      contig = matched_contigs,
      MAG = sub("\\.fa$", "", basename(file)),
      stringsAsFactors = FALSE
    )
    MAG_vfs <- rbind(MAG_vfs, temp)
  }
}

# Now I want to merge the arg information for contigs to the bins, I will actually merge the contig to bin info to the args, because the arg dataframe has multiple rows per contig so a left_join works. Then I will add the taxonomic classification of bins based on the bin name
MAG_vfs_df <- vf_classifications_tidy %>% 
  select(contig, VF_gene, VF_product_class) %>% 
  left_join(MAG_vfs, by = "contig") %>% 
  filter(!is.na(MAG)) %>% 
  left_join(MAG_tax, by = "MAG")

# Get the unique number of MAGs containing contigs with VFs predicted
virulent_MAGs <- length(unique(MAG_vfs_df$MAG))
# Get the number of unique contigs carrying virulence that made it into the MAGs
virulent_MAG_contigs <- length(unique(MAG_vfs_df$contig))
```

```{r MAG-genus-VF}
# Now with the MAGs I can explore the taxa contributions to virulence classes with more specificity
# But I also want to link the genus to phyla so that I can compare with contigs, which I explored mainly at the phylum level
MAG_genus_vf <- MAG_vfs_df %>% 
  select(Phylum, Genus, VF_product_class) %>% 
  group_by(Phylum, Genus, VF_product_class) %>%
  mutate(VF_product_class = gsub("Effector delivery system", "Effector delivery", VF_product_class),
         VF_product_class = gsub("Immune modulation", "Immune", VF_product_class),
         VF_product_class = gsub("Nutritional/Metabolic factor", "Nutritional", VF_product_class)) %>% 
  summarise(GeneCount = n(),
            .groups = "drop")

# Sum the counts for each phyla for inline referencing
MAG_vf_phylum_sum <- MAG_genus_vf %>% 
  group_by(Phylum) %>% 
  summarise(TotalClasses = length(unique(VF_product_class)),
            TotalCount = sum(GeneCount))

# Get the total counts overall
MAG_vf_class_counts <- sum(MAG_vf_phylum_sum$TotalCount)
# Get the number of unqiue classes
MAG_vf_classes <- length(unique(MAG_genus_vf$VF_product_class))

# Sum the counts for each genus for inline referencing
MAG_vf_genus_sum <- MAG_genus_vf %>% 
  group_by(Genus) %>% 
  summarise(TotalClasses = n(),
            TotalCount = sum(GeneCount))

# Make a matrix for a heatmap
MAG_genus_vf_mat <- MAG_genus_vf %>% 
  select(-Phylum) %>% 
  pivot_wider(names_from = Genus, values_from = GeneCount, values_fill = 0) %>%
  column_to_rownames("VF_product_class") 

# Log the values for the colour with a +1 pseudocount
MAG_genus_vf_mat_log <- MAG_genus_vf_mat %>% 
  as.matrix() %>% 
  log1p()

# Convert the original count dataframe to a numeric matrix
MAG_genus_vf_mat_labels <- MAG_genus_vf_mat

# Annotation for VFs
MAG_genus_vf_ann <- MAG_genus_vf %>%
  distinct(Genus, Phylum) %>%
  tibble::column_to_rownames("Genus")
```

```{r MAG-genus-ARG-VF-plot, fig.cap="**MAG contributions to antibiotic resistance and virulence.** The number of incidences of resistance conferred to the antibiotic classes for ARGs, and the number of VF genes assigned to each VF product class for genes identified on contigs binned into MAGs. The colours at the top of each column indicate the phylum to which the genus shown at the bottom of the column belongs.", fig.width=7, fig.height=6, dpi=300}
# define common breaks across all matrices
MAG_genus_all_vals <- c(MAG_genus_arg_mat_log, MAG_genus_vf_mat_log)
MAG_genus_breaks <- seq(min(MAG_genus_all_vals), max(MAG_genus_all_vals), length.out = 101)
# define a common color palette
MAG_genus_colors <- colorRampPalette(c("white", "#4575B4"))(100)

# Combine all unique phyla across both datasets
all_heatmap_phyla <- unique(c(MAG_genus_arg_ann$Phylum, MAG_genus_vf_ann$Phylum))
# Apply to your Phylum color mapping
heatmap_phylum_colors <- setNames(
  add_opacity(my_wes_palette(length(all_heatmap_phyla)), alpha = 0.7),
  all_heatmap_phyla
)
# Wrap it for pheatmap’s annotation_colors argument
heatmap_annotation_colors <- list(Phylum = heatmap_phylum_colors)

# Set font
par(family = "Aptos")
# Heatmap for MAG genus vs args
MAG_genus_arg_pheatmap <- pheatmap(MAG_genus_arg_mat_log,
                                scale = "none",
                                cluster_rows = FALSE,
                                cluster_cols = FALSE,
                                color = MAG_genus_colors,
                                breaks = MAG_genus_breaks,
                                show_rownames = TRUE,
                                fontsize = 24,
                                main = "ARGs",
                                border_color = "grey",
                                legend = FALSE,
                                silent = TRUE,
                                display_numbers = MAG_genus_arg_mat_labels,
                                fontsize_number = 22,
                                number_color = "black",
                                annotation_col = MAG_genus_arg_ann,
                                annotation_colors = heatmap_annotation_colors,
                                annotation_legend = FALSE,
                                annotation_names_col = FALSE)

# Heatmap for MAG genus vs vfs
MAG_genus_vf_pheatmap <- pheatmap(MAG_genus_vf_mat_log,
                                scale = "none",
                                cluster_rows = FALSE,
                                cluster_cols = FALSE,
                                color = MAG_genus_colors,
                                breaks = MAG_genus_breaks,
                                show_rownames = TRUE,
                                fontsize = 24,
                                main = "VFs",
                                border_color = "grey",
                                legend = FALSE,
                                silent = TRUE,
                                display_numbers = MAG_genus_vf_mat_labels,
                                fontsize_number = 22,
                                number_color = "black",
                                annotation_col = MAG_genus_vf_ann,
                                annotation_colors = heatmap_annotation_colors,
                                annotation_names_col = FALSE)

grid.arrange(
    MAG_genus_arg_pheatmap[[4]],
    MAG_genus_vf_pheatmap[[4]],
    ncol = 2,
    widths = c(0.8, 1)
  )
```

```{r MAG-co-ARG-VF}
# Let's see which MAGs have contigs carrying ARGs and VFs
MAG_intersect <- intersect(unique(MAG_args_df$MAG), unique(MAG_vfs_df$MAG))

# Count the number of unique MAGs with both antibiotic resistance and virulence factors
MAG_co_args_vfs <- length(MAG_intersect)

# Extract the bin taxonomies for these intersecting bins
MAG_co_arg_vf_tax <- MAG_tax %>% 
  filter(MAG %in% MAG_intersect)

# Count the number belonging to pseudomonadota
MAG_co_pseudomonadota <- MAG_co_arg_vf_tax %>% 
  filter(Phylum == "Pseudomonadota") %>% 
  nrow()

# Count the number belonging to Bacillota
MAG_co_bacillota <- MAG_co_arg_vf_tax %>% 
  filter(Phylum == "Bacillota") %>% 
  nrow()
```

All `r MAG_num` MAGs were identified at least to the genus level, and `r MAG_species_num` were further identified to the species level. Most MAGs were assigned to the Pseudomonadota phylum (`r MAG_phylum$Count[MAG_phylum$Phylum == "Pseudomonadota"]`), followed by Actinobacteriota (`r MAG_phylum$Count[MAG_phylum$Phylum == "Actinobacteriota"]`), Bacillota (`r MAG_phylum$Count[MAG_phylum$Phylum == "Bacillota"]`), Bacteroidota (`r MAG_phylum$Count[MAG_phylum$Phylum == "Bacteroidota"]`), and `r num_to_word(MAG_phylum$Count[MAG_phylum$Phylum == "Bdellovibrionota"])` MAG was assigned to the phylum Bdellovibrionota  (**Supplementary figure 1**). The full MAG taxonomic classifications, alongside genome accessions, can be seen in (**`r get_supp("Interactive MAG taxonomic classification")`**).

Pseudomonadota phylum classified MAGs (**Figure \@ref(fig:MAG-genus-ARG-VF-plot)**), contributed `r MAG_phylum_sum$TotalCount[MAG_phylum_sum$Phylum == "Pseudomonadota"]` of the `r MAG_arg_class_counts` incidences of resistance conferred to `r MAG_phylum_sum$TotalClasses[MAG_phylum_sum$Phylum == "Pseudomonadota"]` of the `r MAG_arg_classes` classes of antibiotics from ARGs on contigs binned into MAGs. Within the Pseudomonadota phylum, the *Phytobacter* genus contributed `r MAG_genus_sum$TotalCount[MAG_genus_sum$Genus == "Phytobacter"]` incidences of resistance against `r MAG_genus_sum$TotalClasses[MAG_genus_sum$Genus == "Phytobacter"]` classes,the *Pseudomonas* genus contributed `r MAG_genus_sum$TotalCount[MAG_genus_sum$Genus == "Pseudomonas"]` incidences of resistance against `r MAG_genus_sum$TotalClasses[MAG_genus_sum$Genus == "Pseudomonas"]` classes, and the *Kosakonia* genus contributed `r MAG_genus_sum$TotalCount[MAG_genus_sum$Genus == "Kosakonia"]` incidences of resistance against `r MAG_genus_sum$TotalClasses[MAG_genus_sum$Genus == "Kosakonia"]` classes.

Pseudomonadota phylum classified MAGs (**Figure \@ref(fig:MAG-genus-ARG-VF-plot)**), contributed `r MAG_vf_phylum_sum$TotalCount[MAG_vf_phylum_sum$Phylum == "Pseudomonadota"]` of the `r MAG_vf_class_counts` total VFs across all `r num_to_word(MAG_vf_phylum_sum$TotalClasses[MAG_vf_phylum_sum$Phylum == "Pseudomonadota"])` of the VF product classes encoded by these genes on contigs binned into MAGs. Within the Pseudomonadota phylum, the *Pseudomonas* genus contributed `r MAG_vf_genus_sum$TotalCount[MAG_vf_genus_sum$Genus == "Pseudomonas"]` VFs across `r num_to_word(MAG_vf_genus_sum$TotalClasses[MAG_vf_genus_sum$Genus == "Pseudomonas"])` classes, and the *Ochrobactrum* genus contributed `r MAG_vf_genus_sum$TotalCount[MAG_vf_genus_sum$Genus == "Ochrobactrum"]` VFs across `r num_to_word(MAG_vf_genus_sum$TotalClasses[MAG_vf_genus_sum$Genus == "Ochrobactrum"])` classes. Of the `r MAG_num` MAGs, `r num_to_word(MAG_co_args_vfs)` were assembled from both ARG- and VF-encoding contigs and `r num_to_word(MAG_co_pseudomonadota)` of these ARG- and VF-encoding MAGs were classified to the Pseudomonadota phylum, with only `r num_to_word(MAG_co_bacillota)` to the Bacillota phylum.

```{r MAG-cooccurrence-network}
# Build a co-occurrence networks to link MAG taxa to ARGs, VFs, or both
# Take the MAG_args_df dataframe and remove unneccesary columns, and add a column to specify the gene as an ARG
MAG_args_genes <- MAG_args_df %>% 
  select(c(MAG, contig, ARG_gene)) %>% 
  rename(Gene = ARG_gene) %>% 
  mutate(Type = "ARG")

# Repeat for the VFs
MAG_vfs_genes <- MAG_vfs_df %>% 
  select(c(MAG, contig, VF_gene)) %>% 
  rename(Gene = VF_gene) %>% 
  mutate(Type = "VF")

# Bind together and merge the bin (MAG) genus and species taxonomies, then clean the species columns if there is no species level classification
MAG_genes <- rbind(MAG_args_genes, MAG_vfs_genes) %>% 
  left_join(MAG_tax %>% select(MAG, Genus, Species), by = "MAG") %>% 
  mutate(MAG = gsub("bin.", "MAG_", MAG)) %>% 
  mutate(Species = ifelse(
      is.na(Species) | trimws(Species) == "",
      paste0(Genus, "_sp."),
      Species
    )
  ) %>% 
  mutate(Species = gsub(" ", "_", Species)) %>% 
  mutate(Taxon = paste0(MAG, "_", Species))

# Build the edges and nodes (same as before) and now strip away the unique MAG identifiers
MAG_edges <- data.frame(from = MAG_genes$Taxon, to = MAG_genes$Gene)
MAG_nodes <- data.frame(name = unique(c(MAG_genes$Taxon, MAG_genes$Gene)))

# identify shared genes and attach an edge_color column to MAG_edges
shared_genes <- MAG_genes %>%
  group_by(Gene) %>%
  summarise(n_taxa = n_distinct(Taxon), .groups = "drop") %>%
  mutate(shared = n_taxa > 1)

# join the shared flag to edges (match by gene name)
MAG_edges <- MAG_edges %>%
  left_join(shared_genes %>% select(Gene, shared),
            by = c("to" = "Gene")) %>%
  mutate(shared = ifelse(is.na(shared), FALSE, shared),
         edge_color = ifelse(shared, "#D73027", "grey50"))

# --- Keep your MAG_nodes coloring logic (use your existing code for node colors) ---
# (example - adapt if you had this earlier)
MAG_nodes <- MAG_nodes %>%
  left_join(MAG_genes %>% select(Gene, Type) %>% distinct(),
            by = c("name" = "Gene")) %>%
  mutate(
    node_type = ifelse(name %in% MAG_genes$Taxon, "Taxon", "Gene"),
    color = case_when(
      node_type == "Taxon" ~ "grey20",
      Type == "ARG" ~ "#E66100",
      Type == "VF"  ~ "#1B9E77",
      TRUE ~ "grey70"
    )
  )

# --- Build the graph exactly as before but using the MAG_edges that already include edge_color ---
MAG_graph <- graph_from_data_frame(MAG_edges, vertices = MAG_nodes, directed = FALSE)

# Convert to tidygraph (no need to mutate edges inside tidygraph now)
MAG_graph_tbl <- as_tbl_graph(MAG_graph) %>%
  mutate(Type = ifelse(node_type == "Taxon", "Taxon", Type)) %>% 
  mutate(name = ifelse(node_type == "Taxon", gsub("^MAG_[0-9]+_", "", name), name),
         name = ifelse(node_type == "Taxon", gsub("_", " ", name), name))
```

```{r MAG-cooccurrence-network-plot, fig.cap="**ARG and VF encoding MAGs.** The ARG and VF genes encoded by contigs binned into taxonomically assigned MAGs. Unique taxa are represented by dark grey large nodes with ARG nodes in orange, and VF nodes in green. Grey links represent genes identified in the MAGs of a single taxon, red links represent genes identified in the MAGs of more than one taxon.", fig.width=7, fig.height=7, dpi=300}
# Set seed for reproducibility
set.seed(202507)
# Plot the MAG network
MAG_network_plot <- ggraph(MAG_graph_tbl, layout = "fr") +
  geom_edge_link(aes(color = shared), alpha = 0.8, width = 0.65) +
  scale_edge_color_manual(
    name = "Connection type",
    values = c(
      "TRUE" = "#D73027",
      "FALSE" = "grey50"
    ),
    labels = c(
      "TRUE" = "Shared between taxa",
      "FALSE" = "Single-taxon link"
    )
  ) +
  geom_node_point(aes(color = Type, 
                      size = ifelse(node_type == "Taxon", 5.8, 3)),
                  alpha = 0.7, na.rm = TRUE) +
  scale_color_manual(
    name = "Node type",
    values = c(
      "ARG" = "#E66100",
      "VF" = "#1B9E77",
      "Taxon" = "grey40"
    ),
    labels = c(
      "ARG" = "ARG gene",
      "VF" = "VF gene",
      "Taxon" = "MAG taxon"
    ),
    breaks = c("ARG", "VF", "Taxon")
  ) +
  geom_node_text(
    aes(label = name,
        fontface = ifelse(node_type == "Taxon", "bold", "plain"),
        size = ifelse(node_type == "Taxon", 6.5, 5)),
    color = "black",
    repel = TRUE,
    max.overlaps = Inf
  ) +
  scale_size_identity() +
  theme_void(base_family = "Aptos") +
  theme(
     legend.position = "bottom",
  legend.title = element_text(size = 44, face = "bold"),
  legend.text  = element_text(size = 32),
  legend.key.height = unit(0.6, "cm"),
  legend.key.width  = unit(1.2, "cm"),
  legend.box = "vertical",
  legend.box.just = "left",
  legend.spacing.y = unit(0.3, "cm")
  ) +
  guides(
    color = guide_legend(override.aes = list(size = 6)),
    edge_color = guide_legend(override.aes = list(width = 1.5))
  )

MAG_network_plot
```

```{r MAG-cooccurrence-sums}
# Grab the Pseudomonas species with many genes and count the total number of ARGs and VFs and then each individually
MAG_pseudomonas_gene_num <- MAG_genes %>% 
  filter(Genus == "Pseudomonas") %>% 
  nrow()

MAG_pseudomonas_arg_num <- MAG_genes %>% 
  filter(Genus == "Pseudomonas") %>% 
  filter(Type == "ARG") %>% 
  nrow()

MAG_pseudomonas_vf_num <- MAG_genes %>% 
  filter(Genus == "Pseudomonas") %>% 
  filter(Type == "VF") %>% 
  nrow()

# Number of shared genes from this Pseudomonas strain
MAG_pseudomonas_shared <- length(intersect(MAG_genes %>% filter(Genus == "Pseudomonas") %>% pull(Gene),
                                    MAG_genes %>% filter(Genus != "Pseudomonas") %>% pull(Gene))
                                 )

# Count Koskonia genes
MAG_kosakonia_gene_num <- MAG_genes %>% 
  filter(Genus == "Kosakonia") %>% 
  nrow()

# Count Phytobacter genes
MAG_phytobacter_gene_num <- MAG_genes %>% 
  filter(Genus == "Phytobacter") %>% 
  nrow()

# Count the shared the genes
MAG_koskonia_phytobacter_shared <- length(intersect(MAG_genes %>% filter(Genus == "Kosakonia") %>%  pull(Gene),
                                             MAG_genes %>% filter(Genus == "Phytobacter") %>% pull(Gene)))
```

The MAG of one *Pseudomonas* genus taxon consisted of contigs ncoding in total `r MAG_pseudomonas_vf_num` VFs and `r MAG_pseudomonas_arg_num` ARGs (**Figure \@ref(fig:MAG-cooccurrence-network-plot)**). Only `r num_to_word(MAG_pseudomonas_shared)` of these genes, both VFs, were shared with other taxa. The *Kosakonia sp000410515* and *Phytobacter massiliensis* taxa, also belonging to the Pseudomonadota phylum, each had `r MAG_kosakonia_gene_num` and `r MAG_phytobacter_gene_num` genes, respectively. Although, `r MAG_koskonia_phytobacter_shared` of these genes were shared. *Achromobacter xylosoxidans*, *Enterococcus gallinarum*, and a *Stenotrophomonas* species all had both ARG and VF sequences but shared none of their encoded genes with other MAGs. *Ochrobactrum teleogrylli*, a *Rhizobiaceae_63-22* species, and a *Shinella* species all encoded both ARG and VF sequences. An interactive version of this network can be seen in **`r get_supp("Interactive MAG network plot of ARG and VF genes")`**.

\newpage

## Isolation of antibiotic-resistant bacteria from the BSFL midgut

```{r isolate}
# Import the isolates
isolates <- read_excel("isolation/isolation.xlsx")

# Read the confidence scores
maldi_conf <- read.csv("isolation/MALDI_scores.csv", header = TRUE, fileEncoding = "UTF-8-BOM") %>% 
  group_by(ID) %>% 
  summarise(mean_score = mean (scores, na.rm = TRUE),
            n_iso = n()
            )

# Keep original and make a clean one without the diet samples, merge each section of the gut and merge the confidence scores
isolates_gut <- isolates %>% 
  filter(sample != "DIET") %>% 
  select(-c(colony, sample)) %>% 
  distinct() %>% 
  rename("conc" = "conc (µg/mL)") %>% 
  mutate(antibiotic = gsub("ampicillin", "Ampicillin", antibiotic),
         antibiotic = gsub("streptomycin", "Streptomycin", antibiotic),
         antibiotic = gsub("chloramphenicol", "Chloramphenicol", antibiotic),
         antibiotic = gsub("tetracycline", "Tetracycline", antibiotic)) %>% 
  mutate(Resistance = paste0(antibiotic, " (", conc, " µg/mL)")) %>% 
  select(-c(antibiotic, conc)) %>% 
  filter(ID != "Trichosporon asahii") %>% 
  left_join(maldi_conf, by = "ID") %>% 
  mutate(ID_conf = paste0(ID, "\n(Confidence = ", signif(mean_score, 3), "; n = ", n_iso, ")"))
  
# Make one for the diet sample alone also
isolates_diet <- isolates %>% 
  filter(sample == "DIET") %>% 
  select(-c(colony, sample)) %>% 
  distinct() %>% 
  rename("conc" = "conc (µg/mL)") %>% 
  mutate(antibiotic = gsub("ampicillin", "Ampicillin", antibiotic),
         antibiotic = gsub("streptomycin", "Streptomycin", antibiotic),
         antibiotic = gsub("chloramphenicol", "Chloramphenicol", antibiotic),
         antibiotic = gsub("tetracycline", "Tetracycline", antibiotic)) %>% 
  mutate(Resistance = paste0(antibiotic, " (", conc, " µg/mL)")) %>% 
  select(-c(antibiotic, conc)) %>%
  left_join(maldi_conf, by = "ID") %>% 
  mutate(ID_conf = paste0(ID, "\n(Confidence = ", signif(mean_score, 3), "; n = ", n_iso, ")"))

# Get the number of unique antibiotics tested
antibiotics <- length(unique(isolates_gut$Resistance))

# Get the number of unique taxa in the gut for the STD diet
STD_gut_resistant_taxa <- isolates_gut %>% 
  pull(ID) %>% 
  unique() %>% 
  length()

# Get the number of unique taxa in the rearing substrate
STD_diet_resistant_taxa <- isolates_diet %>% 
  pull(ID) %>% 
  unique() %>% 
  length()

# Find the isolates in the CAT contig classification
isolate_contigs <- contig_bacteria_tax %>% 
  filter(Species == "Alcaligenes faecalis" | Species == "Bordetella trematum" | Species == "Klebsiella pneumoniae" | Species == "Leucobacter holotrichiae" | Species == "Pseudomonas aeruginosa" | Species == "Morganella morganii" | Species == "Stenotrophomonas maltophilia" | Species == "Ochrobactrum intermedium" | Species == "Myroides injenensis")

# Get the unique spcies identified from contigs
isolate_contigs_unique <- unique(isolate_contigs$Species)

# Look at the genera for those not identiied to species
isolate_contigs_genera <- contig_bacteria_tax %>% 
  filter(Genus == "Leucobacter" | Genus == "Ochrobactrum" | Genus == "Myroides")
```

```{r isolate-alluvial, fig.cap="**Resistant bacterial isolates of the BSFL bacterial midgut community.** (**A**) Diagram of resistant midgut bacteria isolation. Midgut lumen content (**B**) or rearing diet during larval growth (**C**) was prepared and plated onto LB-agar plates with different antibiotics and incubated at 37 °C for 16 hours. Classification at species level was obtained using MALDI-TOF ID. The average log score of spectral similarity and the number of identifications were presented alongside identifications.", fig.width=6, fig.height=7.5, dpi=300}
# Import the graphical abstract as a ggplot object
isolate_graphic <- wrap_elements(
  grid::rasterGrob(png::readPNG("plots/isolation_graphic.png"), interpolate = TRUE)
)

# Alluvial plot of gut isolation
isolate_alluvial_gut <- ggplot(isolates_gut,
                            aes(axis1 = ID_conf, axis2 = Resistance)) + 
  geom_alluvium(aes(fill = ID), width = 1/3) +
  geom_stratum(width = 1/3, fill = "lightgrey", color = "black", size = 0.1) +
  geom_text(stat = "stratum", aes(label = after_stat(stratum)), size = 7, fontface = "bold", lineheight = 0.3) +
  scale_x_discrete(limits = c("ID_conf", "Resistance"), expand = c(0.05, 0.05)) +
  scale_y_continuous(expand = c(0, 0)) +
  theme_minimal() +
  theme(axis.title = element_blank(),
        axis.text = element_blank(),
        axis.ticks = element_blank(),
        panel.grid = element_blank(),
        legend.position = "none") +
  scale_fill_viridis_d()

# Alluvial plot of diet isolation
isolate_alluvial_diet <- ggplot(isolates_diet,
                            aes(axis1 = ID_conf, axis2 = Resistance)) + 
  geom_alluvium(aes(fill = ID), width = 1/3) +
  geom_stratum(width = 1/3, fill = "lightgrey", color = "black", size = 0.1) +
  geom_text(stat = "stratum", aes(label = after_stat(stratum)), size = 7, fontface = "bold", lineheight = 0.3) +
  scale_x_discrete(limits = c("ID_conf", "Resistance"), expand = c(0.05, 0.05)) +
  scale_y_continuous(expand = c(0, 0)) +
  theme_minimal() +
  theme(axis.title = element_blank(),
        axis.text = element_blank(),
        axis.ticks = element_blank(),
        panel.grid = element_blank(),
        legend.position = "none") +
  scale_fill_viridis_d()

isolate_alluvial <- isolate_graphic /
  isolate_alluvial_gut /
  isolate_alluvial_diet +
  plot_layout(heights = c(0.6, 1,0.4)) +
  plot_annotation(tag_levels = "A") &
    theme(
      plot.tag = element_text(size = 80)
      )

isolate_alluvial
```

In total, `r num_to_word(STD_gut_resistant_taxa)` taxa were identified from plating BSFL midgut lumen contents onto `r num_to_word(antibiotics)` different antibiotic agar plates (**Figure \@ref(fig:isolate-alluvial)A**).  *Alcaligenes faecalis*, *Myroides injenensis*, and *Pseudomonas aeruginosa* species were all identified on more than one type of antibiotic-containing plate. *Bordetella trematum*, *Klebsiella pneumoniae*, *Leucobacter holotrichiae*, *Morganella morganii*, *Ochrobactrum intermedium*, and *Stenotrophomonas maltophilia* species were all identified on a single type of antibiotic plate investigated here. *A. faecalis*, *M. injenensis*, and *P. aeruginosa* species were all identified additionally in the rearing diet during larval growth (**Figure \@ref(fig:isolate-alluvial)B**).

For the *M. morganii* isolate a potentially corresponding MAG was reconstructed (**Figure \@ref(fig:MAG-genus-ARG-VF-plot)**). Contigs were identified classified to *B. trematum*, *S. maltophilia*, *K. pneumoniae*, *A. faecalis*, *M. morganii*, and *P. aeruginosa* species according to CAT taxonomic classification. Although contigs classified to *O. intermedium*, *M. injenensis*, and *L. holotrichiae* species were not identified, contigs classified to these genera were identified.

\newpage

# Discussion

Through the metagenomic analysis of the BSFL midgut microbiota performed in this study, it was possible to detect resistance genes for over one third of the 64 unique antibiotic classes reported in CARD, the most complete peer-reviewed database on antibiotic resistance [@alcock2023]. The most abundant form of antibiotic resistance encoded in the bacterial community was mediated by efflux pumps. This was unsurprising due to the prevalence of this resistance strategy employed by a wide range of bacteria, and due to the lack of chemical or structural specificity with which efflux pumps export diverse antibiotics [@gaurav2023]. Indeed, several of these efflux pumps might exert their activity against antibiotics as secondary targets and be involved in other functions. In the BSFL midgut microbiome, incidences of resistance to the fluoroquinolone class of antibiotics were the most common, reflecting the long-term use of these antibiotics beginning in the early 1960s [@kherroubi2024]. Penam, including penicillins, and cephalosporin class antibiotics were the next most abundant resistance classes. Together these antibiotics constitute 55 % of antibiotic consumption [@wang2025a]. Over 50 incidences of resistance were conferred to the tetracycline class, again a reflection of their long-term use in the past 60 years, and cause concern due to their wide target activity against both Gram-positive and -negative pathogens [@grossman2016]. The glycopeptide class of antibiotics, considered as "last resort" antibiotics [@selim2022], had 14 incidences of resistance. Only for 3 of the 26 classes a single incidence of resistance was observed. Overall, bacterial contigs predicted to encode sequences for ARGs were found to contribute only slightly to the community structure. The low relative abundances of ARGs identified in this study was similar to the low abundances of ARGs observed previously in BSFL [@rong2024; @schokker2025], This is unsurprising, as in the absence of selection ARGs could represent a metabolic burden for bacteria [@martinez2011]. Similarly low numbers of ARGs have also been found in other environmental ARG reservoirs [@zhu2025; @wang2020a], which still represents a serious concern.

A comprehensive profile of VFs encoded in the BSFL midgut microbiota was documented here. This not only provided an understanding of the potential pathogenicity associated with the midgut bacterial community, but may also be useful in combating antibiotic resistance, as it has been suggested that resistant pathogens could be disarmed through suppression of their virulence genes [@zhou2024]. Investigating virulence identified `r CAT_bacteria_vf_count` VF genes, with motility as the most abundant class of VF genes identified in the microbiota likely due to the large numbers of genes required to construct the highly complex structures enabling bacterial movement [@kao2014]. The second most abundant VF gene class in the midgut community was the biofilm formation class. Although biofilm formation is not directly involved in causing disease, it may allow pathogenic or resistant bacteria to persist in the environment, thus increasing the risk of ARG spread [@niu2013; @bai2021]. Other VFs more directly related to pathogenicity such as effector delivery, exotoxins, and exoenzymes, were also identified in the BSFL midgut microbiome. Taxa harboring these genes alongside those for antibiotic resistance could represent a significant public health risk, and as discussed before, the dissemination of DNA encoding these traits could also be dangerous [@lichev2019]. Similarly to the predicted ARGs, the Pseudomonadota phylum was the dominant taxonomic reservoir Pseudomonadota is indeed a diverse phylum containing a range of pathogenic and antibiotic-resistant families [@leao2023]. Furthermore, the dominance of this phylum in the ARG-encoding community of the BSFL midgut microbiota has been highlighted previously [@rong2024; @schokker2025].

Several contigs containing typical plasmid sequences were identified in the BSFL midgut community and in particular `r num_to_word(arg_total_mobile)` mobile plasmids encoded antibiotic resistance. Plasmid JN420336 was predicted to encode resistance against the penam class of antibiotics and the widely used cephalosporins [@wang2025a], while also encoding a VF for adherence and therefore with the potential to form films that increase the likelihood of the transfer of this conjugatively mobile plasmid [@dadehamirfard2024; @stalder2016]. Plasmids JN420336 and JX514065, known vehicles for carbapenemase dissemination [@villa2012; @dolejska2013], were also reported here. The rise of carbapenemase resistant members of Enterobacteriaceae family is considered a serious concern and are therefore highly relevant for public health [@tuhamize2024]. As discussed earlier however, plasmids are not the sole distributors of genes horizontally. Although the risk of horizontal gene transfer is hard to assess for genomic DNA which may become extracellular [@bender2022], transformation with extracellular DNA has contributed to the spread of both resistance and virulence [@lichev2019]. Indeed, in combination with the capacity for biofilm formation in the BSFL midgut bacterial community, the likelihood of cell-cell contacts and therefore the spread of these ARG-encoding plasmids between community members and the environmental microbiome may even be increased [@dadehamirfard2024; @stalder2016].

Members of the BSFL midgut bacterial community with tools to cause disease and resist treatment already pose a risk if introduced into food systems. MAG reconstruction revealed a *Pseudomonas* taxon which encoded `r num_to_word(MAG_pseudomonas_arg_num)` ARGs providing resistance theoretically to `r MAG_genus_sum$TotalClasses[MAG_genus_sum$Genus == "Pseudomonas"]` classes of antibiotics, alongside `r MAG_pseudomonas_vf_num` VFs mainly of the motility class but also of the effector delivery, nutritional modification, biofilm development, exoenzyme production, adherence, and modulation of transcriptional regulation classes of VFs. The disease-causing abilities of many *Pseudomonas* species are well documented, lying within the top six leading causes (73 %) of AMR-associated deaths in 2019 [@girard2021; @ho2025], but this was not the only taxa identified encoding both ARG and VF genes. *Kosakonia sp000410515* from the occasionally human-infecting genus *Kosakonia* [@merlino2025; @chen2023], and its phylogenetic neighbour *Phytobacter massiliensis* share `r MAG_koskonia_phytobacter_shared` ARG and VF genes [@ma2020]. *Phytobacter massiliensis* was part of a cluster of shared genes comprising other Enterobacteraceae species (i.e., *Morganella morganii*, *Providencia rettgeri* and *Providencia stuartii*). Some of the ARGs found in *Phytobacter masiliensis* encoded resistance against carbapenem and "last-resort" antibiotics such as colistin. The spread of carbapenem resistant genes in the Enterobacteriaceae family is of high relevance for public health, as these antibiotics are the last-line of defence for many serious infections [@tuhamize2024]. These results could suggest that lesser known members of the Enterobacteriaceae family such as Phytobacter can be important reservoirs of ARGs and their spread should be monitored, especially in new and emerging contexts like insect rearing facilities. *Ochrobactrum teleogrylli*, *Achromobacter xylosoxidans*, *Enterococcus gallinarum*, a species of the *Rhizobiaceae* genus, a species of the *Shinella* genus, and a species of the *Stenotrophomonas* genus all encoded both ARG and VF genes. Members of some of these taxa were already reported in clinical samples or healthcare-associated infections [@isler2020; @monticelli2018; @ryan2009; @ryan2020], highlighting the need for careful monitoring of the microbiological safety of BSFL derivatives used in agriculture and food system processes.

To validate the bioinformatic analyses of metagenomic data, antibiotic-resistant bacterial species were isolated from the BSFL midgut and from the rearing diet using a culture-based approach with a selection of antibiotics. Nine unique bacterial isolates were identified at species level through MALDI TOF across the eight antibiotics tested. Contigs belonging to all nine genera were identified in the shotgun metagenomic data. Three resistant species, i.e., *A. faecalis*, *M. injenensis*, and *P. aeruginosa* were also found in the frass. The transfer of genes through HGT has been reported for both *A. faecalis* [@lang2022] and *P. aeruginosa* [@karampatakis2025].

Overall, the results suggested that the microbiome of BSFL comprises several potentially pathogenic bacterial species carrying ARGs and VFs in their genomes, with microbiological evidence of the spread of some of these taxa to the rearing diet. This calls attention to the safety of workers in BSF facilities but should also demand the careful monitoring of BSFL-derived products. Moreover, despite the reduction of ARGs observed in some studies on diets with high prevalences of antimicrobial resistant bacteria [@zhao2023; @zhao2023a], these genes are present during and at the end of the bioconversion practices. As suggested by the same authors, it will be important to establish downstream processes to reduce the risk of their spread in the environment, preventing the uptake of ARGs by the environmental microbiome or their introduction in the food chain.

BSFL remains an important biotechnological tool for the development of innovative processes that can offer innovation to agricultural and industrial processes. However, the associated risks should encourage the adoption of procedures to mitigate ARG and VF dissemination into the environmental microbiome. Treatment strategies similar to those applied to manure to reduce ARG abundance could be adopted for BSFL frass [@zalewska2023], or phage-based approaches could be used to target specific high-risk bacterial taxa in BSFL-derived products [@liao2025]. Alongside such innovation, careful monitoring will be essential to support the safe and effective use of BSFL biomass and frass. By providing a complete profile of the ARGs, VFs, and plasmids encoded within the BSFL midgut bacterial community, this study has established a valuable resource for this organism. In addition, it emphasizes the urgent need to carefully assess the agricultural applications of BSFL-derived products. Overall, this study represents an important contribution to the BSFL research community, improving our understanding of the complex microbiome, and the potential health risks associated with its dissemination into food systems and the environment.

\newpage

# References

::: {#refs}
:::

\newpage

# Supplementary material

## Supplementary files

1.  [**Shotgun metagenomic read quality report**](#sf1)**.** MultiQC report of fastqc results of quality control checks for shotgun metagenomic reads files after trimming and filtering with PRINSEQ 0.20.4 for Phred score $\geq$ 20 and read length $\geq$ 75 bp.

2. [**Interactive taxonomic community structures**](#sf2)**.** Interactive document with multiple interactive figures allowing all taxonomic levels from kingdom to species to be explored for the CAT and Kraken2 predicted the BSFL midgut microbiota bacterial community.

3. [**Community, ARG, VF, and Plasmid database**](#sf3)**.** Databases of the entire community of the BSFL midgut containing all raw read depth values and normalised abundance values alongside CAT and kraken2 taxonomic classification raw outputs for all contigs of the coassembly, with predicted ARGs, VFs, and plasmids from the CARD, VFDB, and PlasmidFinder databases, respectively. Additionally, separate databases for ARGs, VFs, and plasmids for contigs containing these genes only.

4. [**MAG taxonomic classification**](#sf4)**.** The GTDB-Tk taxonomic classification of the MAGs of the BSFL midgut microbiota provided as a database of the raw output and also in an interactive hierarchical sunburst plot alongside genus and species classifications with matching reference genome accessions identified with FastANI or pplacer approaches.

5. [**Interactive MAG network plot of ARG and VF genes**](#sf5)**.** Interactive network plot depicting the connections between taxon nodes and encoded ARGs and VFs within MAGs of the BSFL midgut microbiota.

6. [**R Markdown manuscript file**](#sf6)**.** The R markdown file used to generate this manuscript containing all text and the code for analysis and plotting.

\newpage

## Supplementary tables

```{r bacteria-taxon-table}
# Make and print the table of otus assigned to each taxonomic level and the unique classifications within each taxon level
contig_taxon_tab <- contig_taxon_sum %>% 
  flextable(, defaults = TRUE) %>% 
      set_caption(caption = "Taxonomic assignments of bacterial contigs in the BSFL midgut community. The number of contigs assigned at each taxonomic level and the number of unique classifications within each taxonomic level for contigs assigned to the bacterial superkingdom.",
                  style = "Table caption") %>% 
        width(width = c(1.7,1.5,2)) %>% 
          align(align = c("left", "right", "right"), part = "all") %>% 
  style_flextable()
contig_taxon_tab
```

<br>

```{r sup-bacteria-phylum-table}
# Make and print the table of contig counts and percentages to each level of phylum
bacteria_phylum_tab <- bacteria_phylum %>% 
  mutate(across(where(is.numeric), ~ signif(.x, 3))) %>% 
  flextable(, defaults = TRUE) %>% 
      set_caption(caption = "Taxonomic assignments to phyla for bacterial contigs in the BSFL midgut community. The counts and percentages of bacterial contigs assigned to eachtaxonomic phyla.",
                  style = "Table caption") %>% 
        width(width = c(1.5,1,1)) %>% 
          align(align = c("left", "right", "right"), part = "all") %>% 
  style_flextable()
bacteria_phylum_tab
```

<br>

```{r antibiotic-class-count-tab}
antibiotic_class_count_tab <- antibiotic_class_count %>% 
  flextable(, defaults = TRUE) %>% 
      set_caption(caption = "Encoded resistance to antibiotic classes in the BSFL midgut bacterial community. The number of incidences of resistance encoded against antibiotic classes by ARGs identified on bacterial contigs in the BSFL midgut community.",
                  style = "Table caption") %>% 
        width(width = c(1.5,0.5,1)) %>% 
          align(align = c("left", "left", "right"), part = "all") %>% 
  style_flextable()
antibiotic_class_count_tab
```

<br>

```{r plasmid-ARG-tab}
# Table of plasmids with ARGs
co_plasmid_args_tab <- co_plasmid_args_df %>% 
  flextable(, defaults = TRUE) %>% 
      set_caption(caption = "Plasmid gene- and ARG-encoding contigs in the BSFL midgut community. Contigs of the coassembly predicted to encode both plasmid gene sequences and ARGs with identified plasmid accessions, ARG gene names, and CARD product description for the predicted ARG products.",
                  style = "Table caption") %>% 
        width(width = c(1,1,1,3)) %>% 
          align(align = c("left", "left", "left", "left"), part = "all") %>% 
  style_flextable()
co_plasmid_args_tab
```

<br>

```{r plasmid-VF-tab}
# Table of plasmids with ARGs
plasmid_vf_mobility_tab <- plasmid_vf_mobility %>% 
  flextable(, defaults = TRUE) %>% 
      set_caption(caption = "VF-encoding plasmids of the BSFL midgut community. The accessions of plasmids identified in the coassembly with encoding VF sequences alongside the plasmid predicted mobility, predicted VF ID, the VF gene name, the VF category, and the product description.",
                  style = "Table caption") %>% 
        width(width = c(0.8,1,0.6,0.7,1,2)) %>% 
          align(align = c("left", "left", "left", "left", "left", "left"), part = "all") %>% 
  style_flextable()
plasmid_vf_mobility_tab
```

<br>

\newpage

## Supplementary figures

```{r MAG-phylum-donut, fig.width=7, fig.height=5, dpi=300}
MAG_phylum_donut <- ggplot(MAG_phylum, aes(ymax = ymax, ymin = ymin, xmax = 4, xmin = 3, fill = Phylum)) +
  geom_rect(alpha = 0.6) +
  geom_text(x = 3.5, aes(y = label_position, label = label), size = 16) +
  scale_fill_manual(values = my_wes_palette(length(MAG_phylum$Phylum))) +
  coord_polar(theta = "y") +
  xlim(c(1.5,4)) +
  theme_void(base_family = "Aptos") +
  theme(
    legend.position = "right",
    legend.title = element_text(size = 50, color = "black"),
    legend.text = element_text(size = 44, color = "black"))

(MAG_phylum_donut)
```

Supplementary figure 1. **Taxonomic assignments to phyla of MAGs in the BSFL midgut community.** The number of MAGs taxonomically assigned to each bacterial phyla through GTDB-Tk taxonomic classification.
````
